# Comparative genomics of two closely related coral species with different spawning seasons reveals genomic regions possibly associated with gametogenesis

**DOI:** 10.1101/2023.08.03.551769

**Authors:** Shiho Takahashi-Kariyazono, Akira Iguchi, Yohey Terai

**Affiliations:** Geological Survey of Japan, National Institute of Advanced Industrial Science and Technology (AIST), Tsukuba, Ibaraki, Japan; SOKENDAI (The Graduate University for Advanced Studies), Research Center for Integrative Evolutionary Science, Hayama, Kanagawa, Japan; Research Laboratory on Environmentally-Conscious Developments and Technologies [E-code], National Institute of Advanced Industrial Science and Technology, Ibaraki 305-8567, Japan

**Keywords:** mTORC1, oogenesis, cnidaria

## Abstract

Marine invertebrates release their gametes at an optimal time to produce the next generation. In reef-building scleractinian corals, synchronous spawning is essential for reproductive success. Molecular mechanisms of scleractinian gametogenesis have been studied; however, the mechanism by which coral gametes mature at specific times has yet to be discovered. The present study focused on two *Acropora* species with different spawning seasons. In Okinawa, Japan, *Acropora digitifera* spawns from May to June, whereas *Acropora* sp. 1 spawns in August. Comparative genomic analyses revealed that 39 candidate genes are differentiated between the two species, suggesting a possible association with timing of gametogenesis. Among candidate genes, we identified an *Acropora* sp. 1-specific amino acid change in gene *WDR59*, one of the components of a mTORC1 activator, GATOR2. Since regulation of gametogenesis by mTORC1 is widely conserved among eukaryotes, the difference in timing of gamete maturation observed in the two *Acropora* species may be caused by a substitution in WDR59 that slightly affects timing of mTORC1 activation via GATOR2. In addition, this substitution may lead to reproductive isolation between the two species, due to different spawning periods. Thus, we propose that A. digitifera and Acropora sp. 1 species pair is an effective model for studying coral speciation and understanding the molecular mechanisms that control coral spawning timing.

**Significance statement (required):** For successful coral reproduction, conspecific corals must spawn synchronously. Gamete production initiates coral spawning. Regulation of gamete maturation by a protein complex, mTORC1, is widely conserved among organisms, but little is known about it in cnidarians. In this study, we analyzed genomes of two closely related *Acropora* species with different spawning months, May/June and August. Our analyses revealed that 39 genes are genetically differentiated between the two species. One of these is a component of mTORC1 activator, suggesting that this gene may be associated with the difference in spawning times of these two species.

## Introduction

In marine invertebrate reproduction, gametes are released into the water to be fertilized externally (spawning) (Mercier and Hamel 2010). Spawning occurs at an optimal time to produce the next generation (Forrest and Miller-Rushing 2010). Since fertilization in seawater can easily fail due to sperm dilution and other factors, marine organisms have evolved mechanisms such as synchronized spawning (Fukami, et al. 2003; Levitan, et al. 2004).

Spawning in reef-building, scleractinian corals is one of the most massive reproductive events on earth. In the Great Barrier Reef, most corals release their gametes once a year for a few nights (Harrison, et al. 1984). For example, over 100 coral species spawn in the Great Barrier Reef between the full and last quarter moon in late spring (Babcock, et al. 1986). Synchronous spawning within species is essential for fertilization because dilution and aging of sperm reduce fertilization success (Fukami, et al. 2003; Levitan, et al. 2004). In synchronous spawning, gametes spawned by different species are present in the water and may encounter each other. However, many *Acropora* species exhibit species specificity in gamete compatibility (Hatta, et al. 1999; Willis, et al. 1997), and interspecific hybridization rarely occurs in the Indo-Pacific (Hatta and Matsushima 2008; Isomura, et al. 2013).

Environmental cues act on corals to regulate spawning months, days, and times (Babcock, et al. 1986; Baird, et al. 2009). Temperature strongly influences gamete maturation (Baird, et al. 2009), and in several coral species, spawning has become asynchronous, due to effects of recent climate change (Shlesinger and Loya 2019). Therefore, understanding mechanisms of synchronous gamete maturation will help us estimate the impact of climate change on coral reproduction and restoration using coral seedlings produced from gametes (Suzuki, et al. 2020). Gametogenesis in corals has been studied in the field (Harrison 2011) and by molecular biological approaches (Chiu, et al. 2020; Shikina and Chang 2016). However, the mechanism by which coral gametes mature at specific times has yet to be identified.

In the Indo-Pacific region, including Okinawa, Japan, the genus *Acropora* comprises the largest number of coral species (Veron 2000). In Okinawa, most *Acropora* species spawn around the full moon in May or June, with a few species spawning several months later (Hayashibara, et al. 1993). One species that spawns later is *Acropora* sp. 1. This species was initially classified as *Acropora digitifera* (Wallace 1999); however, the two are now recognized as separate species, due to differences in morphology and spawning time (Hayashibara and Shimoike 2002; Nakajima, et al. 2012; Ohki, et al. 2015). *Acropora* sp. 1 has a flatter colony shape and shorter branches than *A. digitifera* (Hayashibara and Shimoike 2002; Ohki, et al. 2015). *Acropora* sp. 1 tends to inhabit reef edges with faster (offshore) currents than *A. digitifera*. In addition, in Okinawa, *A. digitifera* spawns from May to June, whereas *Acropora* sp. 1 spawns in August (Hayashibara and Shimoike 2002). Gametes of both species can cross-fertilize as indicated by artificial fertilization experiments (Ohki, et al. 2015). Under natural conditions, however, the two species do not interbreed because of the different spawning months (Ohki, et al. 2015).

Advances in analysis of genomic data with next-generation sequencers have revealed the genetic basis of specific traits (Ellegren and Sheldon 2008). In particular, comparative genomic analyses between genetically close species have identified genomic regions associated with their phenotypic differences (Poelstra, et al. 2014; Turner, et al. 2005). So far, genomes of various corals have been sequenced (Fuller, et al. 2020; Shinzato, et al. 2021; Shinzato, et al. 2011; Voolstra, et al. 2015), and population genomic approaches have identified loci associated with heat tolerance (Smith, et al. 2022). Comparative genomic analysis has yet to be conducted to identify genomic regions associated with differences in coral spawning timing due to the lack of closely related species pairs to compare.

In this study, we performed a comparative genomic analysis between *A. digitifera* and *Acropora* sp. 1 to identify genomic regions likely involved in trait differences between them. We expected that *A. digitifera* and *Acropora* sp. 1 were genetically closely related based on analysis of short sequences (Nakajima, et al. 2012) and their fertilization ability (Ohki, et al. 2015). Therefore, we determined the genome sequences of both species. This comparative genomic analysis identified genomic regions likely associated with differences in their spawning times. Since differences in spawning time can lead to reproductive isolation, these species will be a useful model to study coral speciation and to understand molecular mechanisms that regulate spawning time in corals.

## Results

### The spawning month of Acropora sp. 1

We collected 16 *Acropora* sp. 1 colonies during 2018-2020 at Sesoko and Bise, Okinawa, Japan (Fig.1), and observed mature oocytes or spawning in August (Table S1). This later-spawning month of *Acropora* sp. 1 is consistent with previous observations (Hayashibara and Shimoike 2002; Nakajima, et al. 2012; Ohki, et al. 2015).

**Figure 1.**
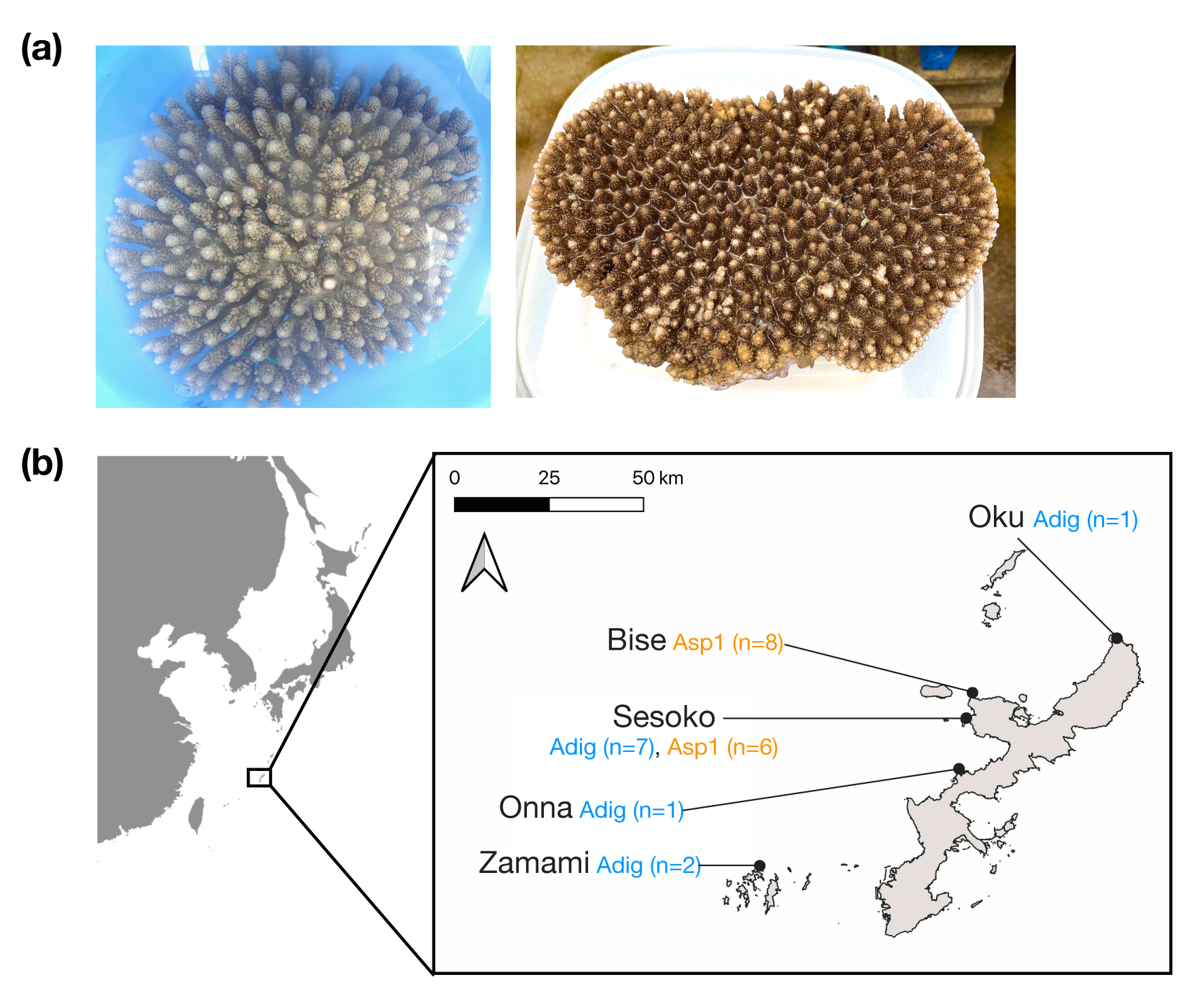
(a) Adult colonies of *Acropora digitifera* (left) and *Acropora* sp. 1 (right). (b) Sampling locations are shown as dots on the map of Okinawa Island.

### Phylogenetic tree and principal component analysis

We sequenced genomes from 16 colonies of *Acropora* sp. 1 (Table S1). Then, we mapped these reads to the *A. digitifera* whole-genome assembly ver. 2.0 (Shinzato, et al. 2021) and selected 14 colonies with coverage >10x for analyses. In addition, we downloaded genomic sequence data for 11 colonies of *A. digitifera* and one colony each of 15 other *Acropora* species from the DNA Data Bank of Japan (DDBJ) and mapped them as well.

First, we investigated the genetic relationship between *A. digitifera* and *Acropora* sp. 1. We extracted 885,405 biallelic single-nucleotide polymorphisms (SNPs) from mapping data of 17 species using our criteria (Materials and Methods). With these SNPs, we constructed a phylogenetic tree (Figure 2a). *Acropora digitifera* and *Acropora* sp. 1 colonies formed a monophyletic clade. In this clade, *A. digitifera* and *Acropora* sp. 1 colonies each formed monophyletic clades. A monophyletic *A. digitifera*/*Acropora* sp. 1 clade formed a monophyletic clade with *A. acuminata, A. microphthalma,* and *A. nasuta* (Fig. 2a). Using five species in this monophyletic clade, we performed principal component analysis (PCA). We used 80,490 SNPs extracted from *A. digitifera*, *Acropora* sp. 1, and three out-group species (*A. acuminata, A. microphthalma*, and *A. nasuta*). The three out-group species were separated along the PC1 axis from *A. digitifera* and *Acropora*. sp. 1 colonies, forming distinct genetic clusters. *Acropora digitifera* colonies were separated from *Acropora* sp. 1 along the PC2 axis (Fig. 2b). Among *Acropora* sp. 1 colonies, two (Colony IDs: Asp1B1906 and Asp1B1904) were separated from other *Acropora* sp. 1 colonies by PCA. In addition, these two colonies (Colony IDs: Asp1B1906 and Asp1B1904) formed a single clade with high bootstrap support in the phylogenetic tree (Fig. 2a). These two colonies were sampled from the same sites as other colonies sampled in the same year, indicating no geographic isolation.

**Figure 2.**
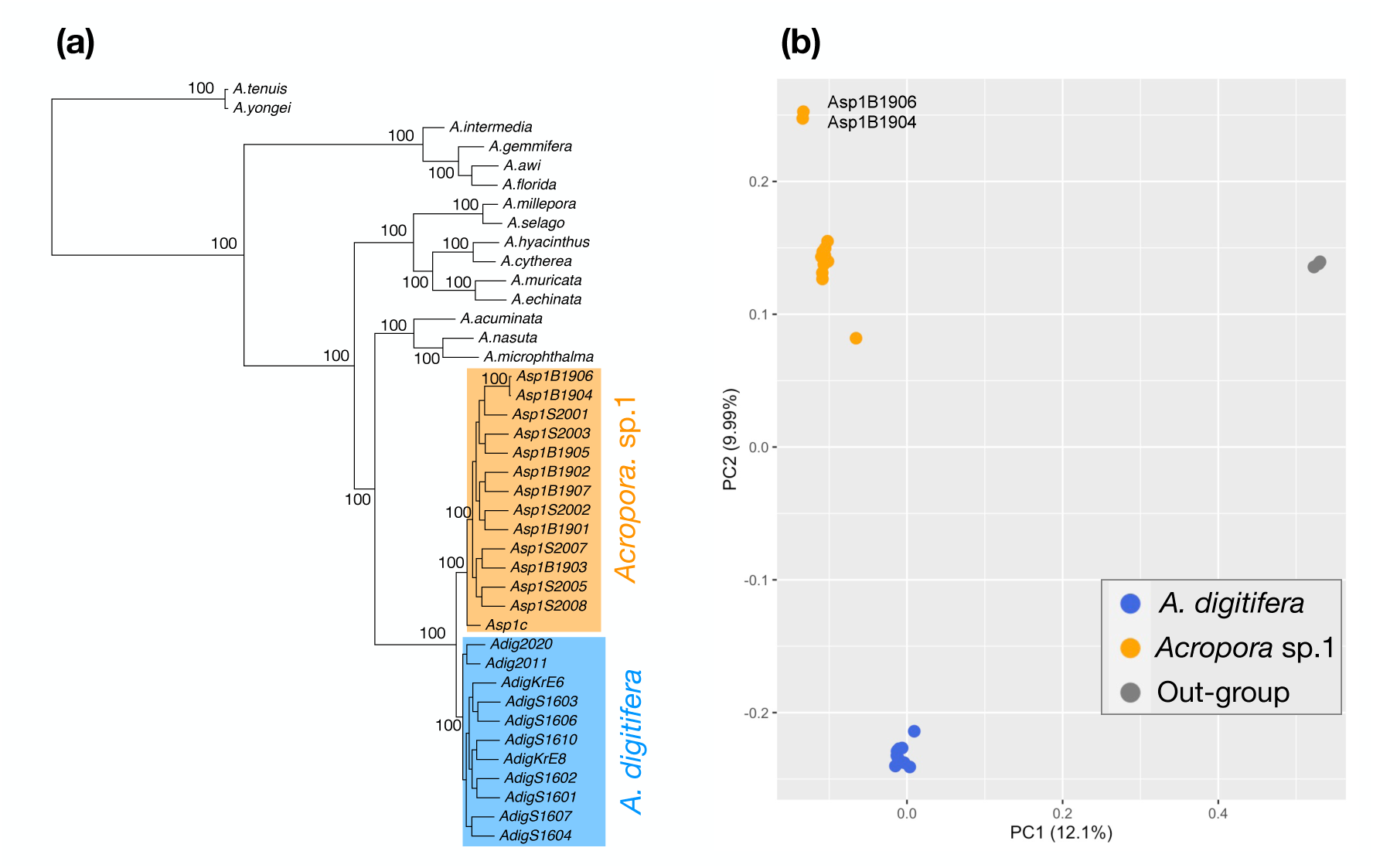
Phylogenetic relationship of *Acropora* sp. 1 (a) Phylogenetic relationships of 17 *Acropora* corals were analyzed based on 885,405 SNPs using the maximum likelihood method with the GTR option. Bootstrap support, shown next to each node for each clade, was obtained from 1,000 replicates. (b) PC1 and PC2 were derived from PCA based on SNPs for all individuals of *A. digitifera*, *Acropora* sp. 1, and three *Acropora* species as an out-group.

### Highly differentiated regions between A. digitifera and Acropora sp. 1

Since phylogenetic analysis indicated that *A. digitifera* and *Acropora* sp. 1 are closely related, the degree of differentiation between the two species was calculated (F_ST_) (Weir and Cockerham 1984) using 1,459,328 SNPs. The F_ST_ (Weir and Cockerham 1984) value across the genomes of these two species was 0.10225. This is comparable to the genetic divergence of species pairs used in comparative genome analysis in previous studies (Ellegren, et al. 2012; Geraldes, et al. 2011; Nadeau, et al. 2013). Despite low differentiation throughout their genomes, genomic regions responsible for differences in traits between *A. digitifera* and *Acropora* sp. 1 are expected to differ in the two species. To extract differentiated regions, we performed a sliding window analysis of 10 kb in 1 kb increments between *A. digitifera* and *Acropora* sp. 1. Genomic regions with the top 0.1% F_ST_ (Hudson, et al. 1992) values (F_ST_ >0.6157) in each 10 kb window were then selected. We further selected windows containing differentiated SNPs (Materials and Methods) from the top 0.1% F_ST_ (Hudson, et al. 1992) windows. When these windows overlapped, they were combined. As a result, 34 genomic regions, called highly differentiated regions (HDRs) (Table S2), were extracted from the whole genome.

### Genes in highly differentiated regions

In the HDRs, 39 genes harbor differentiated SNPs. We performed a Blast search using these 39 genes as queries (Figure S1) and found that 23 of them are similar to high-quality manually annotated genes (Table 1). Ten genes are similar to genes with automated annotations related to known genes (Table S3). Four genes are similar to uncharacterized genes (Table S3), and two genes have no similarity to any others in the NCBI nucleotide database.

**Table 1.**
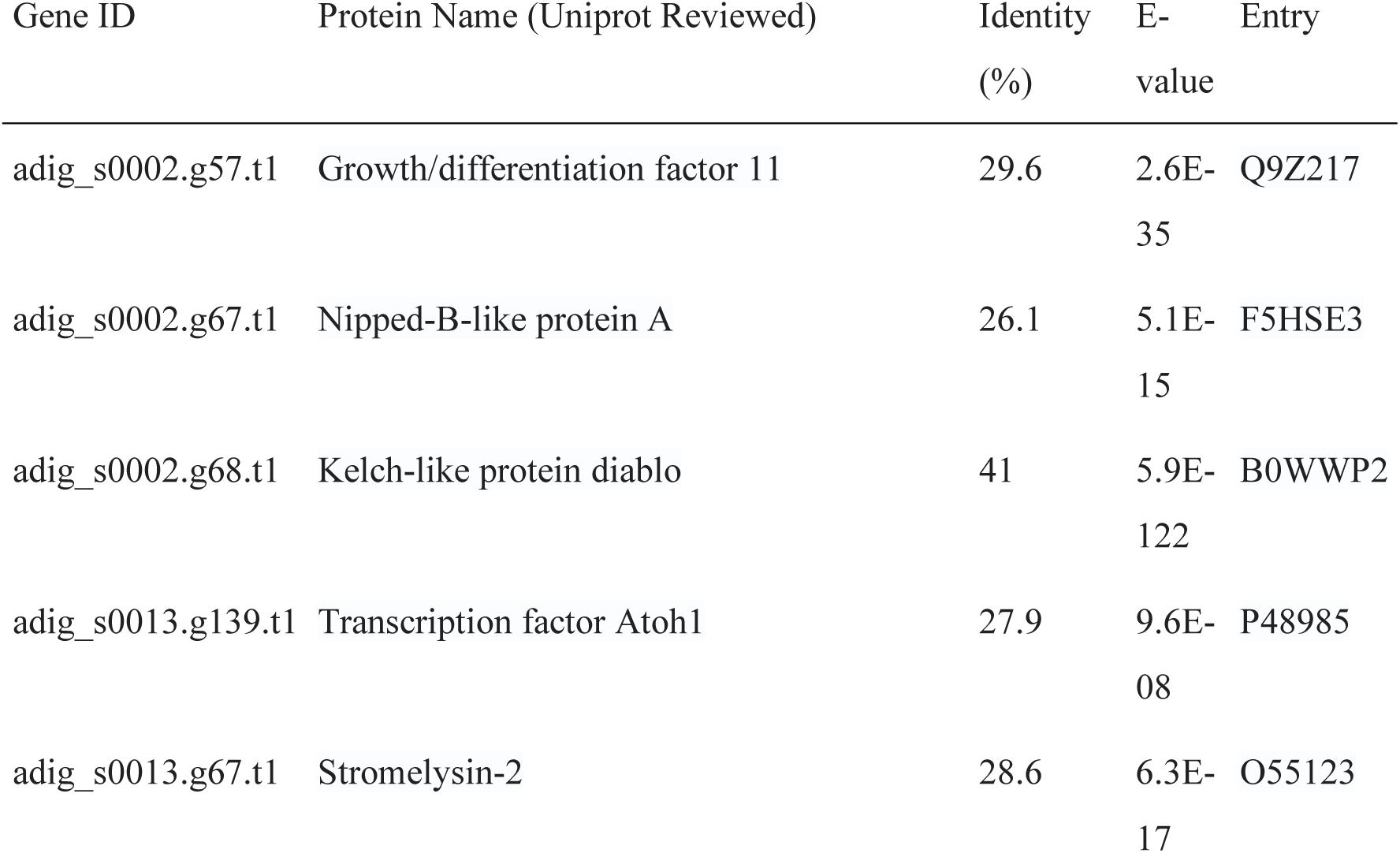
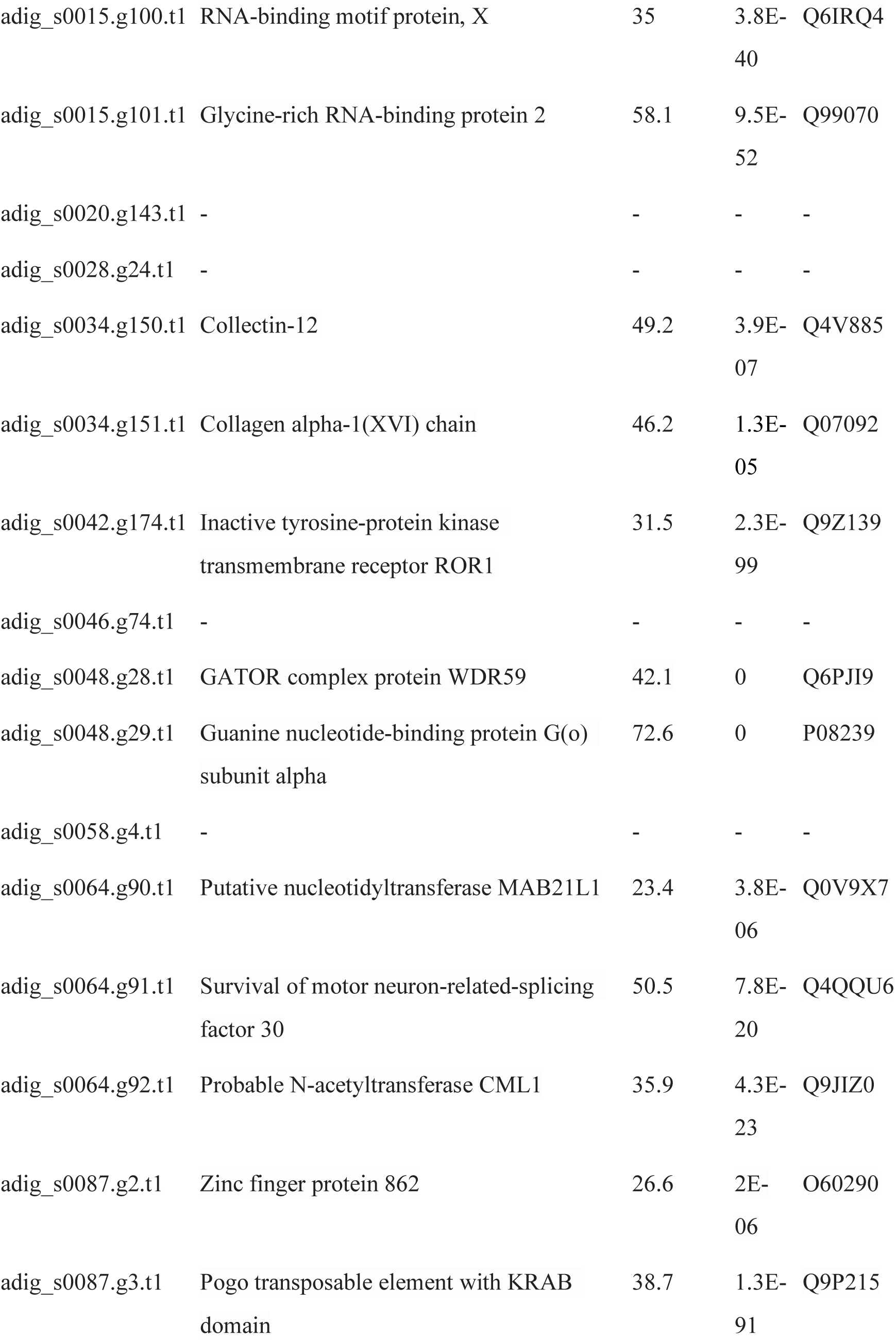
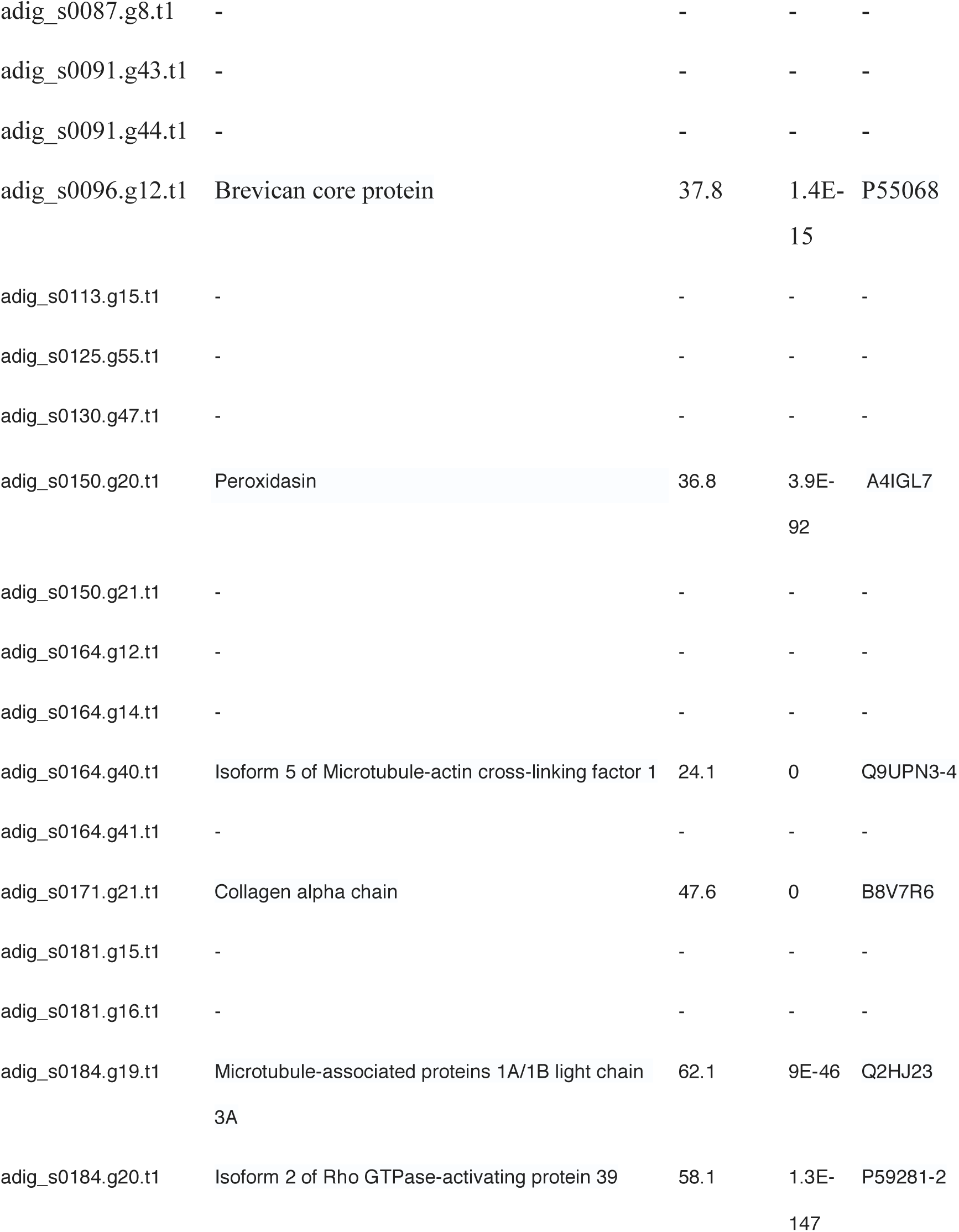

We surveyed the literature related to the annotated genes. Amino acid sequences of two genes (Gene IDs: adig_s0034.g151 and adig_s0171.g21) were similar (Table 1) to collagen alpha chain, which is associated with skeletogenesis in *Acropora* corals (Ramos-Silva, et al. 2013). The amino acid sequence of another gene (Gene ID: adig_s0048.g28) showed similarity (see Table 1) to a gene encoding WD repeat-containing protein 59 (WDR59).

To identify genes whose function is affected by differentiated SNPs, we identified amino acid changes between the two species caused by differentiated SNPs. Among 39 genes, 14 had at least one amino acid change between *A. digitifera* and *Acropora* sp. 1 (Table S4). Compared with the *A. digitifera* reference genome, *Acropora* sp. 1 had three amino acid changes in *WDR59* (Gene ID: adig_s0048.g28) (Table S4). WDR59 is a component of the GTPase-activating protein toward Rags (GATOR) complex, GATOR2 (Bar-Peled, et al. 2013). In *Drosophila*, GATOR2 controls meiotic entry and oocyte development (Wei, et al. 2014). Therefore, we focused further on this gene.

**Figure 3.**
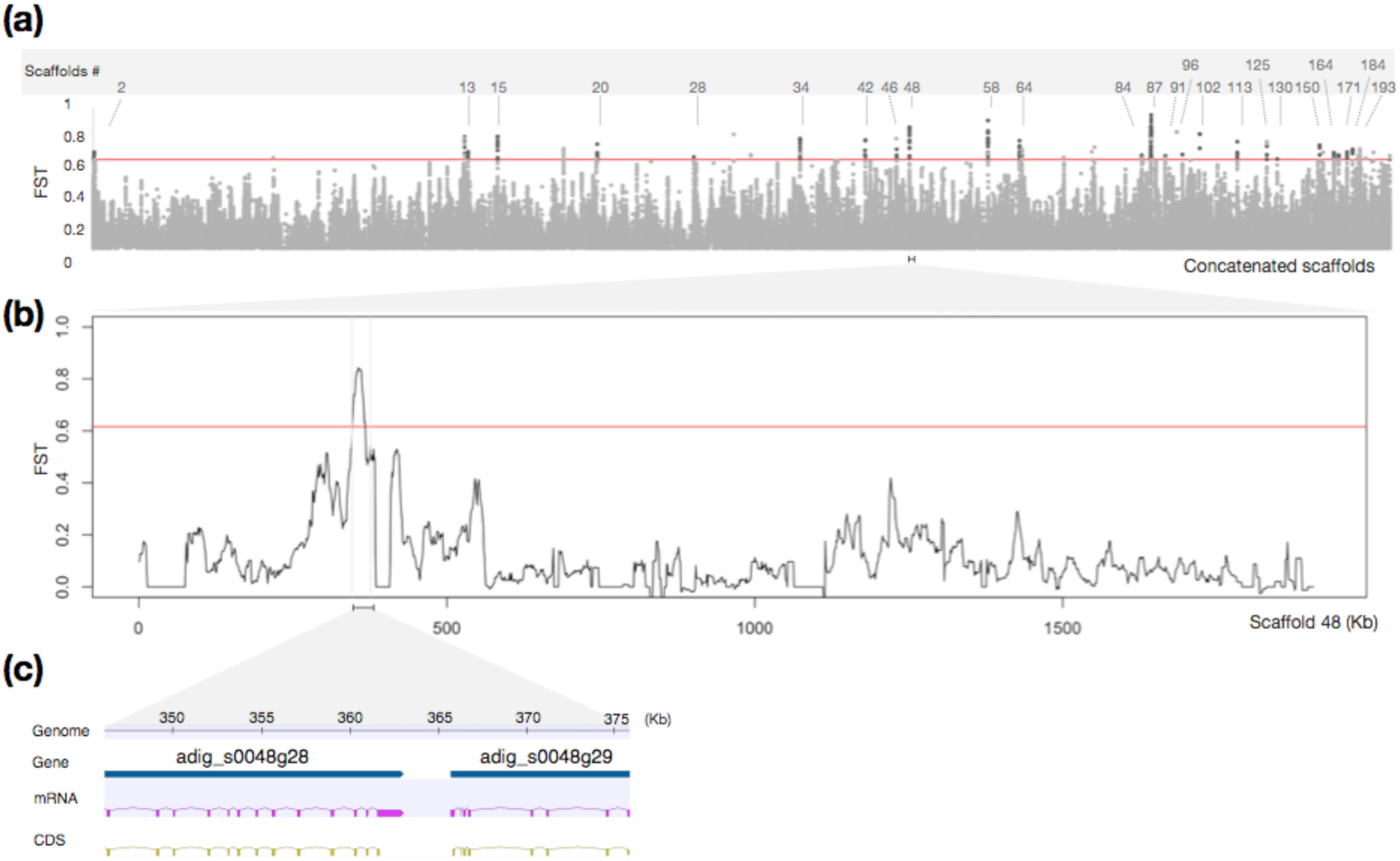
The genome-wide pattern of genetic differences between the two species. (a) Genome-wide FST values were calculated in overlapping windows of 10 kb. The red line indicates the top 0.1% of values. (b) FST was estimated across a region of scaffold 48 (adig_s0048). The red line indicates the top 0.1% of values. (c) A close-up view of predicted gene structures on an HDR in scaffold 48 (adig_s0048). The flanking gene structure of WDR59 (Gene ID: adig_s0048.g28) and guanine nucleotide-binding protein G(o) subunit alpha (Gene ID: adig_s0048.g29) are indicated.

### Differences in WDR59 between A. digitifera and Acropora sp. 1

To determine whether three amino acid differences in *WDR59* between *A. digitifera* and *Acropora* sp. 1 are shared with other species or are specific to *Acropora* sp. 1, we analyzed *WDR59* in 15 *Acropora* species (Table S5). First, we aligned the WDR59 sequence of 14 *Acropora* species (excluding *A. cytherea* due to a possible partial sequence) with that of *A. digitifera* and *Acropora* sp. 1 (Figure S2) and found that one of the three amino acid changes (adig_s0048.g28.t1: CDS; 2239 C>T, amino acid sequence; Pro747Ser) is specific to *Acropora* sp. 1 (Figure S2).

Next, we manually checked mapping reads around *WDR59* and found that *Acropora* sp. 1 colonies have a 24 bp deletion 38 bp downstream of the *Acropora* sp. 1-specific amino acid change. To verify this deletion, we amplified and sequenced the region containing the deletion by PCR from genomic DNAs of *A. digitifera* (n=7) and *Acropora* sp. 1 (n=14). We confirmed the deletion and found two additional amino acid differences between *A. digitifera* and *Acropora* sp. 1, upstream (15 bp) and downstream (14 bp) of the 24 bp deletion (Fig. S3). Among the differences between *A. digitifera* and *Acropora* sp. 1, two amino acid changes and a deletion are shared with *A. nasuta,* and one amino acid change is specific to *Acropora* sp. 1 (Figs. S4 and S5).

To estimate the position of the amino acid change specific to *Acropora.* sp. 1, we used Phyre2 (Kelley, et al. 2015) to search for proteins highly similar to *A. digitifera WDR59* in known structure databases. As a result, *S. cerevisiae* Sea3, the yeast counterpart of mammalian WDR59, was highly similar to *A. digitifera WDR59* (E-value=0, Identity=29%). *S. cerevisiae* Sea3 (WDR59) has an α-solenoid interface region where Sea3 (WDR59) interacts with the other subunit to form a complex, Sea2 (WDR24) (Tafur, et al. 2022). The α-solenoid interface region is located from amino acids 782 to 1,061 of *S. cerevisiae* Sea3 (WDR59) (Tafur, et al. 2022). An alignment of *A. digitifera* WDR59 with *S. cerevisiae* Sea3 (*WDR59*) (Fig. S6) showed that the amino acid changes specific to *Acropora* sp. 1 are located in the α-solenoid interface region.

## Discussion

### A. digitifera and Acropora sp. 1 are useful for understanding timing of gametogenesis in Acropora

Studying the timing of gamete maturation in corals using a population genetic approach, as in this study, provides insights into genetic mechanisms of coral gametogenesis and speciation in corals. Therefore, we propose *A. digitifera* and *Acropora* sp. 1 as a model species pair for studying mechanisms of spawning month determination and speciation in corals.

One of the advantages of using these two species is their clear phenotypic difference in timing of spawning. In Okinawa, *A. digitifera* spawns in May or June, whereas *Acropora* sp. 1 spawns in August (Hayashibara and Shimoike 2002; Nakajima, et al. 2012; Ohki, et al. 2015). Continuous observations of oocyte volume revealed that gamete maturation is later in *Acropora* sp. 1 than in *A. digitifera* (Hayashibara and Shimoike 2002). The difference in gamete maturation is expected to lead to reproductive isolation. Indeed, phylogenetic analysis and PCA showed that the two species are genetically differentiated, despite their low genetic differentiation. Therefore, gene flow between *A. digitifera* and *Acropora* sp. 1 is limited, which is considered an initial stage of speciation.

The low genetic differentiation between *A. digitifera* and *Acropora* sp. 1 is another advantage in studying genes responsible for spawning timing mechanisms and speciation. Genomic differentiation between these two species is low (F_ST_ = 0.10225), consistent with a previous microsatellite marker study (Nakajima, et al. 2012). Using this low-genomic differentiated species pair, we identified 34 HDRs and selected 39 genes located in HDRs. These genomic regions and candidate genes may be responsible for morphological and ecological differences between the two species. Further analyses of gene expression differences in different months, functional changes resulting from highly differentiated substitutions are expected to advance research on the mechanism of spawning month determination and speciation in corals.

### Genes that may determine morphological differentiation between two species

Morphological characteristics of *Acropora* sp. 1 include shorter branches and a flatter colony shape than *A. digitifera* (Hayashibara and Shimoike 2002; Ohki, et al. 2015). These morphological differences reflect differences in skeletal form (Todd 2008). The alpha collagen-like proteins are skeletal organic matrix proteins involved in skeletal formation in *Stylophora pistillat*a (Drake, et al. 2013; Mummadisetti, et al. 2021) and *A. millepora* (Ramos-Silva, et al. 2013). In this study, we identified two alpha collagen-like genes (Gene IDs: adig_s0034.g151 and adig_s0171.g21) in HDRs, and these genes are likely responsible for species-specific differences in skeletal morphology. *ATOH1* (Gene ID: adig_s0013.g139), encodes the transcription factor Atoh1, which regulates primary cilia of calcifying cells in mice (Chang, et al. 2019). Since the possibility of cilia in coral skeletogenesis has been discussed in *S. pistillata* (Tambutté, et al. 2021), *ATOH1* may help to define skeletal morphology in the two species.

**mTOR**C1 may contribute to gametogenesis of *A. digitifera*

In this study, we identified an amino acid change specific to *Acropora* sp. 1 in *WDR59*. WDR59 is one of the components of a mechanistic target-of-rapamycin complex 1 (mTORC1) activator, GATOR2 (Bar-Peled, et al. 2013; Wolfson, et al. 2016) (Fig. 4). mTORC1 is involved in meiotic entry and gametogenesis. Regulation of meiotic entry by mTORC1 is conserved from yeast to mammals. Downregulation of mTORC1 activity promotes the transition from mitotic to meiotic cycles in *Saccharomyces cerevisiae*, *Schizosaccharomyces pombe* (van Werven and Amon 2011; Zheng and Schreiber 1997), and *Drosophila* (Wei, et al. 2014). In mice, mTORC1 is required for spermatogonial differentiation (Busada, et al. 2015) and oogenesis (Guo, et al. 2018). Activated mTORC1 drives oocyte development and growth in *Drosophila* oogenesis (LaFever, et al. 2010). To the best of our knowledge, the function of mTORC1 in gametogenesis among Cnidarians has been little discussed. One exception is a study about the kinase, Mos, which regulates oocyte maturation in the jellyfish, *Clytia hemisphaerica* (Amiel, et al. 2009). Treatment of oocytes with rapamycin, a potent inhibitor of mTORC1, suggested that the mTORC1 signaling pathway controls one *Mos* paralog translation during oocyte growth (Amiel, et al. 2009). Moreover, in *Hydra oligactis*, continuous exposure to rapamycin results in fewer mature sperm cells than in untreated individuals (Tomczyk, et al. 2020). Hence, mTORC1 is likely associated with gametogenesis in cnidarians, including *Acropora* species.

**Figure 4.**
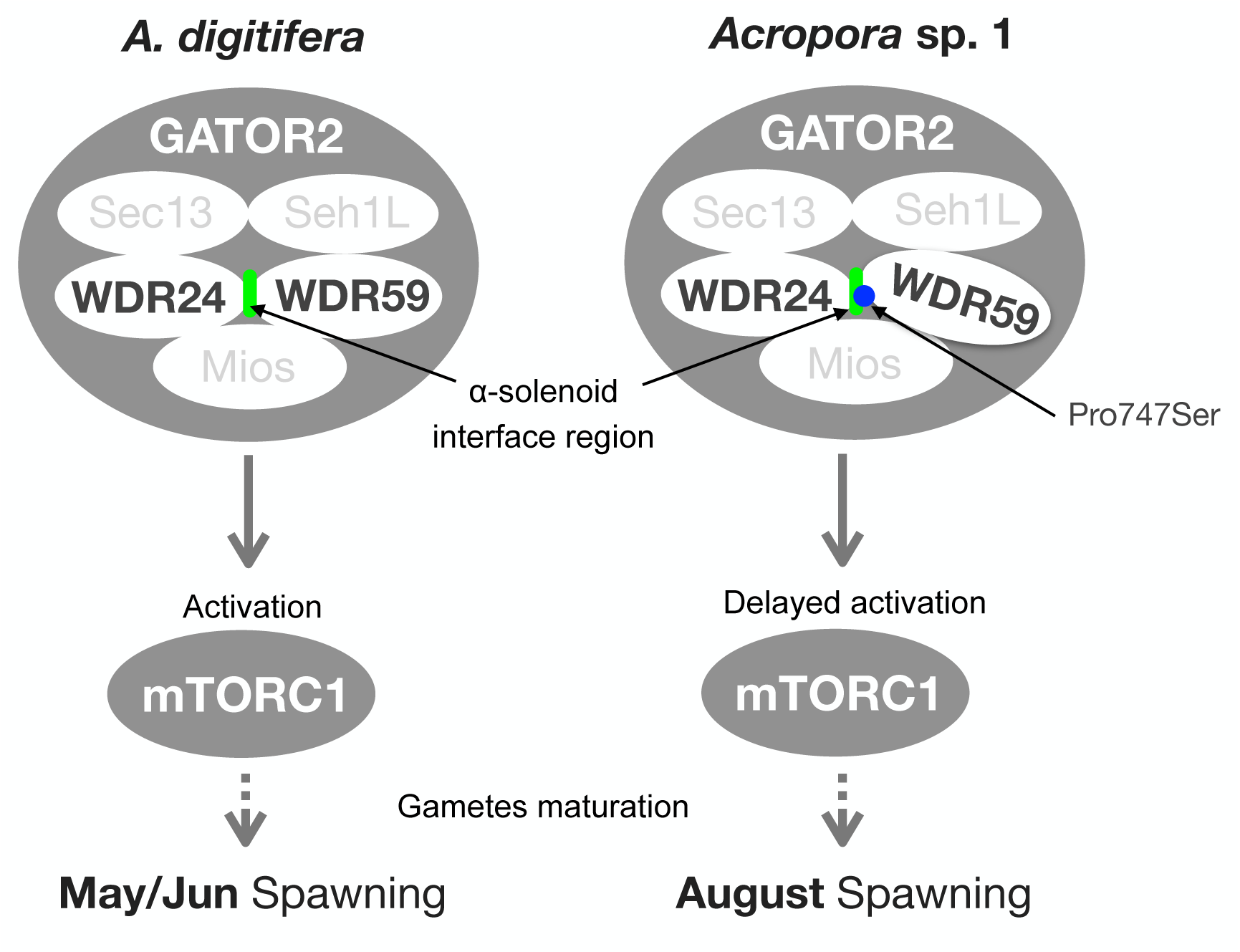
Schematic representation of a hypothesis proposed in this study. Regulation of mTORC1 by GATOR2 and components of GATOR2 is based on previous studies (Bar-Peled, et al. 2013; Valenstein, et al. 2022; Wei, et al. 2014). An *Acropora* sp. 1-specific mutation in the WDR59 / WDR24 interaction region is indicated with a blue circle.

The *Acropora* sp. 1-specific amino acid change in WDR59 is located in a region where WDR59 interacts with one of the other GATOR2 components to form the complex (GATOR2). This amino acid change may cause slight differences in stability or structure of GATOR2 through affinity of WDR59 with its counterpart. In *Drosophila* oogenesis, GATOR2 activates mTORC1, and active mTORC1 is required to start oocyte development (Wei, et al. 2014). Since regulation of gametogenesis by mTORC1 is reported in *Drosophila*, meiotic entry and oocyte development in *Acropora* species is also likely controlled by mTORC1 activity, regulated by GATOR2. In other words, the difference in timing of gamete maturation in *A. digitifera* and *Acropora* sp. 1 (Hayashibara and Shimoike 2002) may be caused by an amino acid substitution in WDR59 that slightly affects timing of mTORC1 activation via GATOR2. Note that even though we focused on WDR59 in this study, a combination of genetic factors, including genes in other HDRs, may be responsible for differences in spawning timing. Since the phylum Cnidaria, including corals, is located in the basal lineage of the animal kingdom, studies revealing the function of mTORC1 in gametogenesis in corals will provide insights into evolution of gametogenesis regulation. Future studies of the two coral species used in this study will shed light on mechanisms that determine the timing of coral spawning.

## Materials and Methods

### Specimen collection and species identification

Coral samples were collected from two reefs at Okinawa, Japan, between 2018 to 2020 (Table S1) with permission of the Aquaculture Agency of Okinawa Prefecture (permit numbers 30-29, 31-43, and 31-68). Sixteen colonies of *Acropora* sp. 1 with visible gametes, were collected in the field and subsequently maintained in an aquarium at the Sesoko Station, Tropical Biosphere Research Center, University of the Ryukyus. In 2018, gametes of one *Acropora* sp. 1 colony were collected during spawning, and sperm were preserved at -80°C until genome extraction. After we placed the coral colonies in the aquarium, we preserved branch fragments in RNAlater (Waltham, MA, USA) for genome extraction in 2019 and 2020.

### DNA extraction and sequencing

We extracted genomic DNAs from 15 branch fragments originating from 15 *Acropora* sp. 1 colonies using a DNeasy Plant Mini Kit (QIAGEN, Hilden, Germany). We used DNeasy Blood & Tissue Kits (QIAGEN, Hilden, Germany) for DNA extraction from sperm originating from one *Acropora* sp. 1 colony. Following the manufacturer’s instructions, we constructed DNA libraries from 16 samples using an NEBNext Ultra II DNA Library Prep Kit (Illumina). The 15 libraries from branch tissues were sequenced on an Illumina HiSeqX Ten, and one library from sperm was sequenced on an Illumina HiSeq 2500.

### Mapping and variant calling

We downloaded genome sequence data from 11 colonies of *A. digitifera* and 15 *Acropora* species (*A. tenuis, A.yongei, A. intermedia, A. gemmifera, A. awi, A. florida, A. millepora, A. selago, A. hyacinthus, A. cytherea, A. muricate, A. echinate, A. acuminata, A. nasuta, and A. microphthalma*). We trimmed raw sequences and removed low-quality reads before mapping with fastp (Chen, et al. 2018). Trimmed reads were mapped to the *A. digitifera* genome assembly ver. 2.0 (Shinzato, et al. 2021) using bowtie2 ver. 2.3.3.1 (Langmead and Salzberg 2012). Among 16 *Acropora* sp. 1 colonies, we used 14 colonies with mapping bam coverage ≥10 for variant calling. Variants were called using Genome Analysis Toolkit (GATK) version 4.0 and filtered according to a GATK-suggested hard-filtering with a minor modification.

### PCA and molecular phylogenetic tree construction

We constructed a molecular phylogenetic tree of these *Acropora* corals using phyML (Guindon, et al. 2010) with the GTR option (Guindon, et al. 2010). We performed PCA analysis of *A. digitifera* and *Acropora* sp. 1 with three species, *A. acuminata*, *A. microphthalma,* and *A. nasuta,* as an out-group, using PLINK v1.90 (www.cog-genomics.org/plink/1.9/) (Weeks 2010).

### Genome scan of highly differentiated regions

We calculated F_ST_ (Hudson, et al. 1992) for 10-kb windows with 1 kb increments along each scaffold (>10 kb) using a sliding window approach with PopGenome (Pfeifer, et al. 2014). First, we extracted 10 kb windows that included the top 0.1% of F_ST_ values. Among these top windows, we selected windows with SNPs for which the allele is fixed in one population and for which there is no homozygote for the allele in the comparison population. We considered these SNPs to be differentiated SNPs. We merged overlapping regions among these selected windows and considered these connected regions highly differentiated.

### Identification of genes in highly differentiated regions (HDRs)

We considered genes with differentiated SNPs in HDRs as candidate genes related to phenotypic differences between the two species. To identify functional annotations of these genes, we searched orthologous genes in the NCBI nucleotide database and UniProt (Bateman, et al. 2022) by Blast search (Altschul, et al. 1990). We regarded the top hit with an *e*-value ≥ 1e^-30^ and identity ≥ 90% for NCBI and *e*-value ≥ 1e^-4^ and identity ≥ 20% for UniProt as an orthologous gene.

### Identification of a deletion in WDR59 among Acropora sp. 1

The presence of one deletion in the WDR59 gene in *Acropora* sp. 1, discovered by visual confirmation of the mapping results, was revealed by amplifying the genomic region containing the deletion using PCR and sequencing it.

### Alignment of WDR59 sequences among Acropora species

To determine whether other *Acropora* species have genetic variants other than those that differentiate *A. digitifera* and *Acropora*. sp. 1, orthologous genes of *WDR59* were searched in the reference genomes of each of the 15 *Acropora* species using Blastn (Altschul, et al. 1990). A *WDR59 sequence of A. millepora* was downloaded from the Kyoto Encyclopedia of Genes and Genomes (KEGG).

## Acknowledgments and funding sources

We thank Jun Ishida for providing the photo of *Acropora* sp. 1. We thank Ryo Kariyazono for helpful discussion. This work was supported by JSPS KAKENHI Grant Number 22J40115. Computations were partially performed on the NIG supercomputer at ROIS National Institute of Genetics.

STK: research concept, all experiments, data analysis, and manuscript preparation.

AI: sample collection planning, species identification.

YT: research concept, research planning, data analysis, and manuscript preparation.

## Spell out all abbreviations

CDS: Coding sequence
DDBJ: DNA Data Bank of Japan
GATK: Genome Analysis Toolkit
HDRs: Highly Differentiated Regions
KEGG: Kyoto Encyclopedia of Genes and Genomes
mTORC1: mechanistic target of rapamycin complex 1
NCBI: National Center for Biotechnology Information
PCA: principal components analysis
SNP: single nucleotide polymorphism
SEA/GATOR: the <u>Se</u>h1 <u>a</u>ssociated/<u>G</u>TPase-<u>a</u>ctivating protein <u>to</u>ward <u>R</u>ags
Sea2: SEA (*Se*h1-*a*ssociated) protein complex 2
Sea3: SEA (*Se*h1-*a*ssociated) protein complex 3
WDR24: WD repeat-containing protein 24
WDR59: WD repeat-containing protein 59

