## Supplementary material for "Comparative genomics of two closely related coral species with different spawning seasons reveals genomic regions possibly associated with gametogenesis": Asp1Spawning_Supp.pdf

Contents

1 Materials and Methods
1.1 Samples
1.1.1 Sample collection
1.1.2 DNA Extraction and Sequencing
1.2 Alignment and variant calling of *Acropora digitifera* and *Acropora* sp.1
1.2.1 Short reads alignment
1.2.2 Variant calling
1.2.3 Variant Filtering

1.3 Alignment and variant calling of 15 other Acropora species
1.4 SNP analysis
1.4.1 Phylogenetic analysis
1.4.2 PCA analysis
1.4.3 Genome scan of highly differentiated regions
1.4.4 Genes in highly differentiated regions (HDRs)
1.5 Variation of WDR59

1.5.1 Identification of a deletion in WDR59 among *Acropora* sp.1

1.5.2 Identification and alignment of WDR59 in other *Acropora* species

2 Supplemental Figures

3 Supplemental Tables

### 1 Materials and Methods

#### 1.1 Samples

##### 1.1.1 Sample collection

In total, 16 samples from 16 *Acropora* sp.1 colonies were collected during 2018-2020 in Okinawa, Japan, under permission of the Aquaculture Agency of Okinawa Prefecture (permit numbers 30-29, 31-43, and 31-68). Dr. Akira Iguchi identified *Acropora* sp.1 colonies based on their morphology.

On 30 July 2018, one *Acropora* sp.1 (Colony ID: Asp1_c) colony was collected from Bise-zaki and maintained in an aquarium at the Sesoko Station, Tropical Biosphere Research Center, University of the Ryukyus. We observed spawning of this colony on 5 August, and sperm was collected from this colony and stored at -80.

On 3 August 2019, we collected seven colonies (Colony IDs: Asp1B1901-Asp1B1907) from Bise-zaki, and a branch fragment was collected and preserved in RNAlater (Waltham, MA USA) from each of seven colonies. We maintained these colonies in an aquarium at the Sesoko Station for spawning observations.

At the beginning of August 2020, we collected eight colonies (individual IDs: Asp1S2001- Asp1S2008) from the Sesoko-jima reef. A branch fragment was collected and preserved in RNAlater (Waltham, MA, USA) from each of the seven colonies.

##### 1.1.2 DNA Extraction and Sequencing

We extracted genomic DNA from *Acropora* sp.1 colonies for the genome DNA library. We used DNeasy Plant Mini Kits (QIAGEN, Hilden, Germany) for DNA extraction from 15 branch fragments. For DNA extraction from the sperm sample, we used DNeasy Blood & Tissue Kits (QIAGEN, Hilden, Germany). Following the manufacturer's instructions, we constructed DNA libraries of 16 samples using an NEBNext Ultra II DNA Library Prep Kit (New England Biolabs, Ipswich, MA, USA) and NEBNext Multiplex Oligos for Illumina 96 Unique Dual Index Primer Pairs (New England Biolabs). The 15 libraries from branch tissues were sequenced on an Illumina HiSeqX Ten, and one library from sperm was sequenced on an Illumina HiSeq 2500.

#### 1.2 Alignment and variant calling of *Acropora digitifera* and *Acropora* sp.1

##### 1.2.1 Short-read alignment

In addition to our sequencing data from *Acropora* sp.1, we downloaded complete genome sequences of 11 colonies of *A. digitifera* from the DNA Data Bank of Japan (DDBJ) (Table S1). Raw 150 bp paired-end Illumina short reads saved as FASTQ files were trimmed with fastp (Chen, et al. 2018) to remove low-quality reads and Illumina adapters using modified parameters (-l 50 -q 30). Trimmed reads were then aligned to the *A. digitifera* genome assembly ver. 2.0 (https://marinegenomics.oist.jp/adig/viewer/download?project_id=87) (Shinzato, et al. 2021) with bowtie2 ver. 2.3.3.1 (Langmead and Salzberg 2012) using modified parameters (--score-min L,0,-0.2). Alignment results were saved as Sequence Alignment/Map (SAM) format files and converted to Binary Alignment/Map (BAM) format files using samtools 1.3.1(Danecek, et al. 2021). If the same sample-derived reads are split into multiple FASTQ files, each read file is aligned separately, and the output BAM files are sorted, indexed, and concatenated for each sample with samtools 1.3.1 (Danecek, et al. 2021) Individual information (read group) was added to each of the BAM files by GATK ver. 4.1.6.0 (McKenna, et al. 2010).

##### 1.2.2 Variant calling

The HaplotypeCaller program, which is provided in GATK ver. 4.1.6.0 (McKenna, et al. 2010) was used to call genetic variants in each sample. Called variants from each sample were saved as individual files (called gvcf files), and individual files were combined using GATK CombineGvcf program. GATK GenotypeGVCFs program was used to call genotypes among all combined samples, and genotyped variants were saved as one file (called "vcf file").

##### 1.2.3 Variant Filtering

Following GATK "best practices" (DePristo, et al. 2011; Poplin, et al. 2017), hard-filtering with minor modifications was applied to genotyped variants. For genotype variants, only the threshold for low mapping quality (MQ) was changed from 40 to 20, while other thresholds were left at default settings. After GATK hard-filtering, we excluded sites with missing sites and indels, low minor allele frequency (maf < 0.05), and low depth values (minDP < 3) using VCFtools version 0.1.16 (Danecek, et al. 2011). We further selected biallelic SNPs with bcftools version 1.9 (Danecek, et al. 2021).

To remove SNPs, a cutoff value for maxDP was set for each vcf file, and sites with DP values higher than twice the average DP of the individual's biallelic SNPs were excluded from the analysis. First, vcf files for each individual were extracted from the combined vcf files with VCFtools version 0.1.16 (Danecek, et al. 2011). For each individual vcf file, the following operations were performed. The vcftools option, “site-depth”, was used to extract the depth of each site, the average DP was calculated, and a bed file was created showing sites where the DP value was higher than twice the average DP of the bi-allelic SNP for that individual. All bed files for each individual were combined, and duplicate entries in the combined bed files were deleted. Bi-allelic SNPs were selected from the combined vcf, excluding the region specified in the bed file.

To remove SNPs that were not under Hardy-Weinberg equilibrium we checked the p-value for each site from a Hardy-Weinberg Equilibrium test (Chiu, et al. 2020) using VCFtools version 0.1.16 (Danecek, et al. 2011). First, the maxDP-filtered vcf file was split into two vcf files for each of the two species. The vcf file for each species was used to determine a p-value for each site from the Hardy-Weinberg Equilibrium test. Only sites with a p-value < 0.05 were used to calculate $F_{ST}$ s (explained in 1.4 .3 Genome scan of the highly differentiated region).

##### 1.3 Alignment and variant calling of 15 other *Acropora* species

We downloaded genome sequences of 15 other *Acropora* species from DDBJ. Downloaded raw reads were trimmed in the same manner as described above (1.2.1 Short reads alignment). Trimmed reads were then aligned to the *A. digitifera* genome assembly ver. 2.0 (Shinzato, et al. 2021) with bowtie2 ver. 2.3.3.1 (Langmead and Salzberg 2012) using the default setting. A gvcf file containing genotyped variants was generated for each of the 15 species and combined with the *A. digitifera* and *Acropora* sp.1 genotyped variants in the same manner described above (1.2.1 Short read alignment and 1.2.2 variant calling) with minor changes. Low minor allele frequency(maf) was 0.03, and the Hardy-Weinberg Equilibrium test was not performed.

##### 1.4 SNP analysis

###### 1.4.1 Phylogenetic analysis

We converted a vcf file containing biallelic SNPs among 17 *Acropora* species into phylip format using Tassel5 (Bradbury, et al. 2007) to import phyML (Guindon, et al. 2010). We constructed a molecular phylogenetic tree of these *Acropora* corals with phyML (Guindon, et al. 2010) using the GTR option and created a phylogenetic tree with MEGA 7 (Kumar, et al. 2016).

##### 1.4.2 PCA analysis

Based on the phylogenetic relationship among 17 *Acropora* species, we use *A. acuminata*, *A. microphthalma,* and *A. nasuta* as an out-group. We extracted 24,955,282 SNPs from five species (*A. digitifera*, *Acropora* sp. 1, and the three *Acropora* out-group species). We performed principal components analysis (PCA) on the genome-wide pruned 80,490 SNPs using PLINK v1.90 (http://pngu.mgh.harvard.edu/purcell/plink/)(Weeks 2010).

##### 1.4 .3 Genome scan of highly differentiated region*s*

We calculated $F_{ST}$ s (Hudson, et al. 1992) for 10-kb windows with 1-kb increments along each scaffold (>10 kb) using a sliding-window approach by PopGenome (Pfeifer, et al. 2014). First, we extracted a 10-kb window containing the top 0.1% of the $F_{ST}$ distribution and created a bed file showing the genomic positions of the top 0.1% windows. Using the bed file, we extracted information on SNPs in the top 0.1% windows and genotype information for each sample using bcftools version 1.9 (Danecek, et al. 2021). Based on the genotype information, we selected SNPs for which the allele is fixed in one population and for which there was no homozygote for the allele in the other population. We considered these SNPs as differentiated SNPs. Among the top 0.1% of windows, we extracted windows with differentiated SNPs and merged the overlapping regions. These combined regions were considered highly differentiated regions (HDRs).

##### 1.4 .4 Genes in highly differentiated regions (HDRs)

We considered genes with differentiated SNPs in HDRs as candidate genes related to differences between the two species. To identify the functional annotation of these genes, we searched orthologous genes in National Center for Biotechnology Information (NCBI) nucleotide database and UniProt (Bateman, et al. 2022) by Blast search (Altschul, et al. 1990). The top hits (with $e \geq 1e^{-30}$ and identity ≥ 90% for NCBI, and $e \geq 1e^{-5}$ and identity ≥ 20% for Uniprot) were regarded as orthologous genes. We determined whether differentiated SNPs cause amino acid changes using CLC Genomics Workbench 11.0 (QIAGEN, Aarhus, Denmark).

#### 1.5 Variation of WDR59

1.5.1 Identification of a deletion in *WDR59* among *Acropora* sp.1

The presence of one deletion in the WDR59 gene among *Acropora*. sp. 1 was confirmed by visual inspection of mapping results. To identify the deletion, we amplified the genomic region containing the expected deletion using PCR with PrimeSTAR GXL DNA Polymerase (Takara, Shiga, Japan) and the following primers: 5’-CTCCATATTCTAACATCTCTG-3' and 5'-AAAACGTAGCTTGCTAAAGC-3'. PCR was performed using GeneAmp PCR System 9700 (Applied Biosystems, Carlsbad, CA, USA) with the following conditions: denaturation for 1 min 30 sec at 93 °C, followed by 30 cycles of denaturation for 30 sec at 93 °C, annealing for 30 sec at 55 °C, and extension for 30 sec at 72 °C. We used the following genomic DNAs as templates for PCR: Genomic DNAs extracted from 7 *A. digitifera* colonies (sample ID: AdigS1601–4, AdigS1606–07 and AdigS1610) and 14 Acropora sp. 1 colonies (sample ID: Asp1B1901-07, Asp1c, Asp1S2001-03, Asp1S2005, and Asp1S2007-08). We determined the sequences of PCR products using a Genetic Analyzer 3500 (Thermo Fisher Scientific) with the same primers used for PCR. All determined sequences were aligned with the reference sequence (*A. digitifera* scaffold: sc0000048_arrow_pilon) using ClustalW with default parameters in MEGA ver. 7 (Kumar, et al. 2016).

##### 1.5.2 Identification and alignment of WDR59 in other *Acropora* species

The genome assemblies of 14 *Acropora* species (*A. tenuis, A.yongei, A. intermedia, A. gemmifera, A. awi, A. florida, A. selago, A. hyacinthus, A. cytherea, A. muricate, A. echinate, A. acuminata, A. nasuta, and A. microphthalma*) were downloaded from the OIST Marine Genomics Unit Genome Browser (https://marinegenomics.oist.jp/gallery). Orthologous genes of *WDR59* were searched in the reference genome of each of the 14 *Acropora* species by Blastn search (Altschul, et al. 1990). Top hit sequences were used for alignment as *WDR59* orthologs in each species (Fig S2, Tables S4-5). A *WDR59* sequence of *A. millepora* was downloaded from KEGG.

#### 2 Supplemental Figures

### Fig S1

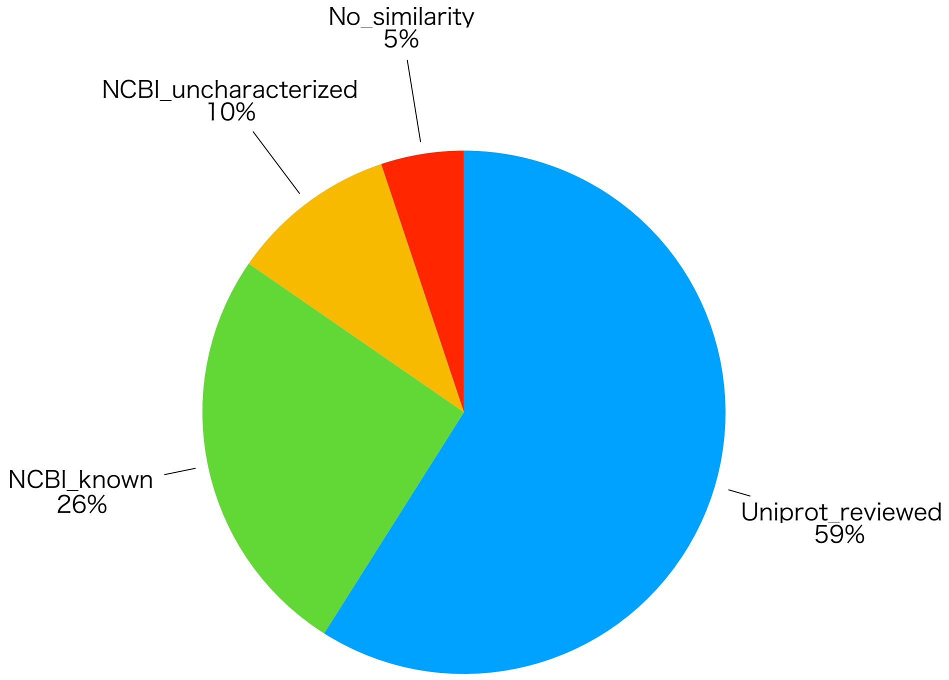


**Figure S1.** In the HDRs, 39 genes harbor differentiated SNPs. Among 39 genes, 23 genes are similar (identity ≥ 23%, E-value ≤ 1.1E-5) to genes in the UniProtKB/Swiss-Prot database (https://www.uniprot.org/blast) with high-quality manually annotated (reviewed) annotations. Ten genes are similar (identity ≥ 90%, E-value = 0) to genes in the NCBI nucleotide database with automatically annotated annotations related to known genes. Four genes are similar (identity ≥ 90%, E-value = 0) to uncharacterized genes. Two genes have no similarity (default settings) to any genes in the NCBI nucleotide database.

### Fig S2

```
aten_s0183.g20.t1  ---------- MLTDDDIHSE TKMAS----- ---------- -YQDAVAPHK
ayon_s0004.g209.t1 ---------- ---------- ---------- ---------- ----------
aint_s0143.g6.t1   ---------- ---------- --MAS----- ---------- -YQDAVAPHK
agem_s0013.g175.t1 ---------- ---------- --MAS----- ---------- -YQDAVAPHK
aawi_s0007.g175.t1 ---------- ---------- --MAS----- ---------- -YQDAVAPHK
aflo_s0310.g20.t1  ---------- -MLTDDIHPE TKMAS----- ---------- -YQDAVAAHK
XM_029340177.2     ---------- -MLTDDIPPE TKMAS----- ---------- -YQDAVAA-H
asel_s0045.g58.t2  ---------- -MLTDDIPPE TKMAS----- ---------- -YQDAVAA-H
ahya_s0003.g68.t2  ---------- ---------- --MAS----- ---------- -YQDAFAAHK
acyt_s1402.g1.t1   VICNALRLWD IKLVAFPSPE AKEARENPQN QRAFICNLQQ HWLTIRKLGH
amur_s0006.g105.t2 ---------- ---------- --MAS----- ---------- -YQDAVAAHK
ech_s0159.g30.t2   ---------- ---------- --MAS----- ---------- -YQDAVAAHK
aacu_s0038.g69.t1  ---------- -MLTDDIPPE TKMAS----- ---------- -YQDAVAA-H
anas_s0109.g72.t2  ---------- -MLTDDIPPE TKMAS----- ---------- -YQDAVAA-H
amic_s0245.g8.t1   ---------- -MLTDDIHPE TKMAS----- ---------- -YQDAVAA-H
adig_s0048.g28.t1  ---------- -MLTDDIHPE TKMAS----- ---------- -YQNAVA-VH

aten_s0183.g20.t1  DWPA------ ---------- ----TVMAVD GTGHFTVLGS RKGL--AFID
ayon_s0004.g209.t1 ---------- ---------- ---------- ---------- ----------
aint_s0143.g6.t1   DWPA------ ---------- ----TVMAVD CTGHFTVLGA RKGL--AFID
agem_s0013.g175.t1 DWPA------ ---------- ----TVMAVD CTGHFTVLGS RKGL--AFID
aawi_s0007.g175.t1 DWPA------ ---------- ----TVMAVD CTGHFTVLGA RKGL--AFID
aflo_s0310.g20.t1  DWPA------ ---------- ----TVMAVD CTGHFTVLGA RKGL--AFID
XM_029340177.2     DWPA------ ---------- ----TVMAVD CTGHFTVLGS RKGL--AFID
asel_s0045.g58.t2  DWPA------ ---------- ----TVMAVD CTGHFTVLGS RKGL--AFID
ahya_s0003.g68.t2  DWPA------ ---------- ----TVMAVD CTGHFTVLGS RKGL--AFID
acyt_s1402.g1.t1   QWFNLNSLLA KPELITETYL LLYLTQLQTD GYSIFVVMGR LPESKADHML
amur_s0006.g105.t2 DWPA------ ---------- ----TVMAVD CTGHFTVLGS RKGL--AFID
ech_s0159.g30.t2   DWPA------ ---------- ----TVMAVD CTGHFTVLGS RKGL--AFID
aacu_s0038.g69.t1  DWPA------ ---------- ----TVMAVD CTGHFTVLGS RKGL--AFID
anas_s0109.g72.t2  DWPA------ ---------- ----TVMAVD CTGHFTVLGS RKGL--AFID
amic_s0245.g8.t1   DWPA------ ---------- ----TVMAVD CTGHFTVLGS RKGL--AFID
adig_s0048.g28.t1  DWPA------ ---------- ----TVMAVD CTGHFTVLGS RKGL--AFID

aten_s0183.g20.t1  LSCPSVITKK VPRNSKWECN ALEWNPHLSH AHIFANASNQ KTEIWSWSNG
ayon_s0004.g209.t1 ---------- ---------- ---------- ---------- ----------
aint_s0143.g6.t1   LNCPNVITKK VPRNSKWECN ALEWNPHLSD AHIFANASNQ KTEIWSWSNG
agem_s0013.g175.t1 LNCPNVITKK VPRNSKWECN ALEWNPHLSD AHIFANASNQ KTEIWSWSNG
aawi_s0007.g175.t1 LNCPNVITKK VPRNSKWECN ALEWNPHLSD AHIFANASNQ KTEIWSWSNG
aflo_s0310.g20.t1  LNCPNVITKK VPRNSKWECN ALEWNPHLSD AHIFANASNQ KTEIWSWSNG
XM_029340177.2     LNFPNVITKK VPRNSKWECN ALEWNPHLSD AHIFANASNQ KTEIWSWSNG
asel_s0045.g58.t2  LNFPNVITKK VPRNSKWECN ALEWNPHLSD AHIFANASNQ KTEIWSWSNG
ahya_s0003.g68.t2  LNFPNVITKK VPRNSKWECN ALEWNPHLSD AHIFANASNQ KTEIWSWSNG
acyt_s1402.g1.t1   KMCPAQLVKK PANKNTF--- ----QKSISK SDL------- ---------T
amur_s0006.g105.t2 LNFPNVITKK VPRNSKWECN ALEWNPHLSD AHIFANASNQ KTEIWSWSNG
ech_s0159.g30.t2   LNFPNVITKK VPRNSKWECN ALEWNPHLSD AHIFANASNQ KTEIWSWSNG
aacu_s0038.g69.t1  LNFPNVITKK VPRNSKWECN ALEWNPHLSD AHIFANASNQ KTEIWSWSNG
anas_s0109.g72.t2  LNFPNVITKK VPRNSKWECN ALEWNPHLSD AHIFANASNQ KTEIWSWSNS
amic_s0245.g8.t1   LNFPNVITKK VPRNSKWECN ALEWNPHLSD AHIFANASNQ KTEIWSWSNG
adig_s0048.g28.t1  LNSPSVITKK VPRNSKWECN ALEWNPHLSD AHIFANASNQ KTEIWSWSNG

aten_s0183.g20.t1  NGLQLQILRG HTRAISDLNW SRFDPQLLST CSMDQFIYIW DLREGK----
ayon_s0004.g209.t1 ---------- ---------- ---------- ---------- ----------
aint_s0143.g6.t1   NGLQLQILRG HTRAISDLNW SWFDPQLLSS CSMDQFIYIW DLREGK----
agem_s0013.g175.t1 NGLQLQILRG HTRAISDLNW SRFDPQLLSS CSMDQFIYIW DLREGK----
aawi_s0007.g175.t1 NGLQLQILRG HTRAISDLNW SWFDPQLLSS CSMDQFIYIW DLREGK----
aflo_s0310.g20.t1  NGLQLQILRG HTRAISDLNW SWFDPQLLSS CSMDQFIYIW DLREGK----
XM_029340177.2     SGLQLQILRG HTRAISDLNW SWFDPQLLSS CSMDQFIYIW DLREGK----
asel_s0045.g58.t2  SGLQLQILRG HTRAISDLNW SWFDPQLLSS CSMDQFIYIW DLREGK----
ahya_s0003.g68.t2  NGLQLQILRG HTRAISDLNW SWFDPQLLSS CSMDQFIYIW DLREGK----
acyt_s1402.g1.t1   AALQQATVGG RDKP-AD--- -------MEE IRRRRELYFT RQQQNQENDT
amur_s0006.g105.t2 NGLQLQILRG HTRAISDLNW SWFDPQLLSS CSMDQFIYIW DLREGK----
ech_s0159.g30.t2   NGLQLQILRG HTRAISDLNW SWFDPQLLSS CSMDQFIYIW DLREGK----
aacu_s0038.g69.t1  SGLQLQILRG HTRAISDLNW SWFDPQLLSS CSMDQFIYIW DLREGK----
anas_s0109.g72.t2  SGLQLQILRG HTRAISDLNW SWFDPQLLSS CSMDQFIYIW DLREGK----
amic_s0245.g8.t1   NGLQLQILRG HTRAISDLNW SWFDPQLLSS CSMDQFIYIW DLREGK----
adig_s0048.g28.t1  NGLQLQILRG HTRAISDLNW SWFDPQLLSS CSMDQFIYIW DLREGK----
```

### Fig S2 continued

```
aten_s0183.g20.t1  -KPASSLQAI VGASQVKWNR VNRHVLATSH DGDVRIWDLR KGNTPVVYLT
ayon_s0004.g209.t1 ---------- ---------- ---------- ---------- ----------
aint_s0143.g6.t1   -KPASSLQAI VGASQVKWNR VNRHVLATSH DGDVRIWDLR KGNTPVVYLT
agem_s0013.g175.t1 -KPASSLQAI VGASQVKWNR VNRHVLASSH DGDVRIWDLR KGNTPVVYLT
aawi_s0007.g175.t1 -KPASSLQAI VGASQVKWNR VNRHVLATSH DGDVRIWDLR KGNTPVVYLT
aflo_s0310.g20.t1  -KPASSLQAI VGASQVKWNR VNRHVLATSH DGDVRIWDLR KGNTPVVYLT
XM_029340177.2     -KPASSLQAI VGASQVKWNR VNRHVLATSH DGDVRIWDLR KGNTPVIYLT
asel_s0045.g58.t2  -KPASSLQAI VGASQVKWNR VNRHVLATSH DGDVRIWDLR KGNTPVIYLT
ahya_s0003.g68.t2  -KPASSLQAI VGASQVKWNR VNRHVLATSH DGDVRIWDLR KGNTPVIYLT
acyt_s1402.g1.t1   TRGQSAVRTD SSQITGGPDI VNRHVLATSH DGDVRIWDLR KGNTPVIYLT
amur_s0006.g105.t2 -KPASSLQAI VGASQVKWNR VNRHVLATSH DGDVRIWDLR KGNTPVIYLT
ech_s0159.g30.t2   -KPASSLQAI VGASQVKWNR VNRHVLATSH DGDVRIWDLR KGNTPVIYLT
aacu_s0038.g69.t1  -KPASSLQAI VGASQVKWNR VNRHVLATSH DGDVRIWDLR KGNTPVIYLT
anas_s0109.g72.t2  -KPASSLQAI VGASQVKWNR VNRHVLATSH DGDVRIWDLR KGNTPVIYLT
amic_s0245.g8.t1   -KPASSLQAI VGASQVKWNR VNRHVLATSH DGDVRIWDLR KGNTPVIYLT
adig_s0048.g28.t1  -KPASSLQAI VGASQVKWNR VNRHVLATSH DGDVRIWDLR KGNTPVIYLT

aten_s0183.g20.t1  AHLSKIHGLD WSRSSGTTLA TCSSDTTVKL WNTEQPQQPE NKLNAKCPVW
ayon_s0004.g209.t1 ---------- ---------- ---------- ---------- ----------
aint_s0143.g6.t1   AHLSKIHGLD WSRSSGTTLA TCSSDTTVKL WNTEQPQQPE NKLNAKCPVW
agem_s0013.g175.t1 AHLSKIHGLD WSRSSGTTLA TCSSDTTVKL WNTEQPQQPE NKLNAKCPVW
aawi_s0007.g175.t1 AHLSKIHGLD WSRSSGTTLA TCSSDTTVKL WNTEQPQQPE NKLNAKCPVW
aflo_s0310.g20.t1  AHLSKIHGLD WSRSSGTTLA TCSSDTTVKL WNTEQPQQPE NKLNAKCPVW
XM_029340177.2     AHLSKIHGLD WSRSSGTTLA TCSSDTTVKL WNTEQPQQPE NKLNAKCPVW
asel_s0045.g58.t2  AHLSKIHGLD WSRSSGTTLA TCSSDTTVKL WNTEQPQQPE NKLNAKCPVW
ahya_s0003.g68.t2  AHLSKIHGLD WSCSSGTTLA TCSSDTTVKL WNTEQPQQPE NKLNAKCPVW
acyt_s1402.g1.t1   AHLSKIHGLD WSRSSGTTLA TCSSDTTVKL WNTEQPQQPE NKLNAKCPVW
amur_s0006.g105.t2 AHLSKIHGLD WSRSSGTTLA TCSSDTTVKL WNTEQPQQPE NKLNAKCPVW
ech_s0159.g30.t2   AHLSKIHGLD WSRSSGTTLA TCSSDTTVKL WNTEQPQQPE NKLNAKCPVW
aacu_s0038.g69.t1  AHLSKIHGLD WSRSSGTTLA TCSSDTTVKL WNTEQPQQPE NKLNAKCPVW
anas_s0109.g72.t2  AHLSKIHGLD WSHSSGTTLA TCSSDTTVKL WNIEQPQQPE NKLNAKCPVW
amic_s0245.g8.t1   AHLSKIHGLD WSRSSGTILA TCSSDTTVKL WNTEQPQQPE NKLNAKCPVW
adig_s0048.g28.t1  AHLSKIHGLD WSRSSGTTLA TCSSDTTVKL WNTEQPQQPE KKLNAKCPVW

aten_s0183.g20.t1  RARFTPFGEG LVTVTLPQLQ RGENSLSLWN IPDVNSPVAP PVNTFVGHSD
ayon_s0004.g209.t1 ---------- ---------- ---------- ---------- ----------
aint_s0143.g6.t1   RARFTPFGEG LVTVTLPQLQ RGENSLSLWN IPDVNSPVAA PVNTFVGHND
agem_s0013.g175.t1 RARFTPFGEG LVTVTLPQLQ RGENSLSLWN IPDVNSPVAA PVNTFVGHND
aawi_s0007.g175.t1 RARFTPFGEG LVTVTLPQLQ RGENSLSLWN IPDVNSPVAA PVNTFVGHND
aflo_s0310.g20.t1  RARFTPFGEG LVTVTLPQLQ RGENSLSLWN IPDVNSPVAA PVNTFVGHSD
XM_029340177.2     RARFTPFGEG LVTVTLPQLQ RGENNLSLWN IPDVNSPVAA PVNTFVGHTD
asel_s0045.g58.t2  RARFTPFGEG LVTVTLPQLQ RGENNLSLWN IPDVNSPVAA PVNTFVGHTD
ahya_s0003.g68.t2  RARFTPFGEG LVTVTLPQLQ RGENSLSLWN IPDVNSPVAA PVNTFVGHTD
acyt_s1402.g1.t1   RARFTPFGEG LVTVTLPQLQ RGENSLSLWN IPDVNSPVAA PVNTFVGHTD
amur_s0006.g105.t2 RARFTPFGEG LVTVTLPQLQ RGENSLSLWN IPDVNSPVAA PVNTFVGHTD
ech_s0159.g30.t2   RARFTPFGEG LVTVTLPQLQ RGENSLSLWN IPDVNSPVAA PVNTFVGHTD
aacu_s0038.g69.t1  RARFTPFGEG LVTVTLPQLQ RGENSLSLWN IPDVNSPVAA PVNTFVGHSD
anas_s0109.g72.t2  RARFTPFGEG LVTVTLPQLQ RGENSLSLWN IPDVNSPVAA PVNTFVGHTD
amic_s0245.g8.t1   RARFTPFGEG LVTVTLPQLQ RGENNLSLWN IPDVDSPVAA PVNTFVGHTD
adig_s0048.g28.t1  RARFTPFGEG LVTVTLPQLQ QGENYLSLWN IPDVNSPVAA PVNTFVGHSD

aten_s0183.g20.t1  VVLDFHWRSQ TLDRDEQFQL ITWAKDCCLR LWVLEPRMIM ACSGDTSTNF
ayon_s0004.g209.t1 ---------- ---------- ---------- -------MIM ACSGDTSTNF
aint_s0143.g6.t1   VVLDFHWRSQ TLDNDEQFQL ITWAKDCCLR LWVLEPRMIM ACSGDTSTNF
agem_s0013.g175.t1 VVLDFHWRSQ TLDKDEQFQL ITWAKDCCLR LWVLEPRMIM ACNGDTSTNF
aawi_s0007.g175.t1 VVLDFHWRSQ TLDKDEQFQL ITWAKDCCLR LWVLEPRMIM ACSGDTSTNF
aflo_s0310.g20.t1  VVLDFHWRSQ TLDKDEQFQL ITWAKDCCLR LWVLEPRMIM ACSGDTSTNF
XM_029340177.2     VVLDFHWRSQ TLDKDEQFQL ITWAKDCCLR LWVLEPRMIM ACSGDTSTNF
asel_s0045.g58.t2  VVLDFHWRSQ TLDKDEQFQL ITWAKDCCLR LWVLEPRMIM ACSGDTSTNF
ahya_s0003.g68.t2  VVLDFHWRSQ TLYEDEQFQL ITWGKDCCLR LWVLEPRMIM ACSGDTSTNF
acyt_s1402.g1.t1   VVLDFHWRSQ TLYEDEQFQL ITWGKDCCLR LWVLEPRMIM ACSGDTSTNF
amur_s0006.g105.t2 VVLDFHWRSQ TLDKDEQFQL ITWGKDCCLR LWVLEPRMIM ACSGDTSTNF
ech_s0159.g30.t2   VVLDFHWRSQ TLDKDEQFQL ITWGKDCCLR LWVLEPRMIM ACSGDTSTNF
aacu_s0038.g69.t1  VVLDFHWRSQ TLDKDEQFQL ITWAKDCCLR LWVLEPRMIM ACSGDTSTNF
anas_s0109.g72.t2  VVLDFHWRSQ TLDKDEQFQL ITWAKDCCLR LWVLEPRMIM ACSGDTSTNF
amic_s0245.g8.t1   VVLDFHWRSQ TLDKDEQFQL ITWAKDCCLR LWVLEPRMIM ACSGDTSTNF
adig_s0048.g28.t1  VVLDFHWRSQ TLDEDEQFQL ITWAKDCCLR LWVLEPRMIM ACSGDTSTNF
```

### Fig S2 continued

```
aten_s0183.g20.t1  PSRSTSETAH SMSDELIEMA IAESVIIDTP KSFPDPAAQA QTLLQEFALI
ayon_s0004.g209.t1 PSRSTSETAH SMSDELIEMA IAESVIIDTP KSFPDPAAQP QTLLQEFALI
aint_s0143.g6.t1   PSRSTSETAH SMSDEMIEMA IAESVIIDTP KSFPDLAAQP QTLLQEFSLI
agem_s0013.g175.t1 PSPSTSETAH SMSDEMIEMA IAE------- ---------- ----------
aawi_s0007.g175.t1 PSRSTSETAH SMSDEMIEMA IAESVIIDTP KSFPDLAAQP QTLLQEFSLI
aflo_s0310.g20.t1  PSRSTSETAH SMSDEMIEMA IAESVIIDTP KSFPDLAAQP QTLLQEFSLI
XM_029340177.2     PSRSTNETAH SMSEELIEMA IAESVIIDTP KSFPDLAAQP QTLQQEFALI
asel_s0045.g58.t2  PSRSTNETAH SMSEELIEMA IAESVIIDTP KSFPDLAAQP QTLQQEFALI
ahya_s0003.g68.t2  PSHSTNETAH SMSEELIEMA IAESVIIDTP KSFPDLAAQP QTLQQEFALI
acyt_s1402.g1.t1   PSHSTNETAH SMSEELIEMA IAESVIIDTP KSFPDLAAQP QTLQQEFALI
amur_s0006.g105.t2 PSRSTNETAH SMSEELIEMA IAESVIIDTP KSFPDLAAQP QTLQQEFALI
ech_s0159.g30.t2   PSRSTNETAH SMSEELIEMA IAESVIIDTP KSFPDLAAQP QTLQQEFALI
aacu_s0038.g69.t1  PSRTTNETAH SMSEELIETA IAESVIIDTP KSFPDLAAQP QTLQQEFALI
anas_s0109.g72.t2  PSRSTNETAH SMSEEQ---- IAESVKIDTP KSFPDLAAQP QTLQQEFALI
amic_s0245.g8.t1   PSHSTNETAH SMSEELIEMT IAESVIIDTP KSFPDHAAQP QTLQQEFALI
adig_s0048.g28.t1  PSRSTNETAH SMSEELIKMA IAES------ ---PDLAAQP QTLQQEFALI

aten_s0183.g20.t1  NVNIPNVTVE QLDAGHRSCT VCATNGPNTV FLVINFPSLY PNKAMPSFEF
ayon_s0004.g209.t1 NVNIPNVTVE QLDAGHRSCT VCATNGPNTV SLVINFPSLY PNKAMPSFEF
aint_s0143.g6.t1   NVNIPNVIVE QLDAGHRSCT VCATNGPDVV SLVINFPSLY PNQAIPSFEF
agem_s0013.g175.t1 ---------- SLDAGHRSCT VCATNGPDVV SLVINFPSLY PNQAIPSFEF
aawi_s0007.g175.t1 NVNIPNVIVE QLDAGHRSCT VCATNGPDVV SLVINFPSLY PNQAIPSFEF
aflo_s0310.g20.t1  NVNIPNVIVE QLDAGHRSCT VCATNGPDIV SLVINFPSLY PNQAIPSFEF
XM_029340177.2     NVNIPNVTVE QLDAGHRSCT VCATSGPDTV SLVINFPSLY PNQAIPSFEF
asel_s0045.g58.t2  NVNIPNVTVE QLDAGHRSCT VCATSGPDTV SLVINFPSLY PNQAIPSFEF
ahya_s0003.g68.t2  NVNIPNVTVE QLDAGHRSCT VCAKNGPDTV SLVINFPSLY PNQAIPSFEF
acyt_s1402.g1.t1   NVNIPNVTVE QLDAGHRSCT VCATNGPNTV SLVINFPSLY PNQAIPSFEF
amur_s0006.g105.t2 NVNIPNVTVE QLDAGHRSCT ACATNGPDTV SLVINFPSLY PNQAIPSFEF
ech_s0159.g30.t2   NVNIPNVTVE QLDAGHRSCT VCATNGPDTV SLVINFPSLY PNQAIPSFEF
aacu_s0038.g69.t1  NVNIPNVTVE QLDAGHRSCT VCATNGPDTV SLVINFPSLY PNQAIPSFEF
anas_s0109.g72.t2  NVNIPNVTVE QLDAGQRSCT VCATIGPDTV SLVINFPSLY PNQAIPSFEF
amic_s0245.g8.t1   NVNIPNVTVE QLDAGHRSCT VCATIGPDTV SLVINFPSLY PNQAIPSFEF
adig_s0048.g28.t1  NVNIPNVTVE QLDAGHRSCT VCATSGPDTV SLVINFPSLY PNQAIPSFEF

aten_s0183.g20.t1  TADTTIDTNT KTKLMKTLRE TAQTHVKLNQ TCLEPCLRQL VNHLDEELTL
ayon_s0004.g209.t1 TADTTIDTNT KTKLMKTLRE TAQTHVKLNQ TCLEPCLRQL VNHLDEELTL
aint_s0143.g6.t1   TADTSIDTNT KTKLMKTLRE TAQTHVKLNQ TCLEPCLRQL VNHLE-ELTL
agem_s0013.g175.t1 TADTSIDTNT KTKLMKTLRE TAQTHVKLNQ TCLEPCLRQL VNHLE-ELTL
aawi_s0007.g175.t1 TADTSIDTNT KTKLMKTLRE TAQTHVKLNQ TCLEPCLRQL VNHLE-ELTL
aflo_s0310.g20.t1  TADTSIDTNT KTKLMKTLRE TAQTHVKLNQ TCLEPCLRQL VNHLE-ELTL
XM_029340177.2     TADTSIDTNT KTKLMKTLRE TAQTHVKLNQ TCLEPCLRQL VSHLE-ELTL
asel_s0045.g58.t2  TADTSIDTNT KTKLMKTLRE TAQTHVKLNQ TCLEPCLRQL VSHLE-ELTL
ahya_s0003.g68.t2  TADTSVDTNT KTKLMKTLRE TAQTHVKLNQ TCLEPCLRQL VSHLE-ELTL
acyt_s1402.g1.t1   TADTSVDTNT KTKLMK---- ---------- ---------- ---------L
amur_s0006.g105.t2 TADTSVDTNT KTKLMKTLRE TAQTHVKLNQ TCLEPCLRQL VSHLE-ELTL
ech_s0159.g30.t2   TADTSIDTNT KTKLMKTLRE TAQTHVKLNQ TCLEPCLRQL VSHLE-ELTL
aacu_s0038.g69.t1  TADTSIDTNT KTKLMKTLRE TAQTHVKLNQ TCLEPCLRQL VSHLE-ELTL
anas_s0109.g72.t2  TADTSIDTNT KTKLMKTLRE TAQTHVKLNQ TCLEPCLRQL VSHLE-ELTL
amic_s0245.g8.t1   TADTSIDTNT KTKLMKTLRE TAQTHVKLNQ TCLEPCLRQL VSHLE-ELTL
adig_s0048.g28.t1  TADTSIDTNT KTKLMKTLCE TAQTHVKLNQ TCLEPCLRQL VSHLE-ELTL

aten_s0183.g20.t1  QERLPMDQYA ATVPQPHGRL DPISYGSYQD AAVPFPRTSG AKFCACGLLV
ayon_s0004.g209.t1 QERLPMDQFA ATVPQPHGRL DPISYGSYQD AAVPFPRTSG AKFCACGLLV
aint_s0143.g6.t1   QERLPMDQFA ATVPQLHGRL DPISYGSYAD AAVPFPRTSG AKFCACGLLV
agem_s0013.g175.t1 QERLPMDQFA ATVPQLHGRL DPISYGSYAD AAVPFPRTSG AKFCACGLLV
aawi_s0007.g175.t1 QERLPMDQFA ATVPQLHGRL DPISYGSYAD AAVPFPRTSG AKFCACGLLV
aflo_s0310.g20.t1  QERLPMDQFA ATVPQLHGRL DPISYGSYAD AAVPFPRTSG AKFCACGLLV
XM_029340177.2     QERLPMDQFA ATVPQLHGRL DSISYGSYAD AAVPFPRTSG AKFCACGLLV
asel_s0045.g58.t2  QERLPMDQFA ATVPQLHGRL DSISYGSYAD AAVPFPRTSG AKFCACGLLV
ahya_s0003.g68.t2  QERLPMDQFA ATAPQLHGRL DPVSYGSYLD AAIPFPRTSG AKFCACGLLV
acyt_s1402.g1.t1   QERLPMDQFA ATAPQLHGRL DPVSYGSYLD AAIPFPRTSG AKFCACGLLV
amur_s0006.g105.t2 QERLPMDQFA ATAPQLPL-- ------NFLD AAIPFPRTSG AKFCACGLLV
ech_s0159.g30.t2   QERLPMDQFA ATVPQLHGRL DPISYGSYAD AAVPFPRTSG AKFCACGLLV
aacu_s0038.g69.t1  QERLPMDQFA ATAPQLHDRL DPISYVSFLD AAIPFPRTSG AKFCACGLLV
anas_s0109.g72.t2  QERLPMDQFA ATAPQLHGQL DPISNASFLD AAIPFPRTSG AKFCACGLLV
amic_s0245.g8.t1   QERLPMDQFA ATAPQLHDRL DPIPYVSFLD AAIPFPRTSG AKFCACGLLV
adig_s0048.g28.t1  QERLPMDQFA ATAPQLHDRL DPISYVSFLD AAIPFPRTSG AKFCACGLLV
```

### Fig S2 continued

```
aten_s0183.g20.t1  CFNLPGRYGG RVGSGGEPTP RSLSAFSAYS SRPSSGPALP PLINRHFPKT
ayon_s0004.g209.t1 CFNLPGRYGG RVGSGGEPTP RSLSAFSAYS SRPSSGPALP PLINRHFAKT
aint_s0143.g6.t1   CFNLPGRYGG RVGSGGEPTP RSLSAFSAYS SRPSSGPALP PLINRHFPKT
agem_s0013.g175.t1 CFNLPGRYGG RVGSGGEPTP RSLSAFSAYS SRPSSGPALP PLINRHFQKN
aawi_s0007.g175.t1 CFNLPGRYGG RVGSGGEPTP RSLSAFSAYS SRPSSGPALP PLINRHFQKN
aflo_s0310.g20.t1  CFNLPGRYGG RVGSGGEPTP RSLSAFSAYS SRPSSGPALP PLINRHFQKN
XM_029340177.2     CFNLPGRYGG RVGSGGEPTP RSLSAFSAYS SRPSSGPALP RLIDRHFQNS
asel_s0045.g58.t2  CFNLPGRYGG RVGSGGEPTP RSLSAFSAYS SRPSSGPALP RLVDRHFQNS
ahya_s0003.g68.t2  CFNLPGRYGG RVGSGGEPTP RSLSAFSAYS SRPSSGPALP RLINRHFQNP
acyt_s1402.g1.t1   CFNLPGRYGG RVGSGGEPTP RSLSAFSAYS SRPSSGPALP RLINRHFQNP
amur_s0006.g105.t2 CFNLPGRYGG RVGSGGEPTP RSLSAFSAFS FRPSSGPALP RLINRHFQKP
ech_s0159.g30.t2   CFNLPGRYGG RVGSGGEPTP RSLSAFSAYS SRPSSGPALP RLINRHFQNP
aacu_s0038.g69.t1  CFNLPGRYGG RVGSGGEPTP RSLSAFSAYS SRPSSGPALP RLINRHFQKP
anas_s0109.g72.t2  CFNLPGRYSG RVGSGGEPTP RSLSAFSAYS SRPSSGPALP RLINRHFQKP
amic_s0245.g8.t1   CFNLPGRYGG RVGSGGEPTP RSLSAFSAYS SRPSSGPALP RLINRHFQKP
adig_s0048.g28.t1  CFNLPGRYGG RVGSGGEPTP RSLSAFSAYS SRPSSGPALP RLINGHFQKP

aten_s0183.g20.t1  HQRHAFGPFG QQARTFTYRA SSKTEDPQDR QKYSGVPSRN SDVGLVVIRD
ayon_s0004.g209.t1 HQRHAFGPFG QQARTFTYRA LSKTEDPQDR QKYSGVPSRN SDVGLVVIRD
aint_s0143.g6.t1   RQRHAFGSFG QQTRTFTYRA LSKTEDPQDR QKYSGVPSRN SDVGLVVIRD
agem_s0013.g175.t1 RQRHAFGSFG QQTRSFTYRA LSKTEDPQDR QKYSGVPSRN SDVGLVVIRD
aawi_s0007.g175.t1 RQRHAFGSFG QQTRSFTYRA LSKTEDPQDR QKYSGVPSRN SDVGLVVIRD
aflo_s0310.g20.t1  RQRHAFGSFG QQSRTFTYRA LSKTEDPQDR QKYSGVPSRN SDVGLVVIRD
XM_029340177.2     RERHALGSFD QQTRTFTYRA LSRTEDPQDK QKYSGLPSRT SDVGLVVIRD
asel_s0045.g58.t2  RERHALGSFD QQTRTFTYRA LSRTEDPQDR QKYSGLPSRT SDVGLVVIRD
ahya_s0003.g68.t2  RERHALGSFD QQTRTFTYRA LSRTEDPQDR QKYSGLPSRT SVVGLVVIRD
acyt_s1402.g1.t1   RERHALGSFD QQTRTFTYRA LSRVGEL*-- ---------- ----------
amur_s0006.g105.t2 RERHALGSFG QQTRTFTYRA LSRTEDPQDR QKYSGLPSRT SDVGLVVIRD
ech_s0159.g30.t2   RERHALGSFD QQTRTFTYRA LSRTDDPQDR QKYSGLPSRT SVVGLVVIRD
aacu_s0038.g69.t1  LQRHAFGSFG QLTRTFTYRA LSRTEDPQDR QKYSGLPSRN SDVGLVVIRD
anas_s0109.g72.t2  LQRHGFGSFG QLTRTFTYRA LSRTEDPQDR QKYSGFPSRN SDVGLVVIRD
amic_s0245.g8.t1   LQRHAFGSFG QLTRTFTYRA LSRTEDPQDR QKYSGFPSRN SDVGLVVIRD
adig_s0048.g28.t1  LQRHTFGSFR QLTRTFTYRA LSRTEDPQDR QKYSGLPSRK SDVGLVVIRD

aten_s0183.g20.t1  VSAMMPIHQS LGQAYTLEGN NITKICKKNL SAAMTTGRKD LVQTWSLLSL
ayon_s0004.g209.t1 VSAMMPIHQS LGQAYTLEGN NITKICKKNL SAAMTTGRKD LVQTWSLLSL
aint_s0143.g6.t1   VSAMMPIHQS LGQAYTLEGN NITKICKKNL SAAMTTGRKD LVQTWSLLSL
agem_s0013.g175.t1 VSAMMPIHQS LGQAYTLEGN NITKICKKNL SAAMTTGRKD LVQTWSLLSL
aawi_s0007.g175.t1 VSAMMPIHQS LGQAYTLEGN NITKICKKNL SAAMTTGRKD LVQTWSLLSL
aflo_s0310.g20.t1  VSAMMPIHQS LGQAYTLEGN NITKICKKNL SAAMTTGRKD LVQTWSLLSL
XM_029340177.2     VSAMMPIHQS LGQAYTLEGN NITKICKKNL SAAMTTGRKD LVQTWSLLSL
asel_s0045.g58.t2  VSAMMPIHQS LGQAYTLEGN NITKICKKNL SAAMTTGRKD LVQTWSLLSL
ahya_s0003.g68.t2  VSAMMPIHQS LGQAYTLEGN NITKICKKNL SAAMTTGRKD LVQTWSLLSL
acyt_s1402.g1.t1   ---------- ---------- ---------- ---------- ----------
amur_s0006.g105.t2 VSAMMPIHQS LGQAYTLEGN NITKICKKNL SAAMTTGRKD LVQTWSLLSL
ech_s0159.g30.t2   VSAMMPIHQS LGQAYTLEGN NITKICKKNL SAAMTTGRKD LVQTWSLLSL
aacu_s0038.g69.t1  VSAMMPIHQS LGQAYTLEGN NITKICKKNL SAAMTTGRKD LVQTWSLLSL
anas_s0109.g72.t2  VSAMMPIHQS LGQAYTLEGN NITKICKKNL SAAMKTGRKD LVQTWSLLSL
amic_s0245.g8.t1   VSAMMPIHQS LGQAYTLEGN NITKICKKNL SAAMTTGRKD LVQTWSLLSL
adig_s0048.g28.t1  VSAMMPIHQS LGQAYTLEGN NITKICKKNL SAAMTTGRKD LVQTWSLLSL

aten_s0183.g20.t1  VLDEKLAPS- -------DSM DEAPWALHPF GKKLVNSLMD YYLNIRDIQT
ayon_s0004.g209.t1 VLDEKLAPS- -------DSM DEAPWALHPF GKKLVNSLMD YYLNIRDIQT
aint_s0143.g6.t1   VLDEKLAPS- -------DSM DEAPWALHPF GKKLVSSLMD YYLNIRDIQT
agem_s0013.g175.t1 VLDEKLAPS- -------DSM DEAPWALHPF GKKLVSSLMD YYLNIRDIQT
aawi_s0007.g175.t1 VLDEKLAPS- -------DSM DEAPWALHPF GKKLVSSLMD YYLNIRDIQT
aflo_s0310.g20.t1  VLDEKLAPS- -------DSM DEAPWALHPF GKKLVSSLMD YYLNIRDIQT
XM_029340177.2     VLDEKLASS- -------DSM DEAPWALHPF GKKLVSSLMD YYLNIRDIQT
asel_s0045.g58.t2  VLDEKLASS- -------DSM DEAPWALHPF GKKLVSSLMD YYLNIRDIQT
ahya_s0003.g68.t2  VLDEKLASS- -------DSM DQAPWALHPF GKKLVSSLMD YYLNIRDIQT
acyt_s1402.g1.t1   ---------- ---------- ---------- ---------- ----------
amur_s0006.g105.t2 VLDEKLASS- -------DSM DEAPWALHPF GKKLVSSLMD YYLNIRDIQT
ech_s0159.g30.t2   VLDEKLASS- -------DSM DEAPWALHPF GKKLVSSLMD YYLNIRDIQT
aacu_s0038.g69.t1  VLDEKLAPSD SIDEAPSDSI DEAPWALHPF GKKLVSSLMD YYLNIRDIQT
anas_s0109.g72.t2  VLDEKLAPSD PIDEAPSDSI DEAPWALHPF GKKLVSSLMD YYLNIRDIQT
amic_s0245.g8.t1   VLDEKLAHS- -------NSM DEAPWALHPF GKKLVSSLME YYLNIRDIQT
adig_s0048.g28.t1  VLDEKLAPS- -------DSI DEAPWALHPF GKKLVSSLMD YYLNIRDIQT
```

### Fig S2 continued

```
                                             Pro747Ser
aten_s0183.g20.t1  LGMLSCVLAH HALSDAFKPP RNSFNTEVIP SSMSFPFGSP PSFNNDSPSQ
ayon_s0004.g209.t1 LGMLSCVLAH HALSDDFKPP RNSFNTEVIP SSMSFPFGSP PSFNNDSPSQ
aint_s0143.g6.t1   LGMLSCVLAH HALSDAFKPP RNSFNTEVIS SSMSFPFGSP PSFNNDTPSQ
agem_s0013.g175.t1 LGMLSCVLAH HALSDAFKPP RNSFNTEVIP SSMSFPFGSP PSFNNDTPSQ
aawi_s0007.g175.t1 LGMLSCVLAH HALSDAFKPP RNSFNTEVIP SSMSFPFGSP PSFNNDTPSQ
aflo_s0310.g20.t1  LGMLSCVLAH HALSDAFKPP RNSFNTEVIS SSMSFPFGSP PSFNNDTPSQ
XM_029340177.2     LGMLSCVLAH HALSDAFKPP RNSFKTGVIS SSMSFPFGSP PSFNNDTPSQ
asel_s0045.g58.t2  LGMLSCVLAH HALSDAFKPP RNSFKTGVIS SSMSFPFGSP PSFNNDTPSQ
ahya_s0003.g68.t2  LGMLSCVLAH HALSDAFKPP RNSFYTEVIS SSMSFPFGSP PSFNNDTPSQ
acyt_s1402.g1.t1   ---------- ---------- ---------- ---------- ----------
amur_s0006.g105.t2 LGMLSCVLAH HALSDAFKPP RNSFNTEVIS SSMSFPFGSP PSFNNDTPSQ
ech_s0159.g30.t2   LGMLSCVLAH HALSDAFKPP RNSFNTEVIS SSMSFPFGSP PSFNNDTPSQ
aacu_s0038.g69.t1  LGMLSCVLAH HALSDAFKPL RNSFNTEVIP SSMSFHFGSP SSFNNDTPAQ
anas_s0109.g72.t2  LGMLSCVLAH HALSDAFKPH RDSFNTEEIS SS-------- TSFNNDTPSQ
amic_s0245.g8.t1   LGMLSCVLAH HALSDAFKPL RNSFNTEVIP SSMSFPFGSP QSFNNDTPLQ
adig_s0048.g28.t1  LGMLSCVLAH HALSDAFKPP RNSFNTEVIS SSMSFPFGSP SSFNNDTPSQ

aten_s0183.g20.t1  SVKRPKPVGS VTDTMVTESS PSE--WSE-- VSFPLSGGLS PPVFIRSKDP
ayon_s0004.g209.t1 SVKRPKPVGS VTDTMVTESS PSE--WSE-- VSFPLSGGLS PPVFIRSKDP
aint_s0143.g6.t1   SVKRPKPVGS VPDPMATESS PSE--WSE-- VSFPLSGGLS PTVFIRSKDP
agem_s0013.g175.t1 SVKRPKPVGS VPDPMATESS PSE--WSE-- VSFPLSGGLS PTVFIRSKDP
aawi_s0007.g175.t1 SVKRPKPVGS VPDPMATESS PSE--WSE-- VSFPLSGGLS PTVFIRSKDP
aflo_s0310.g20.t1  SVKRPKPVGS VPDPMATESS PSE--WSE-- VSFPLSGGLS PTVFIRSKDP
XM_029340177.2     SVKRPKPVGS VSDPMVNESS PSE--WSE-- VTFPLSGGLS PTVFIRSKDP
asel_s0045.g58.t2  SVKRPKPVGS VPDPMGNESS PSE--WSE-- VTFPLSGGLS PTVFIRSKDP
ahya_s0003.g68.t2  SVKRPKPVGS VPDPMGNESS PSE--WSE-- VSFPLSGGLS PTVFIRSKDP
acyt_s1402.g1.t1   ---------- ---------- ---------- ---------- ----------
amur_s0006.g105.t2 SVKRPKPVGS VPDPMVTESS SSE--WSE-- VSFPLSGGLS PTVFIRSKDP
ech_s0159.g30.t2   SVKRPKPVGS VPDPMGNESS PSE--WSE-- VSFPLSGGLS PTVFIRSKDP
aacu_s0038.g69.t1  SAKRPKPVGS LPDPMVTESS PIRIQWKNKE SSFKVTADLS TPR-------
anas_s0109.g72.t2  SVKRPKPVGS VLVPMVTESS PSE--WSE-- VSFPLSGGLS PTVSIRSKDP
amic_s0245.g8.t1   SVKRPKPVGS VPDPTGNESS PSE--WSE-- LSFPLSGGLS PTVFIRSKDP
adig_s0048.g28.t1  SVKRPKPVGS VPDPMVTESS PSE--WSE-- LSFPLSGGLS PTVFIRSKDP

aten_s0183.g20.t1  KEEQRKQYQS DCRLVDPSQV KLHDHYIKQY ADVLYRWSLL GKRAEVTKFL
ayon_s0004.g209.t1 KEEQRKQYQS DCRLVDPSQV KLHDHYIKQY ADVLYRWSLL GKRAEVTKFL
aint_s0143.g6.t1   MEEQRKQYQS DCRFVDPSQV KLHDHYIKQY ADVLYRWSLL GKRAEVTKFL
agem_s0013.g175.t1 MEEQRKQYQS DCRFVDPSQV KLHDHYIKQY ADVLYRWSLL GKRAEVTKFL
aawi_s0007.g175.t1 MEEQRKQYQS DCRFVDPSQV KLHDHYIKQY ADVLYRWSLL GKRAEVTKFL
aflo_s0310.g20.t1  MEEQRKQYQS DCRFVDPSQV KLHDHYIRQY ADVLYRWSLL GKRAEVTKFL
XM_029340177.2     MEEQRKQFQS DCRFVDPSQV KLHDHYIKQY ADVLYRWSLL GKRAEVTKFL
asel_s0045.g58.t2  MEEQRKQFQS DCR*------ ---------- ---------- ----------
ahya_s0003.g68.t2  MEEQRKQFES DCR*------ ---------- ---------- ----------
acyt_s1402.g1.t1   ---------- ---------- ---------- ---------- ----------
amur_s0006.g105.t2 MEEQRKQFQS DCR*------ ---------- ---------- ----------
ech_s0159.g30.t2   MEEQRKQFQS DCR*------ ---------- ---------- ----------
aacu_s0038.g69.t1  --------KS NCMTIT*--- ---------- ---------- ----------
anas_s0109.g72.t2  MEEQKKQFKS DCR*------ ---------- ---------- ----------
amic_s0245.g8.t1   MEEQKKQFKS DSR*------ ---------- ---------- ----------
adig_s0048.g28.t1  MEEQRKQFKS DCRFVDPSQV KLHDHYIKQY ADVLYRWSLL GKRAEVTKFL

aten_s0183.g20.t1  SETQVPHSGA EFSTRCYNCS RNLRGAQCGS CKSFGLQCVI CHVAVRGASN
ayon_s0004.g209.t1 SETQLPHSGA EFSTRCYNCS RNFRGAQCGS CKSFGLQCVI CHVAVRGASN
aint_s0143.g6.t1   SQPQLPHSGA EFSTRCYNCS RNLRGAQCGS CKSFGLQCVI CHVAVRGASN
agem_s0013.g175.t1 SEPQLPHSGA EFSTRCYNCS RNLRGAQCGS CKSFGLQCVI CHVAVRGASN
aawi_s0007.g175.t1 SEPQLPHSGA EFSTRCYNCS RNLRGAQCGS CKSFGLQCVI CHVAVRGASN
aflo_s0310.g20.t1  SEPQLPHSGA EFSTRCYNCS RNLRGAQCGS CKSFGLQCVI CHVAVRGASN
XM_029340177.2     SEPQLPHSGA EFSTRCYNCS RNLRGAQCGS CKSFGLQCVI CHVAVRGASN
asel_s0045.g58.t2  ---------- ---------- ---------- ---------- ----------
ahya_s0003.g68.t2  ---------- ---------- ---------- ---------- ----------
acyt_s1402.g1.t1   ---------- ---------- ---------- ---------- ----------
amur_s0006.g105.t2 ---------- ---------- ---------- ---------- ----------
ech_s0159.g30.t2   ---------- ---------- ---------- ---------- ----------
aacu_s0038.g69.t1  ---------- ---------- ---------- ---------- ----------
anas_s0109.g72.t2  ---------- ---------- ---------- ---------- ----------
amic_s0245.g8.t1   ---------- ---------- ---------- ---------- ----------
adig_s0048.g28.t1  SEPQLPHSGA EFSTRCYNCS RNLRGAQCGS CKSFGLQCVI CHVAVRGASN
```

### Fig S2 continued

```
                                          Glu942Lys             Thr962Ile
aten_s0183.g20.t1  FCVACGHGGH AYHLLTWFES MNVCPTGCGC RCLEVDT-FI VD*
ayon_s0004.g209.t1 FCVACGHGGH AYHLLTWFES MNVCPTGCGC RCLEVDT-FI VD*
aint_s0143.g6.t1   FCVACGHGGH AYHLLTWFES MDVCPTGCGC RCLEVGT-FI VD*
agem_s0013.g175.t1 FCVACGHGGH AYHLLTWFES MDVCPTGCGC RCLEVGT-FI VD*
aawi_s0007.g175.t1 FCVACGHGGH AYHLLTWFES MDVCPTGCGC RCLEVGT-FI VD*
aflo_s0310.g20.t1  FCVACGHGGH AYHLLTWFES MDVCPTGCGC RCLEVGT-FI VD*
XM_029340177.2     FCVACGHGGH AYHLLTWFES MDVCPTGCGC RCREVGVTFI VD*
asel_s0045.g58.t2  ---------- ---------- ---------- ---------- ---
ahya_s0003.g68.t2  ---------- ---------- ---------- ---------- ---
acyt_s1402.g1.t1   ---------- ---------- ---------- ---------- ---
amur_s0006.g105.t2 ---------- ---------- ---------- ---------- ---
ech_s0159.g30.t2   ---------- ---------- ---------- ---------- ---
aacu_s0038.g69.t1  ---------- ---------- ---------- ---------- ---
anas_s0109.g72.t2  ---------- ---------- ---------- ---------- ---
amic_s0245.g8.t1   ---------- ---------- ---------- ---------- ---
adig_s0048.g28.t1  FCVACGHGGH AYHLLTWFES MDVCPTGCGC RCREVGTFTV D*-
```

**Figure S2.** Alignment of WDR59 amino acid sequences from 16 *Acropora* corals. Three amino acid sites that differ between *A. digitifera* and *Acropora* sp.1 are shown in blue.

Fig S3

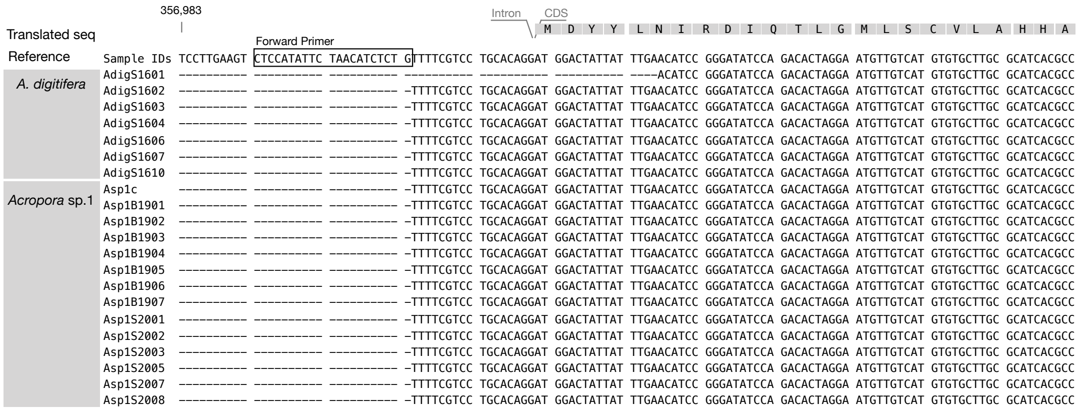


Fig S3 continued

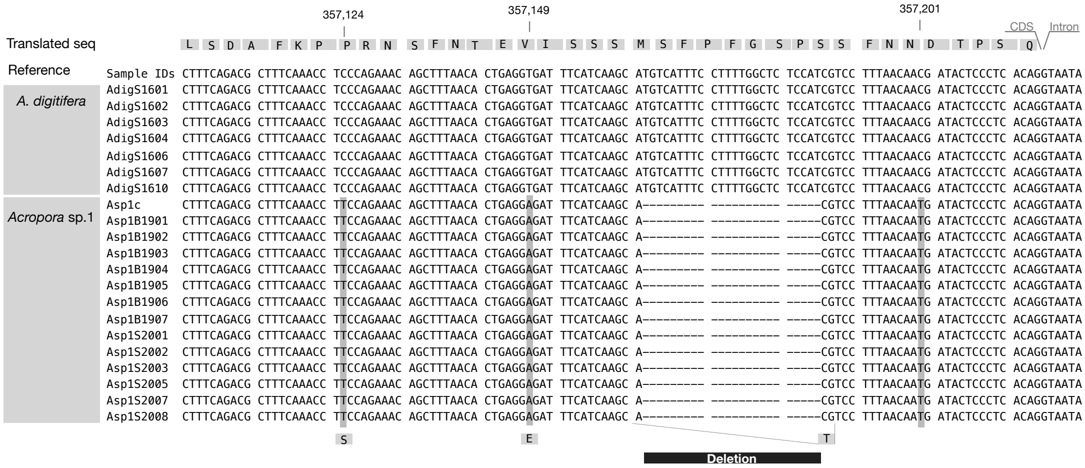


Fig S3 continued

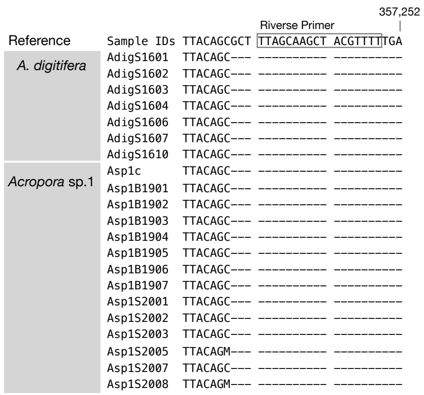

**Figure S3.** Alignment of some *WDR59* sequences from *A. digitifera* and *Acropora* sp. 1 with the *A. digitifera* reference genome. Non-synonymous sites are shown in gray. The deletion is shown with a black line. Primer sequence locations are indicated by squares on the reference genome.

Fig S4

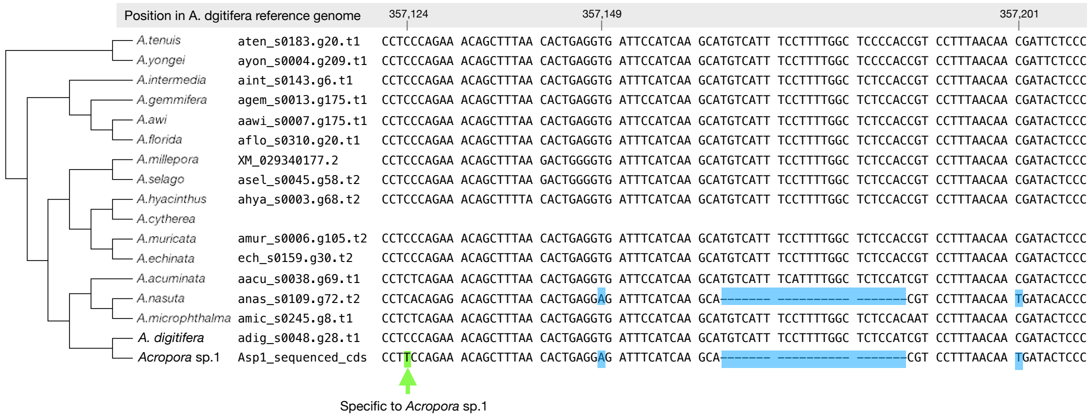


**Figure S4.** A schematic diagram of the alignment of some *WDR59* sequences from 16 *Acropora* corals. Mutations shared with *Acropora* sp. 1 and *A. nasuta* are shown in blue. The mutation specific to *Acropora* sp. 1 is indicated by arrows.

### Fig S5

```
aten_s0183.g20.t1  ATGTTGACTG ACGACGATAT CCATTCTGAA ACCAAGATGG CGTCTTACCA
ayon_s0004.g209.t1 ---------- ---------- ---------- ---------- ----------
aint_s0143.g6.t1   ---------- ---------- ---------- ------ATGG CGTCTTACCA
agem_s0013.g175.t1 ---------- ---------- ---------- ------ATGG CGTCTTACCA
aawi_s0007.g175.t1 ---------- ---------- ---------- ------ATGG CGTCTTACCA
aflo_s0310.g20.t1  ---ATGTTGA CTGACGATAT CCATCCTGAA ACCAAGATGG CGTCTTACCA
XM_029340177.2     ---ATGTTGA CTGACGATAT CCCTCCTGAA ACCAAGATGG CGTCTTACCA
asel_s0045.g58.t2  ---ATGTTGA CTGACGATAT CCCTCCTGAA ACCAAGATGG CGTCTTACCA
ahya_s0003.g68.t2  ---------- ---------- ---------- ------ATGG CGTCTTACCA
acyt_s1402.g1.t1   GTTATTTGCA ATGCTCTTAG ATTGTGGGAC ATTAAGCTAG TGGCATTTCC
amur_s0006.g105.t2 ---------- ---------- ---------- ------ATGG CGTCTTACCA
ech_s0159.g30.t2   ---------- ---------- ---------- ------ATGG CGTCTTACCA
aacu_s0038.g69.t1  ---ATGTTGA CTGACGATAT CCCTCCTGAA ACCAAGATGG CGTCTTACCA
anas_s0109.g72.t2  ---ATGTTGA CTGACGATAT CCCTCCTGAA ACCAAGATGG CGTCTTACCA
amic_s0245.g8.t1   ---ATGTTGA CTGACGATAT CCATCCTGAA ACCAAGATGG CGTCTTACCA
adig_s0048.g28.t1  ---ATGTTGA CTGACGATAT CCATCCTGAA ACCAAGATGG CGTCTTACCA
Asp1_sequenced_cds ---------- ---------- ---------- ---------- ----------

aten_s0183.g20.t1  GGACGCAGTT GCACCGCAC- --AAAGACTG GCCTGCAACA GTCATGGCTG
ayon_s0004.g209.t1 ---------- ---------- ---------- ---------- ----------
aint_s0143.g6.t1   GGATGCAGTT GCACCGCAC- --AAAGACTG GCCTGCAACA GTCATGGCTG
agem_s0013.g175.t1 GGACGCAGTT GCACCGCAC- --AAAGACTG GCCTGCAACA GTCATGGCTG
aawi_s0007.g175.t1 GGACGCAGTT GCACCGCAC- --AAAGACTG GCCTGCAACA GTCATGGCTG
aflo_s0310.g20.t1  GGACGCAGTT GCAGCGCAC- --AAAGACTG GCCTGCAACA GTCATGGCTG
XM_029340177.2     GGACGCAGTT GCAGCGCAC- -----GACTG GCCTGCAACA GTCATGGCTG
asel_s0045.g58.t2  GGACGCAGTT GCAGCGCAC- -----GACTG GCCTGCAACA GTCATGGCTG
ahya_s0003.g68.t2  GGACGCATTT GCAGCGCAC- --AAAGACTG GCCTGCAACA GTCATGGCTG
acyt_s1402.g1.t1   CAGTCCTGAA GCAAAAGAGG CCAGAGAAAA CCCACAGAAC CAGAGGGCCT
amur_s0006.g105.t2 GGACGCAGTT GCAGCGCAC- --AAAGACTG GCCTGCAACA GTCATGGCTG
ech_s0159.g30.t2   GGACGCAGTT GCAGCGCAC- --AAAGACTG GCCTGCAACA GTCATGGCTG
aacu_s0038.g69.t1  GGACGCAGTT GCAGCGCAC- -----GACTG GCCTGCAACA GTCATGGCTG
anas_s0109.g72.t2  GGACGCAGTT GCAGCGCAC- -----GACTG GCCTGCAACA GTCATGGCTG
amic_s0245.g8.t1   GGACGCAGTT GCAGCGCAC- -----GACTG GCCTGCAACA GTCATGGCTG
adig_s0048.g28.t1  GAACGCAGTT GCAGTGCAC- -----GACTG GCCTGCAACA GTCATGGCTG
Asp1_sequenced_cds ---------- ---------- ---------- ---------- ----------

aten_s0183.g20.t1  TGGACGGCAC TGGTCATTTT ACTGTCCTTG GATCGAGAAA A---GGATTG
ayon_s0004.g209.t1 ---------- ---------- ---------- ---------- ----------
aint_s0143.g6.t1   TGGACTGTAC TGGTCATTTT ACTGTCCTTG GAGCGAGAAA A---GGATTG
agem_s0013.g175.t1 TGGACTGTAC TGGTCATTTT ACTGTCCTTG GATCGAGAAA A---GGATTG
aawi_s0007.g175.t1 TGGACTGTAC TGGTCATTTT ACTGTCCTTG GAGCGAGAAA A---GGATTG
aflo_s0310.g20.t1  TGGACTGTAC TGGTCATTTT ACTGTCCTTG GAGCGAGAAA A---GGATTG
XM_029340177.2     TGGACTGTAC TGGTCATTTT ACTGTCCTTG GATCGAGAAA A---GGATTG
asel_s0045.g58.t2  TGGACTGTAC TGGTCATTTT ACTGTCCTTG GATCGAGAAA A---GGATTG
ahya_s0003.g68.t2  TGGACTGTAC TGGTCATTTT ACTGTCCTTG GATCGAGAAA A---GGATTG
acyt_s1402.g1.t1   TTATATGTAA CTTACAACAA CATTGGCTGA CCATACGAAA ACTTGGTCAC
amur_s0006.g105.t2 TGGACTGTAC TGGTCATTTT ACTGTCCTTG GATCGAGAAA A---GGATTG
ech_s0159.g30.t2   TGGACTGTAC TGGTCATTTT ACTGTCCTTG GATCGAGAAA A---GGATTG
aacu_s0038.g69.t1  TGGACTGTAC TGGTCATTTT ACTGTCCTTG GATCGAGAAA A---GGATTG
anas_s0109.g72.t2  TGGACTGTAC TGGTCATTTT ACTGTCCTTG GATCGAGAAA A---GGATTG
amic_s0245.g8.t1   TGGACTGTAC TGGTCATTTT ACTGTCCTTG GATCGAGAAA A---GGATTG
adig_s0048.g28.t1  TGGACTGTAC TGGTCATTTT ACTGTCCTTG GATCGAGAAA A---GGATTG
Asp1_sequenced_cds ---------- ---------- ---------- ---------- ----------

aten_s0183.g20.t1  GCATTCATTG ACTTGAGCTG T--------- ---CCCAGTG TTATCACAAA
ayon_s0004.g209.t1 ---------- ---------- ---------- ---------- ----------
aint_s0143.g6.t1   GCATTCATTG ACTTGAACTG T--------- ---CCCAATG TTATCACAAA
agem_s0013.g175.t1 GCATTCATTG ACTTGAACTG T--------- ---CCCAATG TTATCACAAA
aawi_s0007.g175.t1 GCATTCATTG ACTTGAACTG T--------- ---CCCAATG TTATCACAAA
aflo_s0310.g20.t1  GCATTCATTG ACTTGAACTG T--------- ---CCCAATG TTATCACAAA
XM_029340177.2     GCATTCATTG ACTTGAACTT T--------- ---CCCAATG TTATCACAAA
asel_s0045.g58.t2  GCATTCATTG ACTTGAACTT T--------- ---CCCAATG TTATCACAAA
ahya_s0003.g68.t2  GCATTCATTG ACTTGAACTT T--------- ---CCCAATG TTATCACAAA
acyt_s1402.g1.t1   CAGTGGTTTA ACCTCAATTC ACTTCTGGCC AAACCAGAGC TGATCACAGA
amur_s0006.g105.t2 GCATTCATTG ACTTGAACTT T--------- ---CCCAATG TTATCACAAA
ech_s0159.g30.t2   GCATTCATTG ACTTGAACTT T--------- ---CCCAATG TTATCACAAA
aacu_s0038.g69.t1  GCATTCATTG ACTTGAACTT T--------- ---CCCAATG TTATCACAAA
anas_s0109.g72.t2  GCATTCATTG ACTTGAACTT T--------- ---CCCAATG TTATCACAAA
amic_s0245.g8.t1   GCATTCATTG ACTTGAACTT T--------- ---CCCAATG TTATCACAAA
adig_s0048.g28.t1  GCATTCATTG ACTTGAACTC T--------- ---CCCAGTG TTATCACAAA
Asp1_sequenced_cds ---------- ---------- ---------- ---------- ----------
```

### Fig S5 continued

```
aten_s0183.g20.t1  AAAG------ ---------- ---------- ---------- ----------
ayon_s0004.g209.t1 ---------- ---------- ---------- ---------- ----------
aint_s0143.g6.t1   AAAG------ ---------- ---------- ---------- ----------
agem_s0013.g175.t1 AAAG------ ---------- ---------- ---------- ----------
aawi_s0007.g175.t1 AAAG------ ---------- ---------- ---------- ----------
aflo_s0310.g20.t1  AAAG------ ---------- ---------- ---------- ----------
XM_029340177.2     AAAG------ ---------- ---------- ---------- ----------
asel_s0045.g58.t2  AAAG------ ---------- ---------- ---------- ----------
ahya_s0003.g68.t2  AAAG------ ---------- ---------- ---------- ----------
acyt_s1402.g1.t1   AACATACTTG TTATTATACC TCACACAACT CCAAACTGAT GGTTATTCAA
amur_s0006.g105.t2 AAAG------ ---------- ---------- ---------- ----------
ech_s0159.g30.t2   AAAG------ ---------- ---------- ---------- ----------
aacu_s0038.g69.t1  AAAG------ ---------- ---------- ---------- ----------
anas_s0109.g72.t2  AAAG------ ---------- ---------- ---------- ----------
amic_s0245.g8.t1   AAAG------ ---------- ---------- ---------- ----------
adig_s0048.g28.t1  AAAG------ ---------- ---------- ---------- ----------
Asp1_sequenced_cds ---------- ---------- ---------- ---------- ----------

aten_s0183.g20.t1  --GTGCCAAG GAACAGCAAA TGGGAATGTA ATGCCTTGGA ATGGAATCCT
ayon_s0004.g209.t1 ---------- ---------- ---------- ---------- ----------
aint_s0143.g6.t1   --GTGCCAAG GAACAGCAAA TGGGAATGTA ATGCCCTGGA ATGGAATCCT
agem_s0013.g175.t1 --GTGCCAAG GAACAGCAAA TGGGAATGTA ATGCCCTGGA GTGGAATCCT
aawi_s0007.g175.t1 --GTGCCAAG GAACAGCAAA TGGGAATGTA ATGCCCTGGA ATGGAATCCT
aflo_s0310.g20.t1  --GTGCCAAG GAACAGCAAA TGGGAATGTA ATGCCCTGGA ATGGAATCCT
XM_029340177.2     --GTGCCAAG GAACAGCAAA TGGGAATGTA ATGCCCTGGA ATGGAATCCT
asel_s0045.g58.t2  --GTGCCAAG GAACAGCAAA TGGGAATGTA ATGCCCTGGA ATGGAATCCT
ahya_s0003.g68.t2  --GTGCCAAG GAACAGTAAA TGGGAATGTA ATGCCCTGGA ATGGAATCCT
acyt_s1402.g1.t1   TTTTTGTTGT TATGGGAAGA TTACCAGAGA GTAAAGCCGA CCATATGTTA
amur_s0006.g105.t2 --GTGCCAAG GAACAGTAAA TGGGAATGTA ATGCCCTGGA ATGGAATCCT
ech_s0159.g30.t2   --GTGCCAAG GAACAGCAAA TGGGAATGTA ATGCCCTGGA ATGGAATCCT
aacu_s0038.g69.t1  --GTGCCAAG GAACAGCAAA TGGGAATGTA ATGCCCTGGA ATGGAATCCT
anas_s0109.g72.t2  --GTGCCAAG GAACAGCAAA TGGGAATGTA ATGCCCTGGA ATGGAATCCT
amic_s0245.g8.t1   --GTGCCAAG GAACAGCAAA TGGGAATGTA ATGCCCTGGA ATGGAATCCT
adig_s0048.g28.t1  --GTGCCAAG GAACAGCAAA TGGGAATGTA ATGCCCTGGA ATGGAATCCT
Asp1_sequenced_cds ---------- ---------- ---------- ---------- ----------

aten_s0183.g20.t1  CACCTTTCGC ATGCTCATAT TTTTGCTAAT GCTTCAAATC AAAAAACAGA
ayon_s0004.g209.t1 ---------- ---------- ---------- ---------- ----------
aint_s0143.g6.t1   CACCTTTCGG ATGCTCATAT CTTTGCTAAT GCTTCAAATC AAAAAACAGA
agem_s0013.g175.t1 CACCTTTCGG ATGCTCATAT CTTTGCTAAT GCTTCAAATC AAAAAACAGA
aawi_s0007.g175.t1 CACCTTTCGG ATGCTCATAT CTTTGCTAAT GCTTCAAATC AAAAAACAGA
aflo_s0310.g20.t1  CACCTTTCCG ATGCTCATAT CTTTGCTAAT GCTTCAAATC AAAAAACAGA
XM_029340177.2     CACCTTTCGG ATGCTCATAT CTTTGCTAAT GCTTCAAATC AAAAAACAGA
asel_s0045.g58.t2  CACCTTTCGG ATGCTCATAT CTTTGCTAAT GCTTCAAATC AAAAAACAGA
ahya_s0003.g68.t2  CACCTTTCGG ATGCTCATAT CTTTGCTAAT GCTTCAAATC AAAAAACAGA
acyt_s1402.g1.t1   AAAATGTGTC CCGCTCAACT TGTGAAAAAG CCTGCCAATA AAAACACCTT
amur_s0006.g105.t2 CACCTTTCGG ATGCTCATAT CTTTGCTAAT GCTTCAAATC AAAAAACAGA
ech_s0159.g30.t2   CACCTTTCGG ATGCTCATAT CTTTGCTAAT GCTTCAAATC AAAAAACAGA
aacu_s0038.g69.t1  CACCTTTCGG ATGCTCATAT CTTTGCTAAT GCTTCAAATC AAAAAACAGA
anas_s0109.g72.t2  CACCTTTCGG ATGCTCATAT CTTTGCTAAT GCTTCAAATC AAAAAACAGA
amic_s0245.g8.t1   CACCTTTCGG ATGCTCATAT CTTTGCTAAT GCTTCAAATC AAAAAACAGA
adig_s0048.g28.t1  CACCTTTCGG ATGCTCATAT CTTTGCTAAT GCTTCAAATC AAAAAACAGA
Asp1_sequenced_cds ---------- ---------- ---------- ---------- ----------

aten_s0183.g20.t1  AATCTGGTCT TGGAGTAATG GCAAT----- ----GGATTA CAGCTGCAAA
ayon_s0004.g209.t1 ---------- ---------- ---------- ---------- ----------
aint_s0143.g6.t1   AATCTGGTCT TGGAGTAATG GCAAT----- ----GGATTA CAGCTGCAAA
agem_s0013.g175.t1 AATCTGGTCT TGGAGTAATG GCAAT----- ----GGATTA CAGCTGCAAA
aawi_s0007.g175.t1 AATCTGGTCT TGGAGTAATG GCAAT----- ----GGATTA CAGCTGCAAA
aflo_s0310.g20.t1  AATCTGGTCT TGGAGTAATG GCAAT----- ----GGATTA CAGCTGCAAA
XM_029340177.2     AATCTGGTCT TGGAGTAATG GCAGT----- ----GGATTA CAGCTGCAAA
asel_s0045.g58.t2  AATCTGGTCT TGGAGTAATG GCAGT----- ----GGATTA CAGCTGCAAA
ahya_s0003.g68.t2  AATCTGGTCT TGGAGTAATG GCAAT----- ----GGATTA CAGCTGCAAA
acyt_s1402.g1.t1   CCAAAAGTCT ATCAGTAAAT CTGATTTAAC AGCTGCACTC CAACAGGCTA
amur_s0006.g105.t2 AATCTGGTCT TGGAGTAATG GCAAT----- ----GGATTA CAGCTGCAAA
ech_s0159.g30.t2   AATCTGGTCT TGGAGTAATG GCAAT----- ----GGATTA CAGCTGCAAA
aacu_s0038.g69.t1  AATCTGGTCT TGGAGTAATG GCAGT----- ----GGATTA CAGCTGCAAA
anas_s0109.g72.t2  AATCTGGTCT TGGAGTAATA GCAGT----- ----GGATTA CAGCTGCAAA
amic_s0245.g8.t1   AATCTGGTCT TGGAGTAATG GCAAT----- ----GGATTA CAGCTGCAAA
adig_s0048.g28.t1  AATCTGGTCT TGGAGTAATG GCAAT----- ----GGATTA CAGCTGCAAA
Asp1_sequenced_cds ---------- ---------- ---------- ---------- ----------
```

### Fig S4 continued

```
aten_s0183.g20.t1  TTCTTAGAGG TCATACAAGA GCAATTAGTG ATTTAAACTG GTCTCGGTTT
ayon_s0004.g209.t1 ---------- ---------- ---------- ---------- ----------
aint_s0143.g6.t1   TTCTTAGAGG TCATACAAGA GCAATTAGTG ATCTAAACTG GTCTTGGTTT
agem_s0013.g175.t1 TTCTTAGAGG TCATACAAGA GCAATTAGTG ATCTAAACTG GTCTCGGTTT
aawi_s0007.g175.t1 TTCTTAGAGG TCATACAAGA GCAATTAGTG ATCTAAACTG GTCTTGGTTT
aflo_s0310.g20.t1  TTCTTAGAGG TCATACAAGA GCAATTAGTG ATCTAAACTG GTCTTGGTTT
XM_029340177.2     TTCTTAGAGG TCATACAAGA GCAATTAGTG ATCTAAACTG GTCTTGGTTT
asel_s0045.g58.t2  TTCTTAGAGG TCATACAAGA GCAATTAGTG ATCTAAACTG GTCTTGGTTT
ahya_s0003.g68.t2  TTCTTAGAGG TCATACAAGA GCAATTAGTG ATCTAAACTG GTCTTGGTTT
acyt_s1402.g1.t1   CAGTAGGTGG CAGAGATAAA CCTGCTGATA TGGAAGAAAT TAGAAGAAGA
amur_s0006.g105.t2 TTCTTAGAGG TCATACAAGA GCAATTAGTG ATCTAAACTG GTCTTGGTTT
ech_s0159.g30.t2   TTCTTAGAGG TCATACAAGA GCAATTAGTG ATCTAAACTG GTCTTGGTTT
aacu_s0038.g69.t1  TTCTTAGAGG TCATACAAGA GCAATTAGTG ATCTAAACTG GTCTTGGTTT
anas_s0109.g72.t2  TTCTTAGAGG TCATACAAGA GCAATTAGTG ATCTAAACTG GTCTTGGTTT
amic_s0245.g8.t1   TTCTTAGAGG TCATACAAGA GCAATTAGTG ATCTAAACTG GTCTTGGTTT
adig_s0048.g28.t1  TTCTTAGAGG TCATACAAGA GCAATTAGTG ATCTAAACTG GTCTTGGTTT
Asp1_sequenced_cds ---------- ---------- ---------- ---------- ----------

aten_s0183.g20.t1  GACCCTCAGC TACTGTCAAC CTGCTCAATG GATCAGTTTA TATACATTTG
ayon_s0004.g209.t1 ---------- ---------- ---------- ---------- ----------
aint_s0143.g6.t1   GACCCTCAGC TACTGTCCAG CTGCTCAATG GATCAGTTTA TATACATTTG
agem_s0013.g175.t1 GACCCTCAGC TACTGTCCAG CTGCTCAATG GATCAGTTTA TATACATTTG
aawi_s0007.g175.t1 GACCCTCAGC TACTGTCCAG CTGCTCAATG GATCAATTTA TATACATTTG
aflo_s0310.g20.t1  GACCCTCAGC TACTGTCCAG CTGCTCAATG GATCAGTTTA TATACATTTG
XM_029340177.2     GACCCTCAGC TACTGTCCAG CTGCTCAATG GATCAGTTTA TATACATTTG
asel_s0045.g58.t2  GACCCTCAGC TACTGTCCAG CTGCTCAATG GATCAGTTTA TATACATTTG
ahya_s0003.g68.t2  GACCCTCAGC TACTGTCCAG CTGCTCAATG GATCAGTTTA TATACATTTG
acyt_s1402.g1.t1   AGAGAATTAT ATTTTACTAG GCAACAACAA AACCAA---- -----GAAAA
amur_s0006.g105.t2 GACCCTCAGC TACTGTCCAG CTGCTCAATG GATCAGTTTA TATACATTTG
ech_s0159.g30.t2   GACCCTCAGC TGCTGTCCAG CTGCTCAATG GATCAGTTTA TATACATTTG
aacu_s0038.g69.t1  GACCCTCAGC TACTGTCCAG CTGCTCAATG GATCAGTTTA TATACATTTG
anas_s0109.g72.t2  GACCCTCAGC TACTGTCCAG CTGCTCAATG GATCAGTTTA TATACATTTG
amic_s0245.g8.t1   GACCCTCAGC TACTGTCCAG CTGCTCAATG GATCAGTTTA TATACATTTG
adig_s0048.g28.t1  GACCCTCAGC TACTGTCCAG CTGCTCAATG GATCAGTTTA TATACATTTG
Asp1_sequenced_cds ---------- ---------- ---------- ---------- ----------

aten_s0183.g20.t1  GGACTTGAGA GAGGGTAAAA AGCCAGCT-- -AGCTCATTA CAAGCTATTG
ayon_s0004.g209.t1 ---------- ---------- ---------- ---------- ----------
aint_s0143.g6.t1   GGACTTGAGG GAGGGTAAAA AGCCAGCT-- -AGCTCATTG CAAGCTATTG
agem_s0013.g175.t1 GGACTTGAGG GAGGGTAAAA AGCCAGCT-- -AGCTCATTG CAAGCTATTG
aawi_s0007.g175.t1 GGACTTGAGG GAGGGTAAAA AGCCAGCT-- -AGCTCATTG CAAGCTATTG
aflo_s0310.g20.t1  GGACTTGAGG GAGGGTAAAA AGCCAGCT-- -AGCTCATTG CAAGCTATTG
XM_029340177.2     GGACTTGAGG GAGGGTAAAA AGCCAGCT-- -AGCTCATTG CAAGCTATTG
asel_s0045.g58.t2  GGACTTGAGG GAGGGTAAAA AGCCAGCT-- -AGCTCATTG CAAGCTATTG
ahya_s0003.g68.t2  GGACTTGAGG GAGGGTAAAA AGCCAGCG-- -AGCTCATTG CAAGCTATTG
acyt_s1402.g1.t1   TGATACGACA AGAGGACAAA GTGCAGTAAG AACAGATTCA TCTCAAATAA
amur_s0006.g105.t2 GGACTTGAGG GAGGGTAAAA AGCCAGCG-- -AGCTCATTG CAAGCTATTG
ech_s0159.g30.t2   GGACTTGAGG GAGGGTAAAA AGCCAGCG-- -AGCTCATTG CAAGCTATTG
aacu_s0038.g69.t1  GGACTTGAGG GAGGGTAAAA AGCCAGCT-- -AGCTCATTG CAAGCTATTG
anas_s0109.g72.t2  GGACTTGAGG GAGGGTAAAA AGCCAGCT-- -AGCTCATTG CAAGCTATTG
amic_s0245.g8.t1   GGACTTGAGG GAGGGTAAAA AGCCAGCT-- -AGCTCATTG CAAGCTATTG
adig_s0048.g28.t1  GGACTTGAGG GAGGGTAAAA AGCCAGCT-- -AGCTCATTG CAAGCTATTG
Asp1_sequenced_cds ---------- ---------- ---------- ---------- ----------

aten_s0183.g20.t1  TTGGTGCCTC ACAAGTGAAA TGGAACAGAG TAAACCGGCA TGTCCTTGCA
ayon_s0004.g209.t1 ---------- ---------- ---------- ---------- ----------
aint_s0143.g6.t1   TTGGTGCCTC ACAAGTGAAA TGGAACAGAG TAAACCGGCA TGTCCTTGCA
agem_s0013.g175.t1 TTGGTGCCTC ACAAGTGAAA TGGAACAGAG TAAACCGGCA TGTCCTTGCA
aawi_s0007.g175.t1 TTGGTGCCTC ACAAGTGAAA TGGAACAGAG TAAACCGGCA TGTCCTTGCA
aflo_s0310.g20.t1  TTGGTGCCTC ACAAGTGAAA TGGAACAGAG TAAACCGGCA TGTCCTTGCA
XM_029340177.2     TTGGTGCTTC ACAAGTGAAA TGGAACAGAG TAAACCGGCA TGTCCTTGCA
asel_s0045.g58.t2  TTGGTGCTTC ACAAGTGAAA TGGAACAGAG TAAACCGGCA TGTCCTTGCA
ahya_s0003.g68.t2  TTGGTGCCTC ACAAGTGAAA TGGAACAGAG TAAACCGGCA TGTCCTTGCA
acyt_s1402.g1.t1   CGGGAGGACC AGACATA--- ---------G TAAACCGGCA TGTCCTTGCA
amur_s0006.g105.t2 TTGGTGCCTC ACAAGTGAAA TGGAACAGAG TAAACCGGCA TGTCCTTGCA
ech_s0159.g30.t2   TTGGTGCCTC ACAAGTGAAA TGGAACAGAG TAAACCGGCA TGTCCTTGCA
aacu_s0038.g69.t1  TTGGTGCCTC ACAAGTGAAA TGGAACAGAG TAAACCGGCA TGTCCTTGCA
anas_s0109.g72.t2  TTGGTGCCTC ACAAGTGAAA TGGAACAGAG TAAACCGGCA TGTCCTTGCA
amic_s0245.g8.t1   TTGGTGCCTC ACAAGTGAAA TGGAACAGAG TAAACCGGCA TGTCCTTGCA
adig_s0048.g28.t1  TTGGTGCCTC ACAAGTGAAA TGGAACAGAG TAAACCGGCA TGTCCTTGCA
Asp1_sequenced_cds ---------- ---------- ---------- ---------- ----------
```

### Fig S4 continued

```
aten_s0183.g20.t1  ACAAGCCATG ATGGAGATGT TCGGATCTGG GATCTTAGGA AAGGTAACAC
ayon_s0004.g209.t1 ---------- ---------- ---------- ---------- ----------
aint_s0143.g6.t1   ACAAGCCATG ATGGAGATGT TCGGATCTGG GATCTTAGGA AAGGTAACAC
agem_s0013.g175.t1 TCAAGCCATG ATGGAGATGT TCGGATCTGG GATCTTAGGA AAGGTAACAC
aawi_s0007.g175.t1 ACAAGCCATG ATGGAGATGT TCGGATCTGG GATCTTAGGA AAGGTAACAC
aflo_s0310.g20.t1  ACAAGCCATG ATGGAGATGT TCGGATCTGG GATCTTAGGA AAGGTAACAC
XM_029340177.2     ACAAGCCATG ATGGAGATGT TCGAATCTGG GATCTTAGGA AAGGTAACAC
asel_s0045.g58.t2  ACAAGCCATG ATGGAGATGT TCGAATCTGG GATCTTAGGA AAGGTAACAC
ahya_s0003.g68.t2  ACAAGCCATG ATGGAGATGT TCGGATCTGG GATCTTAGGA AAGGTAACAC
acyt_s1402.g1.t1   ACAAGCCATG ATGGAGATGT TCGGATCTGG GATCTTAGGA AAGGTAACAC
amur_s0006.g105.t2 ACAAGCCATG ATGGAGATGT TCGGATCTGG GATCTTAGGA AAGGTAACAC
ech_s0159.g30.t2   ACAAGCCATG ATGGAGATGT TCGGATCTGG GACCTTAGGA AAGGTAACAC
aacu_s0038.g69.t1  ACAAGCCATG ATGGAGATGT TCGGATCTGG GATCTTAGGA AAGGTAACAC
anas_s0109.g72.t2  ACAAGCCATG ATGGAGATGT TCGGATCTGG GATCTTAGGA AAGGTAACAC
amic_s0245.g8.t1   ACAAGCCATG ATGGAGATGT TCGGATCTGG GATCTTAGGA AAGGTAACAC
adig_s0048.g28.t1  ACAAGCCATG ATGGAGATGT TCGGATCTGG GATCTTAGGA AAGGTAACAC
Asp1_sequenced_cds ---------- ---------- ---------- ---------- ----------

aten_s0183.g20.t1  ACCTGTGGTC TACCTAACAG CCCACTTGTC TAAGATTCAT GGTCTTGATT
ayon_s0004.g209.t1 ---------- ---------- ---------- ---------- ----------
aint_s0143.g6.t1   ACCCGTGGTC TACCTAACAG CCCACTTGTC TAAGATTCAT GGTCTTGATT
agem_s0013.g175.t1 ACCCGTGGTC TACCTAACAG CCCACTTGTC TAAGATTCAT GGTCTTGATT
aawi_s0007.g175.t1 ACCCGTGGTC TACCTAACAG CCCACTTGTC TAAGATTCAT GGTCTTGATT
aflo_s0310.g20.t1  ACCCGTGGTC TACCTAACAG CCCACTTGTC TAAGATTCAT GGTCTTGATT
XM_029340177.2     ACCCGTGATC TACCTAACAG CCCACTTGTC TAAGATTCAT GGTCTTGATT
asel_s0045.g58.t2  ACCCGTGATC TACCTAACAG CCCACTTGTC TAAGATTCAT GGTCTTGATT
ahya_s0003.g68.t2  ACCCGTGATC TACCTAACAG CCCACTTGTC TAAGATTCAT GGTCTTGATT
acyt_s1402.g1.t1   ACCCGTGATC TACCTAACAG CCCACTTGTC TAAGATTCAT GGTCTTGATT
amur_s0006.g105.t2 ACCCGTGATC TACCTAACAG CCCACTTGTC TAAGATTCAT GGTCTTGATT
ech_s0159.g30.t2   ACCCGTGATC TACCTAACAG CCCACTTGTC TAAGATTCAT GGTCTTGATT
aacu_s0038.g69.t1  ACCCGTGATC TACCTAACAG CCCACTTGTC TAAGATTCAT GGTCTTGATT
anas_s0109.g72.t2  ACCCGTGATC TACCTAACAG CCCACTTGTC TAAGATTCAT GGTCTTGATT
amic_s0245.g8.t1   ACCCGTGATC TACCTAACAG CCCACTTGTC TAAGATTCAT GGTCTTGATT
adig_s0048.g28.t1  ACCCGTGATC TACCTAACAG CCCACTTGTC TAAGATTCAT GGTCTTGATT
Asp1_sequenced_cds ---------- ---------- ---------- ---------- ----------

aten_s0183.g20.t1  GGTCACGATC CAGTGGCACA ACTCTGGCAA CTTGCAGTAG TGATACTACC
ayon_s0004.g209.t1 ---------- ---------- ---------- ---------- ----------
aint_s0143.g6.t1   GGTCACGATC CAGTGGCACA ACTCTGGCAA CTTGCAGTAG TGATACTACC
agem_s0013.g175.t1 GGTCACGATC CAGTGGCACA ACTCTGGCAA CTTGCAGTAG TGATACTACG
aawi_s0007.g175.t1 GGTCACGATC CAGTGGCACA ACTCTGGCAA CTTGCAGTAG TGATACTACC
aflo_s0310.g20.t1  GGTCACGATC TAGTGGCACA ACTCTGGCAA CTTGCAGTAG TGATACTACC
XM_029340177.2     GGTCACGCTC CAGTGGCACA ACTCTGGCAA CTTGCAGTAG TGATACTACC
asel_s0045.g58.t2  GGTCACGCTC CAGTGGCACA ACTCTGGCAA CTTGCAGTAG TGATACTACC
ahya_s0003.g68.t2  GGTCATGCTC CAGTGGCACA ACTCTGGCAA CTTGCAGTAG TGATACTACC
acyt_s1402.g1.t1   GGTCACGCTC CAGTGGCACA ACTCTGGCAA CTTGCAGTAG TGATACTACC
amur_s0006.g105.t2 GGTCACGCTC CAGTGGCACA ACTCTGGCAA CTTGCAGTAG TGATACTACC
ech_s0159.g30.t2   GGTCACGCTC CAGTGGCACA ACTCTGGCAA CTTGCAGTAG TGATACTACC
aacu_s0038.g69.t1  GGTCACGCTC CAGTGGCACA ACTCTGGCAA CTTGCAGTAG TGATACTACC
anas_s0109.g72.t2  GGTCACACTC CAGTGGCACA ACTCTGGCAA CTTGCAGTAG TGATACTACC
amic_s0245.g8.t1   GGTCACGCTC CAGTGGCACA ATTCTGGCAA CTTGCAGTAG TGATACTACC
adig_s0048.g28.t1  GGTCACGCTC CAGTGGCACA ACTCTGGCAA CTTGCAGTAG TGATACTACC
Asp1_sequenced_cds ---------- ---------- ---------- ---------- ----------

aten_s0183.g20.t1  GTCAAGTTGT GGAATACTGA ACAACCTCAA CAACCAGAGA ATAAGTTGAA
ayon_s0004.g209.t1 ---------- ---------- ---------- ---------- ----------
aint_s0143.g6.t1   GTCAAGTTGT GGAATACTGA ACAACCTCAA CAACCAGAGA ATAAGTTGAA
agem_s0013.g175.t1 GTCAAGTTGT GGAATACTGA ACAACCTCAA CAACCAGAGA ATAAGTTGAA
aawi_s0007.g175.t1 GTCAAGTTGT GGAATACTGA ACAACCTCAA CAACCAGAGA ATAAGTTGAA
aflo_s0310.g20.t1  GTCAAGTTGT GGAATACTGA ACAACCTCAA CAACCAGAGA ATAAGTTGAA
XM_029340177.2     GTCAAGTTGT GGAATACTGA ACAACCTCAG CAACCAGAGA ATAAGTTGAA
asel_s0045.g58.t2  GTCAAGTTGT GGAATACTGA ACAACCTCAG CAACCAGAGA ATAAGTTGAA
ahya_s0003.g68.t2  GTCAAGTTGT GGAATACTGA ACAACCTCAG CAGCCAGAGA ATAAGTTGAA
acyt_s1402.g1.t1   GTCAAGTTGT GGAATACTGA ACAACCTCAG CAGCCAGAGA ATAAGTTGAA
amur_s0006.g105.t2 GTCAAGTTGT GGAATACTGA ACAACCTCAG CAACCAGAGA ATAAGTTGAA
ech_s0159.g30.t2   GTCAAGTTGT GGAATACTGA ACAACCTCAG CAACCAGAGA ATAAGTTGAA
aacu_s0038.g69.t1  GTCAAGTTGT GGAATACTGA ACAACCTCAG CAACCAGAGA ATAAGTTGAA
anas_s0109.g72.t2  GTCAAGTTGT GGAATATTGA ACAACCTCAG CAACCAGAGA ATAAGTTGAA
amic_s0245.g8.t1   GTCAAGTTGT GGAATACTGA ACAACCTCAG CAACCAGAGA ATAAGTTGAA
adig_s0048.g28.t1  GTCAAGTTGT GGAATACTGA ACAACCTCAG CAACCAGAGA AGAAGTTGAA
Asp1_sequenced_cds ---------- ---------- ---------- ---------- ----------
```

### Fig S4 continued

```
aten_s0183.g20.t1  TGCAAAGTGC CCTGTCTGGA GAGCAAGATT TACACCATTT GGAGAGGGAC
ayon_s0004.g209.t1 ---------- ---------- ---------- ---------- ----------
aint_s0143.g6.t1   TGCAAAGTGC CCTGTCTGGA GAGCAAGATT TACACCATTT GGAGAGGGAC
agem_s0013.g175.t1 TGCAAAGTGC CCTGTCTGGA GAGCAAGATT TACACCATTT GGAGAGGGAC
aawi_s0007.g175.t1 TGCAAAGTGC CCTGTCTGGA GAGCAAGATT TACACCATTT GGAGAGGGAC
aflo_s0310.g20.t1  TGCAAAGTGC CCTGTCTGGA GAGCAAGATT TACACCATTT GGAGAAGGAC
XM_029340177.2     TGCAAAGTGC CCTGTCTGGA GAGCAAGATT TACACCATTT GGAGAGGGAC
asel_s0045.g58.t2  TGCAAAGTGC CCTGTCTGGA GAGCAAGATT TACACCATTT GGCGAGGGAC
ahya_s0003.g68.t2  TGCAAAGTGC CCTGTCTGGA GAGCAAGATT TACACCATTT GGAGAGGGAC
acyt_s1402.g1.t1   TGCAAAGTGC CCTGTCTGGA GAGCAAGATT TACACCATTT GGAGAGGGAC
amur_s0006.g105.t2 TGCAAAGTGC CCTGTCTGGA GAGCAAGATT TACACCATTT GGAGAGGGAC
ech_s0159.g30.t2   TGCAAAGTGC CCTGTCTGGA GAGCAAGATT TACACCATTT GGAGAGGGAC
aacu_s0038.g69.t1  TGCAAAGTGC CCTGTCTGGA GAGCAAGATT TACACCATTT GGAGAGGGAC
anas_s0109.g72.t2  TGCAAAGTGC CCTGTCTGGA GAGCAAGATT TACACCATTT GGAGAGGGAC
amic_s0245.g8.t1   TGCAAAGTGC CCTGTCTGGA GAGCAAGATT TACACCATTT GGAGAGGGAC
adig_s0048.g28.t1  TGCAAAGTGC CCTGTCTGGA GAGCAAGATT TACACCATTT GGAGAGGGAC
Asp1_sequenced_cds ---------- ---------- ---------- ---------- ----------

aten_s0183.g20.t1  TTGTTACTGT AACTCTACCA CAGTTACAGC GGGGAGAGAA CAGCTTGTCT
ayon_s0004.g209.t1 ---------- ---------- ---------- ---------- ----------
aint_s0143.g6.t1   TTGTTACTGT AACTCTACCA CAATTACAGC GGGGAGAGAA CAGCTTGTCT
agem_s0013.g175.t1 TTGTTACTGT AACTCTACCA CAATTACAGC GGGGAGAGAA CAGCTTGTCT
aawi_s0007.g175.t1 TTGTTACTGT AACTCTACCA CAATTACAGC GGGGAGAGAA CAGCTTGTCT
aflo_s0310.g20.t1  TTGTTACTGT AACTCTACCA CAATTACAGC GGGGAGAGAA CAGTTTGTCT
XM_029340177.2     TTGTTACTGT AACTCTACCA CAATTACAGC GGGGAGAGAA CAACTTGTCT
asel_s0045.g58.t2  TTGTTACTGT AACTCTACCA CAATTACAGC GGGGAGAGAA CAACTTGTCT
ahya_s0003.g68.t2  TTGTTACTGT AACTCTACCA CAATTACAGC GGGGAGAGAA CAGCTTGTCT
acyt_s1402.g1.t1   TTGTTACTGT AACTCTACCA CAATTACAGC GGGGAGAGAA CAGCTTGTCT
amur_s0006.g105.t2 TTGTTACTGT AACTCTACCA CAATTACAGC GGGGAGAGAA CAGCTTGTCT
ech_s0159.g30.t2   TTGTTACTGT AACTCTACCA CAATTACAGC GGGGAGAGAA CAGCTTGTCT
aacu_s0038.g69.t1  TTGTTACTGT AACTCTACCG CAATTACAGC GGGGAGAGAA CAGCTTGTCT
anas_s0109.g72.t2  TTGTTACTGT AACTCTACCG CAATTACAGC GGGGAGAGAA CAGCTTGTCT
amic_s0245.g8.t1   TTGTTACTGT AACTCTACCA CAATTACAGC GGGGAGAGAA CAACTTGTCT
adig_s0048.g28.t1  TTGTTACTGT AACTCTACCA CAATTACAGC AGGGAGAGAA CTACTTGTCT
Asp1_sequenced_cds ---------- ---------- ---------- ---------- ----------

aten_s0183.g20.t1  CTGTGGAACA TTCCAGATGT CAACTCTCCA GTGGCCCCAC CTGTCAACAC
ayon_s0004.g209.t1 ---------- ---------- ---------- ---------- ----------
aint_s0143.g6.t1   CTGTGGAACA TTCCAGATGT CAACTCTCCA GTGGCCGCAC CTGTCAACAC
agem_s0013.g175.t1 CTGTGGAACA TTCCAGATGT CAACTCTCCA GTGGCCGCAC CTGTCAACAC
aawi_s0007.g175.t1 CTGTGGAACA TTCCAGATGT CAACTCTCCA GTGGCCGCAC CTGTCAACAC
aflo_s0310.g20.t1  CTGTGGAACA TTCCAGATGT CAACTCTCCA GTGGCCGCAC CTGTCAACAC
XM_029340177.2     CTGTGGAACA TTCCAGATGT CAACTCTCCA GTGGCCGCAC CTGTCAACAC
asel_s0045.g58.t2  CTGTGGAACA TTCCAGATGT CAACTCTCCA GTGGCCGCAC CTGTCAACAC
ahya_s0003.g68.t2  CTGTGGAACA TTCCAGATGT CAACTCTCCA GTGGCCGCAC CTGTCAACAC
acyt_s1402.g1.t1   CTGTGGAACA TTCCAGATGT CAACTCTCCA GTGGCCGCAC CTGTCAACAC
amur_s0006.g105.t2 CTGTGGAACA TTCCAGATGT CAACTCTCCA GTGGCCGCAC CTGTCAACAC
ech_s0159.g30.t2   CTGTGGAACA TTCCAGATGT CAACTCTCCA GTGGCCGCAC CTGTCAACAC
aacu_s0038.g69.t1  CTGTGGAACA TTCCAGATGT CAACTCTCCA GTGGCTGCAC CTGTCAACAC
anas_s0109.g72.t2  CTGTGGAACA TTCCAGATGT CAACTCTCCA GTGGCTGCAC CTGTCAACAC
amic_s0245.g8.t1   CTGTGGAACA TTCCAGATGT CGACTCTCCA GTGGCCGCAC CTGTCAACAC
adig_s0048.g28.t1  CTGTGGAACA TTCCAGATGT CAACTCTCCA GTGGCCGCAC CTGTCAACAC
Asp1_sequenced_cds ---------- ---------- ---------- ---------- ----------

aten_s0183.g20.t1  TTTTGTCGGC CACAGTGATG TGGTGTTGGA CTTTCATTGG AGATCCCAAA
ayon_s0004.g209.t1 ---------- ---------- ---------- ---------- ----------
aint_s0143.g6.t1   TTTTGTCGGC CACAATGATG TGGTGTTGGA CTTTCATTGG AGATCCCAAA
agem_s0013.g175.t1 TTTTGTCGGC CACAATGATG TGGTGTTGGA CTTTCATTGG AGATCCCAAA
aawi_s0007.g175.t1 TTTTGTCGGC CACAATGATG TGGTGTTGGA CTTTCATTGG AGATCCCAAA
aflo_s0310.g20.t1  TTTTGTTGGC CACAGTGATG TGGTGTTGGA CTTTCATTGG AGATCCCAAA
XM_029340177.2     TTTTGTCGGC CACACTGATG TGGTGTTGGA CTTTCATTGG AGATCCCAAA
asel_s0045.g58.t2  TTTTGTCGGC CACACTGATG TGGTGTTGGA CTTTCATTGG AGATCCCAAA
ahya_s0003.g68.t2  TTTTGTCGGC CACACTGATG TGGTGTTGGA CTTTCATTGG AGATCCCAAA
acyt_s1402.g1.t1   TTTTGTCGGC CACACTGATG TGGTGTTGGA CTTTCATTGG AGATCCCAAA
amur_s0006.g105.t2 TTTTGTCGGC CACACTGATG TGGTGTTGGA CTTTCATTGG AGATCCCAAA
ech_s0159.g30.t2   TTTTGTCGGC CACACTGATG TGGTGTTGGA CTTTCATTGG AGATCCCAAA
aacu_s0038.g69.t1  TTTTGTCGGC CACAGTGATG TGGTGTTGGA CTTTCATTGG AGATCCCAAA
anas_s0109.g72.t2  TTTTGTCGGC CACACTGATG TGGTGTTGGA CTTTCATTGG AGATCCCAAA
amic_s0245.g8.t1   TTTTGTCGGC CACACTGATG TGGTGTTGGA CTTTCATTGG AGATCCCAAA
adig_s0048.g28.t1  TTTTGTCGGC CACAGTGATG TGGTGTTGGA CTTTCATTGG AGATCCCAAA
Asp1_sequenced_cds ---------- ---------- ---------- ---------- ----------
```

### Fig S4 continued

```
aten_s0183.g20.t1  CTCTTGATAG AGATGAGCAG TTTCAACTTA TCACTTGGGC AAAGGACTGT
ayon_s0004.g209.t1 ---------- ---------- ---------- ---------- ----------
aint_s0143.g6.t1   CTCTTGATAA TGATGAGCAG TTTCAGCTTA TCACTTGGGC AAAGGACTGT
agem_s0013.g175.t1 CTCTTGATAA AGATGAGCAG TTTCAGCTTA TCACTTGGGC AAAGGACTGT
aawi_s0007.g175.t1 CTCTTGATAA AGATGAGCAG TTTCAGCTTA TCACTTGGGC AAAGGACTGT
aflo_s0310.g20.t1  CTCTTGATAA AGATGAGCAG TTTCAGCTTA TCACTTGGGC AAAGGACTGT
XM_029340177.2     CTCTTGATAA AGATGAGCAG TTTCAGCTTA TCACTTGGGC AAAGGACTGT
asel_s0045.g58.t2  CTCTTGATAA AGATGAGCAG TTTCAGCTTA TCACTTGGGC AAAGGACTGT
ahya_s0003.g68.t2  CTCTTTATGA AGATGAGCAG TTTCAGCTTA TCACTTGGGG AAAGGACTGT
acyt_s1402.g1.t1   CTCTTTATGA AGATGAGCAG TTTCAGCTTA TCACTTGGGG AAAGGACTGT
amur_s0006.g105.t2 CTCTTGATAA AGATGAGCAG TTTCAGCTTA TCACTTGGGG AAAGGACTGT
ech_s0159.g30.t2   CTCTTGATAA AGATGAGCAG TTTCAGCTTA TCACTTGGGG AAAGGACTGT
aacu_s0038.g69.t1  CTCTTGATAA AGATGAGCAG TTTCAGCTCA TCACTTGGGC AAAGGACTGT
anas_s0109.g72.t2  CTCTTGATAA AGATGAGCAA TTTCAGCTTA TCACTTGGGC AAAAGACTGT
amic_s0245.g8.t1   CTCTTGATAA AGATGAGCAG TTTCAGCTTA TCACTTGGGC AAAGGACTGT
adig_s0048.g28.t1  CTCTTGATGA AGATGAGCAG TTTCAGCTTA TCACTTGGGC AAAGGACTGT
Asp1_sequenced_cds ---------- ---------- ---------- ---------- ----------

aten_s0183.g20.t1  TGTCTTCGTT TATGGGTTCT TGAGCCAAGG ATGATTATGG CATGCAGTGG
ayon_s0004.g209.t1 ---------- ---------- ---------- ATGATTATGG CATGCAGTGG
aint_s0143.g6.t1   TGTCTTCGTT TATGGGTTCT TGAGCCAAGG ATGATTATGG CATGCAGTGG
agem_s0013.g175.t1 TGTCTTCGTT TATGGGTTCT TGAGCCAAGG ATGATTATGG CATGCAATGG
aawi_s0007.g175.t1 TGTCTTCGTT TATGGGTTCT TGAGCCAAGG ATGATTATGG CATGCAGTGG
aflo_s0310.g20.t1  TGTCTTCGTT TATGGGTTCT TGAGCCAAGG ATGATTATGG CATGCAGTGG
XM_029340177.2     TGTCTTCGTT TATGGGTTCT TGAGCCAAGG ATGATTATGG CATGCAGTGG
asel_s0045.g58.t2  TGTCTTCGTT TATGGGTTCT TGAGCCAAGG ATGATTATGG CATGCAGTGG
ahya_s0003.g68.t2  TGTCTTCGTT TATGGGTTCT TGAGCCAAGG ATGATTATGG CATGCAGTGG
acyt_s1402.g1.t1   TGTCTTCGTT TATGGGTTCT TGAGCCAAGG ATGATTATGG CATGCAGTGG
amur_s0006.g105.t2 TGTCTTCGTT TATGGGTTCT TGAGCCAAGG ATGATTATGG CATGCAGTGG
ech_s0159.g30.t2   TGTCTTCGTT TATGGGTTCT TGAGCCAAGG ATGATTATGG CATGCAGTGG
aacu_s0038.g69.t1  TGTCTTCGTT TATGGGTTCT TGAGCCAAGG ATGATTATGG CATGCAGTGG
anas_s0109.g72.t2  TGTCTTCGTT TATGGGTTCT TGAGCCAAGG ATGATTATGG CATGCAGTGG
amic_s0245.g8.t1   TGTCTTCGTT TATGGGTTCT TGAGCCAAGG ATGATTATGG CATGCAGTGG
adig_s0048.g28.t1  TGTCTTCGTT TATGGGTTCT TGAGCCAAGG ATGATTATGG CATGCAGTGG
Asp1_sequenced_cds ---------- ---------- ---------- ---------- ----------

aten_s0183.g20.t1  TGATACATCC ACCAATTTCC CTTCTCGTTC AACCAGTGAA ACAGCCCACT
ayon_s0004.g209.t1 TGATACATCC ACCAATTTCC CTTCTCGTTC AACCAGTGAA ACAGCCCACT
aint_s0143.g6.t1   TGATACATCC ACCAATTTCC CTTCTCGTTC AACCAGTGAA ACAGCCCACT
agem_s0013.g175.t1 TGATACATCC ACCAATTTCC CTTCTCCTTC AACCAGTGAA ACAGCCCACT
aawi_s0007.g175.t1 TGATACATCC ACCAATTTCC CTTCTCGTTC AACCAGTGAA ACAGCCCACT
aflo_s0310.g20.t1  TGATACATCC ACCAATTTCC CTTCTCGTTC AACCAGTGAA ACAGCCCACT
XM_029340177.2     TGATACATCC ACCAATTTCC CTTCTCGTTC AACCAATGAA ACAGCCCACT
asel_s0045.g58.t2  TGATACATCC ACCAATTTCC CTTCTCGTTC AACCAATGAA ACAGCCCACT
ahya_s0003.g68.t2  TGATACATCC ACCAATTTCC CTTCTCATTC AACCAATGAA ACAGCCCACT
acyt_s1402.g1.t1   TGATACATCC ACCAATTTCC CTTCTCATTC AACCAATGAA ACAGCCCACT
amur_s0006.g105.t2 TGATACATCC ACCAATTTCC CTTCTCGTTC AACCAATGAA ACAGCCCACT
ech_s0159.g30.t2   TGATACATCC ACCAATTTCC CTTCTCGTTC AACCAATGAA ACAGCCCACT
aacu_s0038.g69.t1  TGATACATCC ACCAATTTCC CTTCTCGTAC AACCAATGAA ACAGCCCACT
anas_s0109.g72.t2  TGATACATCC ACCAATTTCC CTTCTCGTTC AACCAATGAA ACAGCCCACT
amic_s0245.g8.t1   TGATACTTCC ACCAATTTCC CTTCTCATTC AACCAATGAA ACAGCCCACT
adig_s0048.g28.t1  TGATACATCC ACCAATTTCC CTTCTCGTTC AACCAATGAA ACAGCCCACT
Asp1_sequenced_cds ---------- ---------- ---------- ---------- ----------

aten_s0183.g20.t1  CCATGTCTGA TGAGCTGATA GAGATGGCAA TCGCAGAGTC TGTCATCATT
ayon_s0004.g209.t1 CCATGTCTGA TGAGCTGATA GAGATGGCAA TCGCAGAGTC TGTCATCATT
aint_s0143.g6.t1   CCATGTCTGA TGAGATGATA GAGATGGCAA TTGCAGAGTC TGTCATAATT
agem_s0013.g175.t1 CCATGTCTGA TGAGATGATA GAGATGGCAA TTGCAGAGTC T---------
aawi_s0007.g175.t1 CCATGTCTGA TGAGATGATA GAGATGGCAA TTGCAGAGTC TGTCATAATT
aflo_s0310.g20.t1  CCATGTCTGA TGAGATGATA GAGATGGCAA TTGCAGAGTC TGTCATAATT
XM_029340177.2     CCATGTCTGA GGAGCTGATA GAGATGGCAA TTGCAGAGTC TGTCATAATT
asel_s0045.g58.t2  CCATGTCTGA GGAGCTGATA GAGATGGCAA TTGCAGAGTC TGTCATAATT
ahya_s0003.g68.t2  CCATGTCTGA GGAGCTGATA GAGATGGCAA TTGCAGAGTC TGTCATAATT
acyt_s1402.g1.t1   CCATGTCTGA GGAGCTGATA GAGATGGCAA TTGCAGAGTC TGTCATAATT
amur_s0006.g105.t2 CCATGTCTGA GGAGCTGATA GAGATGGCAA TTGCAGAGTC TGTCATAATT
ech_s0159.g30.t2   CCATGTCTGA GGAGCTGATA GAGATGGCAA TTGCAGAGTC TGTCATAATT
aacu_s0038.g69.t1  CCATGTCTGA GGAGCTGATA GAGACGGCAA TTGCAGAGTC TGTCATAATT
anas_s0109.g72.t2  CCATGTCTGA GGAG------ ------CAAA TTGCAGAGTC TGTCAAAATT
amic_s0245.g8.t1   CCATGTCTGA GGAGCTGATA GAGATGACAA TTGCAGAGTC TGTCATAATT
adig_s0048.g28.t1  CCATGTCTGA GGAGCTGATA AAGATGGCAA TTGCAGAGTC A---------
Asp1_sequenced_cds ---------- ---------- ---------- ---------- ----------
```

### Fig S4 continued

```
aten_s0183.g20.t1  GACACACCCA AGAGTTTTCC TGATCCTGCA GCACAGGCAC AGACTTTACT
ayon_s0004.g209.t1 GACACACCCA AGAGTTTTCC TGATCCTGCA GCACAGCCAC AGACTTTACT
aint_s0143.g6.t1   GACACACCCA AGAGTTTTCC TGATCTTGCC GCACAGCCAC AGACATTACT
agem_s0013.g175.t1 ---------- ---------- ---------- ---------- ----------
aawi_s0007.g175.t1 GACACACCCA AGAGTTTTCC TGATCTTGCC GCACAGCCAC AGACATTACT
aflo_s0310.g20.t1  GACACACCCA AGAGTTTTCC TGATCTTGCC GCACAGCCAC AGACATTACT
XM_029340177.2     GACACACCTA AGAGTTTTCC TGATCTTGCA GCACAGCCAC AGACATTACA
asel_s0045.g58.t2  GACACACCTA AGAGTTTTCC TGATCTTGCA GCACAGCCAC AGACATTACA
ahya_s0003.g68.t2  GACACACCTA AGAGTTTTCC TGATCTTGCA GCACAGCCAC AGACATTACA
acyt_s1402.g1.t1   GACACACCTA AGAGTTTTCC TGATCTTGCA GCACAGCCAC AGACATTACA
amur_s0006.g105.t2 GACACACCTA AGAGTTTTCC TGATCTTGCA GCACAGCCAC AGACATTACA
ech_s0159.g30.t2   GACACACCTA AGAGTTTTCC TGATCTTGCA GCACAGCCAC AGACATTACA
aacu_s0038.g69.t1  GACACACCTA AGAGTTTTCC TGATCTTGCA GCACAGCCAC AGACATTACA
anas_s0109.g72.t2  GACACACCCA AGAGTTTTCC TGATCTTGCA GCACAGCCAC AGACATTACA
amic_s0245.g8.t1   GACACACCTA AGAGTTTTCC TGATCATGCA GCACAGCCAC AGACATTACA
adig_s0048.g28.t1  ---------- --------CC TGATCTTGCA GCACAGCCAC AGACATTACA
Asp1_sequenced_cds ---------- ---------- ---------- ---------- ----------

aten_s0183.g20.t1  GCAGGAATTT GCTCTCATTA ATGTCAACAT TCCAAATGTC ACTGTTGAGC
ayon_s0004.g209.t1 GCAGGAATTT GCTCTCATTA ATGTCAACAT TCCAAATGTC ACTGTTGAGC
aint_s0143.g6.t1   GCAGGAATTC TCTCTCATTA ATGTCAACAT TCCAAATGTC ATTGTTGAGC
agem_s0013.g175.t1 ---------- ---------- ---------- ---------- ----------
aawi_s0007.g175.t1 GCAGGAATTC TCTCTCATTA ATGTCAACAT TCCAAATGTC ATTGTTGAGC
aflo_s0310.g20.t1  GCAGGAATTC TCTCTCATTA ATGTCAACAT TCCAAATGTC ATTGTTGAGC
XM_029340177.2     GCAGGAATTC GCTCTCATTA ATGTCAACAT TCCAAATGTC ACTGTTGAGC
asel_s0045.g58.t2  GCAGGAATTC GCTCTCATTA ATGTCAACAT TCCAAATGTC ACTGTTGAGC
ahya_s0003.g68.t2  GCAGGAATTT GCTCTCATTA ATGTCAACAT TCCAAATGTC ACTGTTGAGC
acyt_s1402.g1.t1   GCAGGAATTT GCTCTCATTA ATGTCAACAT TCCAAATGTC ACTGTTGAGC
amur_s0006.g105.t2 GCAGGAATTT GCTCTCATTA ATGTCAACAT TCCAAATGTC ACTGTTGAGC
ech_s0159.g30.t2   GCAGGAATTC GCTCTCATTA ATGTCAACAT TCCAAATGTC ACTGTTGAGC
aacu_s0038.g69.t1  GCAGGAATTC GCTCTCATTA ATGTCAACAT TCCAAATGTC ACTGTTGAGC
anas_s0109.g72.t2  GCAGGAATTC GCTCTCATTA ATGTCAACAT TCCAAATGTC ACTGTTGAGC
amic_s0245.g8.t1   GCAGGAATTC GCTCTCATTA ATGTCAACAT TCCAAATGTC ACTGTTGAGC
adig_s0048.g28.t1  GCAGGAATTC GCTCTCATTA ATGTCAACAT TCCAAATGTC ACTGTTGAGC
Asp1_sequenced_cds ---------- ---------- ---------- ---------- ----------

aten_s0183.g20.t1  AGTTGGATGC TGGACACAGG AGCTGTACTG TGTGTGCTAC AAATGGACCG
ayon_s0004.g209.t1 AGTTGGATGC TGGACACAGG AGCTGTACTG TGTGTGCTAC AAATGGACCG
aint_s0143.g6.t1   AGTTGGATGC TGGGCACAGG AGCTGTACTG TGTGTGCTAC GAATGGACCG
agem_s0013.g175.t1 --TTGGATGC TGGGCACAGG AGCTGTACTG TGTGTGCTAC AAATGGACCG
aawi_s0007.g175.t1 AGTTGGATGC TGGGCACAGG AGCTGTACTG TGTGTGCTAC AAATGGACCG
aflo_s0310.g20.t1  AGTTGGATGC TGGGCACAGG AGCTGTACTG TGTGTGCTAC AAATGGACCG
XM_029340177.2     AGTTGGATGC TGGGCACAGG AGCTGTACTG TGTGTGCTAC AAGTGGACCG
asel_s0045.g58.t2  AGTTGGATGC TGGGCACAGG AGCTGTACTG TGTGTGCTAC AAGTGGACCG
ahya_s0003.g68.t2  AGTTGGATGC TGGGCACAGG AGCTGTACTG TGTGTGCTAA AAATGGACCG
acyt_s1402.g1.t1   AGTTGGATGC TGGGCACAGG AGCTGTACTG TGTGTGCTAC AAATGGACCG
amur_s0006.g105.t2 AGTTGGATGC TGGGCACAGG AGCTGTACTG CGTGTGCTAC AAATGGACCG
ech_s0159.g30.t2   AGTTGGATGC TGGGCACAGG AGCTGTACTG TGTGTGCTAC AAATGGACCG
aacu_s0038.g69.t1  AGTTGGATGC TGGGCACAGG AGCTGTACTG TGTGTGCTAC AAATGGACCG
anas_s0109.g72.t2  AGTTGGATGC TGGGCAGAGG AGCTGTACTG TGTGTGCTAC AATTGGACCG
amic_s0245.g8.t1   AGTTGGATGC TGGGCACAGG AGCTGTACTG TGTGTGCTAC AATTGGACCG
adig_s0048.g28.t1  AGTTGGATGC TGGGCACAGG AGCTGTACTG TGTGTGCTAC AAGTGGACCC
Asp1_sequenced_cds ---------- ---------- ---------- ---------- ----------

aten_s0183.g20.t1  AATACTGTTT TCTTGGTGAT AAACTTTCCC TCCCTGTACC CTAACAAAGC
ayon_s0004.g209.t1 AATACTGTTT CCTTGGTGAT AAACTTTCCC TCCCTGTACC CTAACAAAGC
aint_s0143.g6.t1   GATGTTGTTT CCTTGGTGAT AAACTTTCCC TCCCTGTACC CTAACCAAGC
agem_s0013.g175.t1 GATGTTGTTT CCTTGGTGAT AAACTTTCCC TCCCTGTACC CTAACCAAGC
aawi_s0007.g175.t1 GATGTTGTTT CCTTGGTGAT AAACTTTCCC TCCCTGTACC CTAACCAAGC
aflo_s0310.g20.t1  GATATTGTTT CCTTGGTGAT AAACTTTCCC TCCCTGTACC CTAACCAAGC
XM_029340177.2     GATACTGTTT CCTTGGTGAT AAACTTTCCC TCCCTGTACC CTAACCAAGC
asel_s0045.g58.t2  GATACTGTTT CCTTGGTGAT AAACTTTCCC TCCCTGTACC CTAACCAAGC
ahya_s0003.g68.t2  GATACTGTTT CCTTGGTGAT AAACTTTCCC TCCCTGTACC CTAACCAAGC
acyt_s1402.g1.t1   AATACTGTTT CCTTGGTGAT AAACTTTCCC TCCCTGTACC CTAACCAAGC
amur_s0006.g105.t2 GATACTGTTT CCTTGGTGAT AAACTTTCCC TCCCTGTACC CTAACCAAGC
ech_s0159.g30.t2   GATACTGTTT CCTTGGTGAT AAACTTTCCC TCCCTGTACC CTAACCAAGC
aacu_s0038.g69.t1  GATACTGTTT CCTTGGTGAT AAACTTTCCC TCCCTGTACC CTAACCAAGC
anas_s0109.g72.t2  GATACTGTTT CCTTGGTGAT AAACTTTCCC TCCCTGTACC CTAACCAAGC
amic_s0245.g8.t1   GATACTGTTT CCTTGGTGAT AAACTTTCCC TCCCTGTACC CTAACCAAGC
adig_s0048.g28.t1  GATACTGTTT CCTTGGTGAT AAACTTTCCC TCCCTGTACC CTAACCAAGC
Asp1_sequenced_cds ---------- ---------- ---------- ---------- ----------
```

### Fig S4 continued

```
aten_s0183.g20.t1  CATGCCATCA TTTGAGTTTA CAGCGGACAC CACGATTGAC ACCAATACTA
ayon_s0004.g209.t1 CATGCCATCA TTTGAGTTTA CAGCGGACAC CACGATTGAC ACCAATACTA
aint_s0143.g6.t1   CATCCCATCA TTTGAGTTTA CAGCGGACAC CTCGATTGAT ACCAATACTA
agem_s0013.g175.t1 CATCCCATCA TTTGAGTTTA CAGCGGACAC CTCGATTGAT ACCAATACTA
aawi_s0007.g175.t1 CATCCCATCA TTTGAGTTTA CAGCGGACAC CTCGATTGAT ACCAATACTA
aflo_s0310.g20.t1  CATCCCATCA TTTGAGTTTA CAGCGGACAC CTCGATTGAT ACCAATACTA
XM_029340177.2     CATCCCATCA TTTGAGTTTA CAGCGGACAC CTCGATTGAT ACCAATACTA
asel_s0045.g58.t2  CATCCCATCA TTTGAGTTTA CAGCGGACAC CTCGATTGAT ACCAATACTA
ahya_s0003.g68.t2  CATCCCATCA TTTGAGTTTA CAGCGGACAC CTCGGTTGAT ACCAATACTA
acyt_s1402.g1.t1   CATCCCATCA TTTGAGTTTA CAGCGGACAC CTCTGTTGAT ACCAATACTA
amur_s0006.g105.t2 CATCCCATCA TTTGAGTTTA CAGCGGACAC CTCGGTTGAT ACCAATACTA
ech_s0159.g30.t2   CATCCCATCA TTTGAGTTTA CAGCGGACAC CTCGATTGAT ACCAATACTA
aacu_s0038.g69.t1  CATCCCATCA TTTGAGTTTA CAGCGGACAC CTCGATTGAT ACCAATACTA
anas_s0109.g72.t2  CATCCCATCA TTTGAGTTTA CAGCGGACAC CTCGATTGAT ACCAATACTA
amic_s0245.g8.t1   CATCCCATCA TTTGAGTTTA CAGCGGACAC CTCGATTGAT ACCAATACTA
adig_s0048.g28.t1  CATCCCATCA TTTGAGTTTA CAGCGGACAC CTCGATTGAT ACCAATACTA
Asp1_sequenced_cds ---------- ---------- ---------- ---------- ----------

aten_s0183.g20.t1  AGACAAAGCT AATGAAGACC CTACGTGAAA CAGCTCAAAC TCATGTGAAA
ayon_s0004.g209.t1 AGACAAAGCT AATGAAGACC CTACGTGAAA CAGCTCAAAC TCATGTGAAA
aint_s0143.g6.t1   AGACAAAGCT AATGAAGACC CTACGTGAAA CAGCTCAGAC TCATGTGAAA
agem_s0013.g175.t1 AGACAAAGCT AATGAAGACC CTACGTGAAA CAGCTCAGAC TCATGTGAAA
aawi_s0007.g175.t1 AGACAAAGCT AATGAAGACC CTACGTGAAA CAGCTCAGAC TCATGTGAAA
aflo_s0310.g20.t1  AGACAAAGCT AATGAAGACC CTACGTGAAA CAGCTCAGAC TCATGTGAAA
XM_029340177.2     AGACAAAGCT AATGAAGACC CTACGTGAAA CAGCTCAGAC TCATGTGAAA
asel_s0045.g58.t2  AGACAAAGCT AATGAAGACC CTACGTGAAA CAGCTCAGAC TCATGTGAAA
ahya_s0003.g68.t2  AGACAAAGCT AATGAAGACC CTACGTGAAA CAGCTCAGAC TCATGTGAAA
acyt_s1402.g1.t1   AGACAAAGCT AATGAAG--- ---------- ---------- ----------
amur_s0006.g105.t2 AGACGAAGCT AATGAAGACC CTACGTGAAA CAGCTCAGAC TCATGTGAAA
ech_s0159.g30.t2   AGACAAAGCT AATGAAGACC CTACGTGAAA CAGCTCAGAC TCATGTGAAA
aacu_s0038.g69.t1  AGACAAAGCT AATGAAGACC CTACGTGAAA CAGCTCAGAC TCATGTGAAA
anas_s0109.g72.t2  AGACAAAGCT AATGAAGACC CTACGTGAAA CAGCTCAGAC TCATGTGAAA
amic_s0245.g8.t1   AGACAAAGCT AATGAAGACC CTACGTGAAA CAGCTCAGAC TCATGTGAAA
adig_s0048.g28.t1  AGACAAAGCT AATGAAGACC CTATGTGAAA CAGCTCAGAC TCATGTGAAA
Asp1_sequenced_cds ---------- ---------- ---------- ---------- ----------

aten_s0183.g20.t1  TTAAACCAAA CTTGTCTTGA ACCCTGCCTG AGGCAGCTGG TCAATCATCT
ayon_s0004.g209.t1 TTAAACCAAA CTTGTCTTGA ACCCTGCCTG AGGCAGCTGG TCAATCATCT
aint_s0143.g6.t1   TTAAACCAAA CTTGTCTTGA ACCCTGCCTA AGGCAGCTGG TCAACCATCT
agem_s0013.g175.t1 TTAAACCAAA CTTGTCTTGA ACCATGCCTG AGGCAGCTGG TCAACCATCT
aawi_s0007.g175.t1 TTAAACCAAA CTTGTCTTGA ACCCTGCCTG AGGCAGCTGG TCAACCATCT
aflo_s0310.g20.t1  TTAAACCAAA CTTGTCTTGA ACCATGCCTG AGGCAGCTGG TCAACCATCT
XM_029340177.2     TTAAACCAAA CTTGTCTTGA ACCCTGCCTG AGGCAGCTGG TCAGCCATCT
asel_s0045.g58.t2  TTAAACCAAA CTTGTCTTGA ACCCTGCCTG AGGCAGCTGG TCAGCCATCT
ahya_s0003.g68.t2  TTAAACCAAA CTTGTCTTGA ACCCTGCCTG AGGCAGCTGG TCAGCCATCT
acyt_s1402.g1.t1   ---------- ---------- ---------- ---------- ----------
amur_s0006.g105.t2 TTAAACCAAA CTTGTCTTGA ACCCTGCCTG AGGCAGCTGG TCAGCCATCT
ech_s0159.g30.t2   TTAAACCAAA CTTGTCTTGA ACCCTGCCTG AGGCAGCTGG TCAGCCATCT
aacu_s0038.g69.t1  TTAAACCAAA CTTGTCTTGA ACCCTGCCTG AGGCAGCTGG TCAGCCATCT
anas_s0109.g72.t2  TTAAACCAAA CTTGTCTTGA ACCCTGCCTG AGGCAGCTGG TCAGCCATCT
amic_s0245.g8.t1   TTAAACCAAA CTTGTCTTGA ACCCTGCCTG AGGCAGCTGG TCAGCCATCT
adig_s0048.g28.t1  TTAAACCAAA CTTGTCTTGA ACCCTGCCTG AGGCAGCTGG TCAGCCATCT
Asp1_sequenced_cds ---------- ---------- ---------- ---------- ----------

aten_s0183.g20.t1  TGATGAGGAA CTAACGCTTC AAGAAAGATT ACCCATGGAT CAGTATGCCG
ayon_s0004.g209.t1 TGATGAGGAA CTAACGCTTC AAGAAAGATT ACCCATGGAT CAGTTTGCCG
aint_s0143.g6.t1   T---GAGGAA CTAACGCTTC AAGAACGATT ACCCATGGAT CAGTTTGCTG
agem_s0013.g175.t1 T---GAGGAA CTAACGCTTC AAGAACGATT ACCCATGGAT CAGTTTGCTG
aawi_s0007.g175.t1 T---GAGGAA CTAACGCTTC AAGAACGATT ACCCATGGAT CAGTTTGCTG
aflo_s0310.g20.t1  T---GAGGAA CTAACGCTTC AAGAACGATT ACCCATGGAT CAGTTTGCTG
XM_029340177.2     T---GAGGAA CTAACGCTTC AAGAACGATT ACCCATGGAT CAGTTTGCTG
asel_s0045.g58.t2  T---GAGGAA CTAACGCTTC AAGAACGATT ACCCATGGAT CAGTTTGCTG
ahya_s0003.g68.t2  T---GAGGAA CTAACGCTTC AAGAACGATT ACCCATGGAT CAGTTTGCCG
acyt_s1402.g1.t1   ---------- ------CTTC AAGAACGATT ACCCATGGAT CAGTTTGCCG
amur_s0006.g105.t2 T---GAGGAA CTAACGCTTC AAGAACGATT ACCCATGGAT CAGTTTGCCG
ech_s0159.g30.t2   T---GAGGAA CTAACGCTTC AAGAACGATT ACCCATGGAT CAGTTTGCTG
aacu_s0038.g69.t1  T---GAGGAA CTAACGCTTC AAGAACGATT ACCCATGGAT CAGTTTGCCG
anas_s0109.g72.t2  T---GAGGAA CTAACGCTTC AAGAACGATT ACCCATGGAT CAGTTTGCCG
amic_s0245.g8.t1   T---GAGGAA CTAACGCTTC AAGAACGATT ACCCATGGAT CAGTTTGCCG
adig_s0048.g28.t1  T---GAGGAA CTAACGCTTC AAGAACGATT ACCCATGGAT CAGTTTGCCG
Asp1_sequenced_cds ---------- ---------- ---------- ---------- ----------
```

### Fig S4 continued

```
aten_s0183.g20.t1  CTACTGTGCC TCAGCCGCAT GGCCGACTCG ATCCAATCTC TTACGGTAGT
ayon_s0004.g209.t1 CTACTGTGCC TCAGCCGCAT GGCCGACTCG ATCCAATCTC TTACGGTAGT
aint_s0143.g6.t1   CTACTGTGCC TCAGCTGCAT GGCCGACTCG ATCCCATCTC TTACGGTAGT
agem_s0013.g175.t1 CTACTGTGCC TCAGCTGCAT GGCCGACTCG ATCCCATCTC TTACGGTAGT
aawi_s0007.g175.t1 CTACTGTGCC TCAGCTGCAT GGCCGACTCG ATCCCATCTC TTACGGTAGT
aflo_s0310.g20.t1  CTACTGTGCC TCAGCTGCAT GGCCGACTCG ATCCCATCTC TTACGGTAGT
XM_029340177.2     CTACTGTGCC TCAGCTGCAC GGCCGACTCG ATTCCATCTC TTACGGTAGT
asel_s0045.g58.t2  CTACTGTGCC TCAGCTGCAC GGCCGACTCG ATTCCATCTC TTACGGTAGT
ahya_s0003.g68.t2  CTACTGCCCC TCAACTGCAT GGCCGACTCG ATCCAGTCTC TTACGGTAGT
acyt_s1402.g1.t1   CTACTGCGCC TCAACTGCAT GGCCGACTCG ATCCAGTCTC TTACGGTAGT
amur_s0006.g105.t2 CTACTGCGCC TCAACTG--- ---------- --CCACTC-- -------AAC
ech_s0159.g30.t2   CTACTGTGCC TCAGCTGCAC GGCCGACTCG ATCCCATCTC TTACGGTAGT
aacu_s0038.g69.t1  CTACTGCGCC TCAACTGCAT GACCGACTCG ATCCAATCTC TTACGTTAGC
anas_s0109.g72.t2  CTACTGCGCC TCAACTGCAT GGCCAACTCG ATCCAATCTC TAACGCTAGC
amic_s0245.g8.t1   CTACTGCGCC TCAACTGCAT GACCGACTCG ATCCAATCCC TTACGTTAGC
adig_s0048.g28.t1  CTACTGCGCC TCAACTGCAT GACCGACTCG ATCCAATCTC TTACGTTAGC
Asp1_sequenced_cds ---------- ---------- ---------- ---------- ----------

aten_s0183.g20.t1  TATCAAGACG CAGCTGTTCC TTTCCCACGC ACATCTGGCG CGAAATTCTG
ayon_s0004.g209.t1 TATCAAGACG CAGCTGTTCC TTTCCCACGC ACATCTGGCG CGAAATTCTG
aint_s0143.g6.t1   TATGCTGACG CAGCTGTTCC TTTCCCACGC ACATCTGGCG CGAAATTCTG
agem_s0013.g175.t1 TATGCTGACG CAGCTGTTCC TTTCCCACGC ACATCTGGCG CGAAATTCTG
aawi_s0007.g175.t1 TATGCTGACG CAGCTGTCCC TTTCCCACGC ACATCTGGCG CGAAATTCTG
aflo_s0310.g20.t1  TACGCTGACG CAGCTGTTCC TTTCCCACGC ACATCTGGCG CGAAATTCTG
XM_029340177.2     TATGCTGACG CAGCTGTTCC ATTCCCACGC ACATCTGGCG CGAAATTCTG
asel_s0045.g58.t2  TATGCTGACG CAGCTGTTCC ATTCCCACGC ACATCTGGCG CGAAATTCTG
ahya_s0003.g68.t2  TATCTTGACG CAGCTATTCC TTTCCCACGC ACATCTGGCG CGAAATTCTG
acyt_s1402.g1.t1   TATCTTGACG CAGCTATTCC TTTCCCACGC ACATCTGGCG CGAAATTCTG
amur_s0006.g105.t2 TTTCTTGACG CAGCTATTCC TTTCCCACGC ACATCTGGCG CGAAATTCTG
ech_s0159.g30.t2   TATGCTGACG CAGCTGTTCC ATTCCCACGC ACATCTGGCG CGAAATTCTG
aacu_s0038.g69.t1  TTTCTTGACG CAGCTATTCC TTTCCCACGC ACATCTGGCG CGAAATTCTG
anas_s0109.g72.t2  TTTCTTGACG CAGCTATTCC TTTCCCACGC ACATCTGGCG CGAAATTCTG
amic_s0245.g8.t1   TTTCTTGACG CAGCTATTCC TTTCCCACGC ACATCTGGCG CGAAATTCTG
adig_s0048.g28.t1  TTTCTTGACG CAGCTATTCC TTTCCCACGC ACATCTGGCG CGAAATTCTG
Asp1_sequenced_cds ---------- ---------- ---------- ---------- ----------

aten_s0183.g20.t1  TGCTTGTGGA TTGCTGGTGT GTTTTAATTT GCCGGGAAGA TACGGTGGCC
ayon_s0004.g209.t1 TGCTTGTGGA TTGCTGGTGT GTTTTAATTT GCCGGGAAGA TACGGTGGCC
aint_s0143.g6.t1   TGCTTGTGGA TTGCTGGTGT GTTTTAATTT GCCGGGAAGA TACGGTGGCC
agem_s0013.g175.t1 TGCTTGTGGA TTGCTGGTGT GTTTTAATTT GCCGGGAAGA TACGGTGGCC
aawi_s0007.g175.t1 TGCTTGTGGA TTGCTGGTGT GTTTTAATTT GCCGGGAAGA TACGGTGGCC
aflo_s0310.g20.t1  TGCTTGTGGA TTGCTGGTGT GTTTTAATTT GCCGGGAAGA TACGGTGGCC
XM_029340177.2     TGCTTGTGGA TTGCTGGTGT GTTTTAATTT GCCGGGAAGA TACGGTGGCC
asel_s0045.g58.t2  TGCTTGTGGA TTGCTGGTGT GTTTTAATTT GCCGGGAAGA TACGGTGGCC
ahya_s0003.g68.t2  TGCTTGTGGA TTGCTGGTGT GTTTTAATTT GCCGGGAAGA TACGGTGGCC
acyt_s1402.g1.t1   TGCTTGTGGA TTGCTGGTGT GTTTTAATTT GCCGGGAAGA TACGGTGGCC
amur_s0006.g105.t2 TGCTTGTGGA TTGCTGGTGT GTTTTAATTT GCCGGGAAGA TACGGTGGCC
ech_s0159.g30.t2   TGCTTGTGGA TTGCTGGTGT GTTTTAATTT GCCGGGAAGA TACGGTGGCC
aacu_s0038.g69.t1  TGCTTGTGGA TTGCTGGTGT GTTTTAATTT GCCGGGAAGA TACGGTGGCC
anas_s0109.g72.t2  TGCTTGTGGA TTGCTGGTGT GTTTTAATTT GCCGGGAAGA TACAGTGGCC
amic_s0245.g8.t1   TGCTTGTGGA TTGCTGGTGT GTTTTAATTT GCCGGGAAGA TACGGTGGCC
adig_s0048.g28.t1  TGCTTGTGGA TTGCTGGTGT GTTTTAATTT GCCGGGAAGA TACGGTGGCC
Asp1_sequenced_cds ---------- ---------- ---------- ---------- ----------

aten_s0183.g20.t1  GGGTGGGGTC TGGTGGAGAA CCGACTCCAA GATCTCTATC AGCTTTCTCG
ayon_s0004.g209.t1 GGGTGGGGTC TGGTGGAGAA CCGACTCCAA GATCTCTATC AGCTTTCTCG
aint_s0143.g6.t1   GGGTGGGGTC TGGTGGAGAA CCGACTCCAA GATCTCTATC AGCTTTCTCG
agem_s0013.g175.t1 GGGTGGGGTC TGGTGGAGAA CCGACTCCAA GATCTCTATC AGCTTTCTCG
aawi_s0007.g175.t1 GGGTGGGGTC TGGTGGAGAA CCGACTCCAA GATCTCTATC AGCTTTCTCG
aflo_s0310.g20.t1  GGGTGGGGTC TGGTGGAGAA CCGACTCCAA GATCTCTATC AGCTTTCTCG
XM_029340177.2     GGGTGGGGTC TGGTGGAGAA CCGACTCCAA GATCTCTATC AGCTTTCTCG
asel_s0045.g58.t2  GGGTGGGGTC TGGTGGAGAA CCGACTCCAA GATCTCTATC AGCTTTCTCG
ahya_s0003.g68.t2  GGGTGGGGTC TGGTGGAGAA CCGACTCCAA GATCTCTATC AGCTTTCTCG
acyt_s1402.g1.t1   GGGTGGGGTC TGGTGGAGAA CCGACTCCAA GATCTCTATC AGCTTTCTCG
amur_s0006.g105.t2 GGGTGGGGTC TGGTGGAGAA CCGACTCCAA GATCTCTATC AGCTTTCTCG
ech_s0159.g30.t2   GGGTGGGGTC TGGTGGAGAA CCGACTCCAA GATCTCTATC AGCTTTCTCG
aacu_s0038.g69.t1  GGGTGGGGTC TGGTGGAGAA CCGACTCCAA GATCTCTATC AGCTTTCTCG
anas_s0109.g72.t2  GGGTGGGGTC TGGTGGAGAA CCGACTCCAA GATCTCTATC AGCTTTCTCG
amic_s0245.g8.t1   GGGTGGGTTC TGGTGGAGAA CCGACTCCAA GATCTCTATC AGCTTTCTCG
adig_s0048.g28.t1  GGGTGGGGTC TGGTGGAGAA CCGACTCCAA GATCTCTATC AGCTTTCTCG
Asp1_sequenced_cds ---------- ---------- ---------- ---------- ----------
```

### Fig S4 continued

```
aten_s0183.g20.t1  GCATACAGTT CTCGCCCTAG TAGTGGGCCA GCTCTTCCCC CGCTTATCAA
ayon_s0004.g209.t1 GCATACAGTT CTCGCCCTAG TAGTGGGCCA GCTCTTCCCC CGCTTATCAA
aint_s0143.g6.t1   GCATATAGTT CTCGCCCTAG TAGTGGCCCG GCTCTTCCCC CGCTTATCAA
agem_s0013.g175.t1 GCATATAGTT CTCGCCCTAG TAGTGGCCCG GCTCTTCCCC CGCTTATCAA
aawi_s0007.g175.t1 GCATATAGTT CTCGCCCTAG TAGTGGCCCT GCTCTTCCCC CGCTTATCAA
aflo_s0310.g20.t1  GCATATAGTT CTCGCCCTAG TAGTGGCCCG GCTCTTCCCC CGCTTATCAA
XM_029340177.2     GCATACAGTT CTCGCCCTAG TAGTGGCCCG GCTCTTCCCC GGCTTATCGA
asel_s0045.g58.t2  GCATACAGTT CTCGCCCTAG TAGTGGCCCG GCTCTTCCCC GGCTTGTCGA
ahya_s0003.g68.t2  GCATACAGTT CTCGCCCTAG TAGTGGCCCG GCTCTTCCCC GGCTTATCAA
acyt_s1402.g1.t1   GCATACAGTT CTCGCCCTAG TAGTGGCCCG GCTCTTCCCC GGCTTATCAA
amur_s0006.g105.t2 GCATTCAGTT TTCGCCCTAG TAGTGGCCCG GCTCTTCCCC GGCTTATCAA
ech_s0159.g30.t2   GCATACAGTT CTCGCCCTAG TAGTGGCCCG GCTCTTCCCC GGCTTATCAA
aacu_s0038.g69.t1  GCATACAGTT CTCGCCCTAG TAGTGGCCCG GCTCTTCCCC GGCTTATCAA
anas_s0109.g72.t2  GCATACAGTT CTCGCCCTAG TAGTGGCCCG GCTCTTCCCC GACTTATCAA
amic_s0245.g8.t1   GCATACAGTT CTCGCCCTAG TAGTGGCCCG GCTCTTCCCC GGCTTATCAA
adig_s0048.g28.t1  GCATACAGTT CTCGCCCTAG TAGTGGCCCG GCTCTTCCCC GACTTATCAA
Asp1_sequenced_cds ---------- ---------- ---------- ---------- ----------

aten_s0183.g20.t1  CAGACATTTT CCAAAGACTC ACCAAAGACA TGCTTTCGGT CCGTTTGGCC
ayon_s0004.g209.t1 CAGACATTTT GCAAAGACTC ACCAAAGACA TGCTTTCGGT CCGTTTGGCC
aint_s0143.g6.t1   CAGACATTTT CCAAAGACTC GCCAAAGACA TGCTTTCGGT TCGTTTGGCC
agem_s0013.g175.t1 CAGACATTTT CAAAAGAATC GCCAAAGACA TGCTTTCGGT TCGTTTGGCC
aawi_s0007.g175.t1 CAGACATTTT CAAAAGAATC GCCAAAGACA TGCTTTCGGT TCGTTTGGCC
aflo_s0310.g20.t1  CAGACATTTT CAAAAGAATC GCCAAAGACA TGCTTTCGGT TCGTTTGGCC
XM_029340177.2     CAGACATTTT CAAAATTCTC GCGAAAGACA TGCTTTGGGT TCGTTTGACC
asel_s0045.g58.t2  CAGACATTTT CAAAATTCTC GCGAAAGACA TGCTTTGGGT TCGTTTGACC
ahya_s0003.g68.t2  CAGACATTTT CAAAATCCTC GCGAAAGACA TGCTTTGGGT TCGTTTGACC
acyt_s1402.g1.t1   CAGACATTTT CAAAATCCTC GCGAAAGACA TGCTTTGGGT TCGTTTGACC
amur_s0006.g105.t2 CAGACATTTT CAAAAGCCTC GCGAAAGACA TGCTTTGGGT TCGTTTGGCC
ech_s0159.g30.t2   CAGACATTTT CAAAATCCTC GCGAAAGACA TGCTTTGGGT TCGTTTGACC
aacu_s0038.g69.t1  CAGACATTTT CAAAAGCCTC TCCAAAGACA TGCTTTCGGT TCGTTTGGCC
anas_s0109.g72.t2  CAGACATTTT CAAAAGCCTC TCCAAAGACA TGGTTTCGGT TCGTTTGGCC
amic_s0245.g8.t1   CAGACATTTT CAAAAGCCTC TCCAAAGACA TGCTTTCGGT TCGTTTGGCC
adig_s0048.g28.t1  CGGACATTTT CAAAAGCCTC TCCAAAGACA TACTTTCGGT TCGTTTCGCC
Asp1_sequenced_cds ---------- ---------- ---------- ---------- ----------

aten_s0183.g20.t1  AGCAGGCAAG GACCTTCACT TACAGAGCTT CAAGCAAGAC AGAAGACCCA
ayon_s0004.g209.t1 AGCAGGCAAG GACCTTCACT TACAGAGCTT TAAGCAAGAC AGAAGACCCA
aint_s0143.g6.t1   AGCAGACAAG GACCTTCACG TACAGAGCTT TAAGCAAGAC AGAAGACCCA
agem_s0013.g175.t1 AGCAGACAAG GTCCTTCACG TACAGAGCTT TAAGCAAGAC AGAAGACCCA
aawi_s0007.g175.t1 AGCAGACAAG GTCCTTCACG TACAGAGCTT TAAGCAAGAC AGAAGACCCA
aflo_s0310.g20.t1  AGCAGTCAAG GACCTTCACG TACAGAGCTT TAAGCAAGAC AGAAGACCCA
XM_029340177.2     AGCAGACAAG GACCTTCACG TACAGAGCTT TAAGCAGGAC AGAAGACCCA
asel_s0045.g58.t2  AGCAGACAAG GACCTTCACG TACAGAGCTT TAAGCAGGAC AGAAGACCCA
ahya_s0003.g68.t2  AGCAGACAAG GACCTTCACG TACAGAGCTT TAAGCAGGAC AGAAGACCCA
acyt_s1402.g1.t1   AGCAGACAAG GACCTTCACG TACAGAGCTT TAAGCAGGGT T---------
amur_s0006.g105.t2 AGCAGACAAG GACCTTCACG TACAGAGCTT TAAGCAGGAC AGAAGACCCA
ech_s0159.g30.t2   AGCAGACAAG GACCTTCACG TACAGAGCTT TAAGCAGGAC AGACGACCCA
aacu_s0038.g69.t1  AGCTGACAAG GACCTTCACG TACAGAGCTT TAAGCAGGAC AGAAGACCCA
anas_s0109.g72.t2  AGCTGACAAG GACCTTCACG TACAGAGCTT TAAGCAGGAC AGAAGACCCA
amic_s0245.g8.t1   AGCTGACAAG GACCTTCACG TACAGAGCTT TAAGCAGGAC AGAAGACCCA
adig_s0048.g28.t1  AGCTGACAAG GACCTTCACG TACAGAGCTT TAAGCAGGAC AGAAGACCCA
Asp1_sequenced_cds ---------- ---------- ---------- ---------- ----------

aten_s0183.g20.t1  CAGGATAGGC AAAAGTACAG TGGGGTACCC TCGAGGAACA GTGATGTTGG
ayon_s0004.g209.t1 CAGGATAGGC AAAAGTACAG TGGGGTACCC TCGAGGAACA GTGATGTTGG
aint_s0143.g6.t1   CAGGATAGGC AAAAGTACAG TGGGGTACCC TCTAGGAACA GTGATGTTGG
agem_s0013.g175.t1 CAGGATAGGC AAAAGTACAG TGGGGTACCC TCTAGGAACA GTGATGTTGG
aawi_s0007.g175.t1 CAGGATAGGC AAAAGTACAG TGGGGTACCC TCTAGGAACA GTGATGTTGG
aflo_s0310.g20.t1  CAGGATAGGC AAAAGTACAG TGGGGTACCC TCTAGGAACA GTGATGTTGG
XM_029340177.2     CAGGATAAGC AAAAGTACAG TGGGTTACCC TCTAGGACCA GTGATGTTGG
asel_s0045.g58.t2  CAGGATAGGC AAAAGTACAG TGGGTTACCC TCTAGGACCA GTGATGTTGG
ahya_s0003.g68.t2  CAGGATAGGC AAAAGTACAG TGGGTTACCC TCTAGGACCA GTGTTGTTGG
acyt_s1402.g1.t1   ---------- ---------- ---------- ---------- ----------
amur_s0006.g105.t2 CAGGATAGGC AAAAGTACAG TGGGTTACCC TCTAGGACCA GTGATGTTGG
ech_s0159.g30.t2   CAGGATAGGC AAAAGTACAG TGGGTTACCC TCTAGGACCA GTGTTGTTGG
aacu_s0038.g69.t1  CAGGATAGGC AAAAGTACAG TGGGTTACCC TCTAGGAACA GTGATGTTGG
anas_s0109.g72.t2  CAGGACAGGC AAAAGTACAG TGGGTTCCCC TCTAGGAACA GTGATGTTGG
amic_s0245.g8.t1   CAGGATAGGC AAAAGTACAG TGGGTTCCCC TCTAGGAACA GTGATGTTGG
adig_s0048.g28.t1  CAGGATAGGC AAAAGTACAG TGGGTTACCC TCTAGGAAGA GTGATGTTGG
Asp1_sequenced_cds ---------- ---------- ---------- ---------- ----------
```

### Fig S4 continued

```
aten_s0183.g20.t1  CCTGGTTGTG ATTCGTGACG TCAGTGCTAT GATGCCCATC CATCAGTCGC
ayon_s0004.g209.t1 CCTGGTTGTG ATTCGTGACG TCAGTGCTAT GATGCCCATC CATCAGTCGC
aint_s0143.g6.t1   CCTGGTTGTG ATTCGTGACG TCAGTGCTAT GATGCCCATC CATCAGTCGC
agem_s0013.g175.t1 CCTGGTTGTG ATTCGTGACG TCAGTGCTAT GATGCCCATC CATCAGTCGC
aawi_s0007.g175.t1 CCTGGTTGTG ATTCGTGACG TCAGTGCTAT GATGCCCATC CATCAGTCGC
aflo_s0310.g20.t1  CCTGGTTGTG ATTCGTGACG TCAGTGCTAT GATGCCCATC CATCAGTCGC
XM_029340177.2     CCTGGTTGTG ATTCGTGACG TCAGTGCTAT GATGCCCATC CATCAGTCGC
asel_s0045.g58.t2  CCTGGTTGTG ATTCGTGACG TCAGTGCTAT GATGCCCATC CATCAGTCGC
ahya_s0003.g68.t2  CCTGGTTGTG ATTCGTGACG TCAGTGCTAT GATGCCCATC CATCAGTCGC
acyt_s1402.g1.t1   ---------- ---------- ---------- ---------- ----------
amur_s0006.g105.t2 CCTGGTTGTG ATTCGTGACG TCAGTGCTAT GATGCCCATC CATCAGTCGC
ech_s0159.g30.t2   CCTGGTTGTG ATTCGTGACG TCAGTGCTAT GATGCCCATC CATCAGTCGC
aacu_s0038.g69.t1  CCTGGTTGTG ATTCGCGACG TCAGTGCTAT GATGCCCATC CATCAGTCGC
anas_s0109.g72.t2  CCTGGTTGTG ATTCGTGACG TCAGTGCTAT GATGCCCATC CATCAGTCGC
amic_s0245.g8.t1   CCTGGTTGTG ATTCGTGACG TCAGTGCTAT GATGCCCATC CATCAGTCGC
adig_s0048.g28.t1  CCTGGTTGTG ATTCGTGACG TCAGTGCTAT GATGCCCATC CATCAGTCGC
Asp1_sequenced_cds ---------- ---------- ---------- ---------- ----------

aten_s0183.g20.t1  TGGGTCAGGC ATACACGTTG GAAGGGAACA ATATCACTAA GATTTGCAAG
ayon_s0004.g209.t1 TGGGTCAGGC ATACACGTTG GAAGGGAACA ATATCACTAA GATTTGCAAG
aint_s0143.g6.t1   TGGGTCAGGC ATACACGTTG GAAGGGAACA ATATCACTAA GATTTGCAAG
agem_s0013.g175.t1 TGGGTCAGGC ATACACGTTG GAAGGGAACA ATATCACTAA GATTTGCAAG
aawi_s0007.g175.t1 TGGGTCAGGC ATACACGTTG GAAGGGAACA ATATCACTAA GATTTGCAAG
aflo_s0310.g20.t1  TGGGTCAGGC ATACACGTTG GAAGGGAACA ATATCACTAA GATTTGCAAG
XM_029340177.2     TGGGTCAGGC ATACACGTTG GAAGGGAACA ATATCACTAA GATTTGCAAG
asel_s0045.g58.t2  TGGGTCAGGC ATACACGTTG GAAGGGAACA ATATCACTAA GATTTGCAAG
ahya_s0003.g68.t2  TGGGTCAGGC ATACACGTTG GAAGGGAACA ATATCACTAA GATTTGCAAG
acyt_s1402.g1.t1   ---------- ---------- ---------- ---------- ----------
amur_s0006.g105.t2 TGGGTCAGGC ATACACGTTG GAAGGGAACA ATATCACTAA GATTTGCAAG
ech_s0159.g30.t2   TGGGTCAGGC ATACACGTTG GAAGGGAACA ATATCACTAA GATTTGCAAG
aacu_s0038.g69.t1  TGGGTCAGGC ATACACGTTG GAAGGGAACA ATATCACTAA GATTTGCAAG
anas_s0109.g72.t2  TGGGTCAGGC ATACACGTTG GAAGGGAACA ATATCACTAA GATTTGCAAG
amic_s0245.g8.t1   TGGGTCAGGC ATACACGTTG GAAGGGAACA ACATCACTAA GATTTGCAAG
adig_s0048.g28.t1  TGGGTCAGGC ATACACGTTG GAAGGGAACA ATATCACTAA GATTTGCAAG
Asp1_sequenced_cds ---------- ---------- ---------- ---------- ----------

aten_s0183.g20.t1  AAAAACCTCT CTGCTGCCAT GACAACGGGA CGAAAAGACC TGGTGCAGAC
ayon_s0004.g209.t1 AAAAACCTCT CTGCTGCCAT GACAACGGGA CGAAAAGACC TGGTGCAGAC
aint_s0143.g6.t1   AAAAACCTGT CTGCTGCCAT GACAACGGGA CGAAAAGACC TAGTGCAGAC
agem_s0013.g175.t1 AAAAACCTGT CTGCTGCCAT GACAACGGGA CGAAAAGACC TAGTGCAGAC
aawi_s0007.g175.t1 AAAAACCTGT CTGCTGCCAT GACAACGGGA CGAAAAGACC TAGTGCAGAC
aflo_s0310.g20.t1  AAAAACCTGT CTGCTGCCAT GACAACGGGA CGAAAAGACC TAGTGCAGAC
XM_029340177.2     AAAAACCTGT CTGCTGCCAT GACAACGGGA CGAAAAGACC TGGTGCAGAC
asel_s0045.g58.t2  AAAAACCTGT CTGCTGCCAT GACAACGGGA CGAAAAGACC TGGTGCAGAC
ahya_s0003.g68.t2  AAAAACCTGT CTGCTGCCAT GACAACGGGA CGAAAAGACC TGGTGCAGAC
acyt_s1402.g1.t1   ---------- ---------- ---------- ---------- ----------
amur_s0006.g105.t2 AAAAACCTGT CTGCTGCCAT GACAACGGGA CGAAAAGACC TGGTGCAGAC
ech_s0159.g30.t2   AAAAACCTGT CTGCTGCCAT GACAACGGGA CGAAAAGACC TGGTGCAGAC
aacu_s0038.g69.t1  AAAAACCTGT CTGCTGCCAT GACAACGGGA CGAAAAGACC TGGTGCAGAC
anas_s0109.g72.t2  AAAAACCTGT CTGCTGCCAT GAAAACGGGA CGAAAAGACC TGGTGCAGAC
amic_s0245.g8.t1   AAAAACCTGT CTGCTGCCAT GACAACGGGA CGAAAAGACC TGGTGCAGAC
adig_s0048.g28.t1  AAAAACCTGT CTGCTGCCAT GACAACGGGA CGAAAAGACC TGGTGCAGAC
Asp1_sequenced_cds ---------- ---------- ---------- ---------- ----------

aten_s0183.g20.t1  ATGGTCTCTT TTGAGCTTAG TACTCGATGA GAAGCTG--- ----------
ayon_s0004.g209.t1 ATGGTCTCTT TTGAGCTTAG TACTCGATGA GAAGCTG--- ----------
aint_s0143.g6.t1   ATGGTCTCTT TTAAGCTTAG TACTCGATGA GAAGCTG--- ----------
agem_s0013.g175.t1 ATGGTCTCTT TTAAGCTTAG TACTCGATGA GAAGCTG--- ----------
aawi_s0007.g175.t1 ATGGTCTCTT TTAAGCTTAG TACTCGATGA GAAGCTG--- ----------
aflo_s0310.g20.t1  ATGGTCTCTT TTAAGCTTAG TACTCGATGA GAAGCTG--- ----------
XM_029340177.2     ATGGTCTCTT TTGAGCTTAG TGCTCGATGA GAAGCTG--- ----------
asel_s0045.g58.t2  ATGGTCTCTT TTGAGCTTAG TGCTCGATGA GAAGCTG--- ----------
ahya_s0003.g68.t2  ATGGTCTCTT TTGAGCTTAG TGCTCGATGA GAAGCTG--- ----------
acyt_s1402.g1.t1   ---------- ---------- ---------- ---------- ----------
amur_s0006.g105.t2 ATGGTCTCTT TTGAGCTTAG TGCTCGATGA GAAGCTG--- ----------
ech_s0159.g30.t2   ATGGTCTCTT TTGAGCTTAG TGCTCGATGA GAAGCTG--- ----------
aacu_s0038.g69.t1  ATGGTCTCTT TTGAGCTTAG TGCTCGATGA GAAGCTGGCT CCCTCAGACT
anas_s0109.g72.t2  ATGGTCTCTT TTGAGCTTAG TGCTCGATGA GAAGCTGGCT CCCTCAGACC
amic_s0245.g8.t1   ATGGTCTCTT TTGAGCTTAG TGCTCGATGA GAAGCTG--- ----------
adig_s0048.g28.t1  ATGGTCTCTT TTGAGCTTAG TGCTCGATGA GAAGCTG--- ----------
Asp1_sequenced_cds ---------- ---------- ---------- ---------- ----------
```

### Fig S4 continued

```
aten_s0183.g20.t1  ---------- -GCTCCGTCA GACTCCATGG ACGAGGCGCC CTGGGCGCTG
ayon_s0004.g209.t1 ---------- -GCTCCGTCA GACTCCATGG ACGAGGCGCC CTGGGCGCTG
aint_s0143.g6.t1   ---------- -GCTCCCTCA GACTCCATGG ACGAGGCGCC CTGGGCGCTG
agem_s0013.g175.t1 ---------- -GCTCCCTCA GACTCCATGG ACGAGGCGCC CTGGGCGCTG
aawi_s0007.g175.t1 ---------- -GCTCCCTCA GACTCCATGG ACGAGGCGCC CTGGGCGCTG
aflo_s0310.g20.t1  ---------- -GCTCCCTCA GACTCCATGG ACGAGGCGCC CTGGGCGCTG
XM_029340177.2     ---------- -GCTTCCTCA GACTCCATGG ACGAGGCGCC CTGGGCGCTG
asel_s0045.g58.t2  ---------- -GCTTCCTCA GACTCCATGG ACGAGGCGCC CTGGGCGCTG
ahya_s0003.g68.t2  ---------- -GCTTCCTCA GACTCCATGG ACCAGGCGCC CTGGGCGCTG
acyt_s1402.g1.t1   ---------- ---------- ---------- ---------- ----------
amur_s0006.g105.t2 ---------- -GCTTCCTCA GACTCCATGG ACGAGGCGCC CTGGGCGCTG
ech_s0159.g30.t2   ---------- -GCTTCCTCA GACTCCATGG ACGAGGCGCC CTGGGCGCTG
aacu_s0038.g69.t1  CCATCGACGA GGCTCCCTCA GACTCCATCG ACGAGGCGCC CTGGGCGCTG
anas_s0109.g72.t2  CCATCGACGA GGCTCCCTCA GACTCCATCG ACGAGGCGCC CTGGGCGCTG
amic_s0245.g8.t1   ---------- -GCTCATTCA AACTCCATGG ACGAGGCGCC CTGGGCGCTG
adig_s0048.g28.t1  ---------- -GCTCCCTCA GACTCCATCG ACGAGGCGCC CTGGGCGCTG
Asp1_sequenced_cds ---------- ---------- ---------- ---------- ----------

aten_s0183.g20.t1  CACCCATTTG GCAAGAAATT GGTCAATTCC CTGATGGACT ATTATTTGAA
ayon_s0004.g209.t1 CACCCATTTG GCAAGAAATT GGTCAATTCC CTGATGGACT ATTATTTGAA
aint_s0143.g6.t1   CATCCATTTG GCAAGAAATT GGTCAGTTCC CTGATGGACT ATTATTTGAA
agem_s0013.g175.t1 CATCCATTTG GCAAGAAATT GGTCAGTTCC CTGATGGACT ATTATTTGAA
aawi_s0007.g175.t1 CATCCATTTG GCAAGAAATT GGTCAGTTCC CTTATGGACT ATTATTTGAA
aflo_s0310.g20.t1  CATCCATTTG GCAAGAAATT GGTCAGTTCC CTGATGGACT ATTATTTGAA
XM_029340177.2     CATCCCTTTG GCAAGAAATT GGTCAGTTCC CTGATGGACT ATTATTTGAA
asel_s0045.g58.t2  CATCCCTTTG GCAAGAAATT GGTCAGTTCC CTGATGGACT ATTATTTGAA
ahya_s0003.g68.t2  CATCCCTTTG GCAAGAAATT GGTCAGTTCC CTGATGGACT ATTATTTGAA
acyt_s1402.g1.t1   ---------- ---------- ---------- ---------- ----------
amur_s0006.g105.t2 CATCCCTTTG GCAAGAAATT GGTCAGTTCC CTGATGGACT ATTATTTGAA
ech_s0159.g30.t2   CATCCCTTTG GCAAGAAATT GGTCAGTTCC CTGATGGACT ATTATTTAAA
aacu_s0038.g69.t1  CATCCATTTG GCAAGAAATT GGTCAGTTCC CTGATGGACT ATTATTTGAA
anas_s0109.g72.t2  CATCCATTTG GCAAGAAATT GGTCAGTTCC CTGATGGACT ATTATTTGAA
amic_s0245.g8.t1   CATCCATTTG GCAAGAAATT GGTCAGTTCC CTGATGGAAT ATTATTTGAA
adig_s0048.g28.t1  CATCCATTTG GCAAGAAATT AGTCAGTTCC CTGATGGACT ATTATTTGAA
Asp1_sequenced_cds ---------- ---------- ---------- ---ATGGACT ATTATTTGAA

aten_s0183.g20.t1  CATCCGGGAT ATCCAGACAC TAGGAATGTT GTCATGTGTG CTTGCGCATC
ayon_s0004.g209.t1 CATCCGGGAT ATCCAGACAC TAGGAATGTT GTCATGTGTG CTTGCGCATC
aint_s0143.g6.t1   CATCCGGGAT ATCCAGACAC TAGGAATGTT GTCATGTGTG CTTGCGCATC
agem_s0013.g175.t1 CATCCGGGAT ATCCAGACAC TAGGAATGTT GTCATGTGTG CTTGCGCATC
aawi_s0007.g175.t1 CATCCGGGAT ATCCAGACAC TAGGAATGTT GTCATGTGTG CTTGCGCATC
aflo_s0310.g20.t1  CATCCGGGAT ATCCAGACAC TAGGAATGTT GTCATGTGTG CTTGCGCATC
XM_029340177.2     CATCCGGGAT ATCCAGACAC TAGGAATGTT GTCATGTGTG CTTGCACATC
asel_s0045.g58.t2  CATCCGGGAT ATCCAGACAC TAGGAATGTT GTCATGTGTG CTTGCGCATC
ahya_s0003.g68.t2  CATCCGGGAT ATCCAGACAC TAGGAATGTT GTCATGTGTG CTTGCGCATC
acyt_s1402.g1.t1   ---------- ---------- --GGTGAATT GTGA------ ----------
amur_s0006.g105.t2 CATCCGGGAT ATCCAGACAC TAGGAATGTT GTCATGTGTG CTTGCGCATC
ech_s0159.g30.t2   CATCCGGGAT ATCCAGACAC TAGGAATGTT GTCATGTGTG CTTGCGCATC
aacu_s0038.g69.t1  CATCCGGGAT ATCCAGACAC TAGGAATGTT GTCATGTGTA CTTGCGCATC
anas_s0109.g72.t2  CATCCGGGAT ATCCAGACAC TAGGAATGTT GTCATGTGTG CTTGCGCATC
amic_s0245.g8.t1   CATCCGGGAT ATCCAGACAC TAGGAATGTT GTCATGTGTG CTTGCGCATC
adig_s0048.g28.t1  CATCCGGGAT ATCCAGACAC TAGGAATGTT GTCATGTGTG CTTGCGCATC
Asp1_sequenced_cds CATCCGGGAT ATCCAGACAC TAGGAATGTT GTCATGTGTG CTTGCGCATC

aten_s0183.g20.t1  ACGCACTTTC AGACGCTTTC AAACCTCCCA GAAACAGCTT TAACACTGAG
ayon_s0004.g209.t1 ACGCACTTTC AGACGATTTC AAACCTCCCA GAAACAGCTT TAACACTGAG
aint_s0143.g6.t1   ACGCCCTTTC AGACGCTTTC AAACCTCCCA GAAACAGCTT TAACACTGAG
agem_s0013.g175.t1 ACGCCCTTTC AGACGCTTTC AAACCTCCCA GAAACAGCTT TAACACTGAG
aawi_s0007.g175.t1 ACGCCCTTTC AGACGCTTTC AAACCTCCCA GAAACAGCTT TAACACTGAG
aflo_s0310.g20.t1  ACGCCCTTTC AGACGCTTTC AAACCTCCCA GAAACAGCTT TAACACTGAG
XM_029340177.2     ACGCCCTTTC AGACGCTTTC AAACCTCCCA GAAACAGCTT TAAGACTGGG
asel_s0045.g58.t2  ACGCCCTTTC AGACGCTTTC AAACCTCCCA GAAACAGCTT TAAGACTGGG
ahya_s0003.g68.t2  ACGCCCTTTC AGACGCTTTC AAACCTCCCA GAAACAGCTT TTACACTGAG
acyt_s1402.g1.t1   ---------- ---------- ---------- ---------- ----------
amur_s0006.g105.t2 ACGCCCTTTC AGACGCTTTC AAACCTCCCA GAAACAGCTT TAACACTGAG
ech_s0159.g30.t2   ACGCCCTTTC AGACGCTTTC AAACCTCCCA GAAACAGCTT TAACACTGAG
aacu_s0038.g69.t1  ACGCTCTTTC AGACGCTTTC AAACCTCTCA GAAACAGCTT TAACACTGAG
anas_s0109.g72.t2  ACGCCCTTTC AGACGCTTTC AAACCTCACA GAGACAGCTT TAACACTGAG
amic_s0245.g8.t1   ACGCCCTTTC AGACGCTTTC AAACCTCTCA GAAACAGCTT TAACACTGAG
adig_s0048.g28.t1  ACGCCCTTTC AGACGCTTTC AAACCTCCCA GAAACAGCTT TAACACTGAG
Asp1_sequenced_cds ACGCCCTTTC AGACGCTTTC AAACCTTCCA GAAACAGCTT TAACACTGAG
```

### Fig S4 continued

```
aten_s0183.g20.t1  GTGATTCCAT CAAGCATGTC ATTTCCTTTT GGCTCCCCAC CGTCCTTTAA
ayon_s0004.g209.t1 GTGATTCCAT CAAGCATGTC ATTTCCTTTT GGCTCCCCAC CGTCCTTTAA
aint_s0143.g6.t1   GTGATTTCAT CAAGCATGTC ATTTCCTTTT GGCTCTCCAC CGTCCTTTAA
agem_s0013.g175.t1 GTGATTCCAT CAAGCATGTC ATTTCCTTTT GGCTCTCCAC CGTCCTTTAA
aawi_s0007.g175.t1 GTGATTCCAT CAAGCATGTC ATTTCCTTTT GGCTCTCCAC CGTCCTTTAA
aflo_s0310.g20.t1  GTGATTTCAT CAAGCATGTC ATTTCCTTTT GGCTCTCCAC CGTCCTTTAA
XM_029340177.2     GTGATTTCAT CAAGCATGTC ATTTCCTTTT GGCTCTCCAC CGTCCTTTAA
asel_s0045.g58.t2  GTGATTTCAT CAAGCATGTC ATTTCCTTTT GGCTCTCCAC CGTCCTTTAA
ahya_s0003.g68.t2  GTGATTTCAT CAAGCATGTC ATTTCCTTTT GGCTCTCCAC CGTCCTTTAA
acyt_s1402.g1.t1   ---------- ---------- ---------- ---------- ----------
amur_s0006.g105.t2 GTGATTTCAT CAAGCATGTC ATTTCCTTTT GGCTCTCCAC CGTCCTTTAA
ech_s0159.g30.t2   GTGATTTCAT CAAGCATGTC ATTTCCTTTT GGCTCTCCAC CGTCCTTTAA
aacu_s0038.g69.t1  GTGATTCCAT CAAGCATGTC ATTTCATTTT GGCTCTCCAT CGTCCTTTAA
anas_s0109.g72.t2  GAGATTTCAT CAAGCA---- ---------- ---------- CGTCCTTTAA
amic_s0245.g8.t1   GTGATTCCAT CAAGCATGTC ATTTCCTTTT GGCTCTCCAC AATCCTTTAA
adig_s0048.g28.t1  GTGATTTCAT CAAGCATGTC ATTTCCTTTT GGCTCTCCAT CGTCCTTTAA
Asp1_sequenced_cds GAGATTTCAT CAAGCA---- ---------- ---------- CGTCCTTTAA

aten_s0183.g20.t1  CAACGATTCT CCCTCACAGT CTGTCAAGCG CCCTAAACCT GTTGGGAGTG
ayon_s0004.g209.t1 CAACGATTCT CCCTCACAGT CTGTCAAGCG CCCTAAACCT GTTGGGAGTG
aint_s0143.g6.t1   CAACGATACT CCCTCACAGT CTGTCAAGCG CCCTAAACCT GTTGGGAGTG
agem_s0013.g175.t1 CAACGATACT CCCTCACAGT CTGTCAAGCG CCCTAAACCT GTTGGGAGTG
aawi_s0007.g175.t1 CAACGATACT CCCTCACAGT CTGTCAAGCG CCCTAAACCT GTTGGGAGTG
aflo_s0310.g20.t1  CAACGATACT CCCTCACAGT CTGTCAAGCG CCCTAAACCT GTTGGGAGTG
XM_029340177.2     CAACGATACT CCCTCACAGT CTGTCAAGCG CCCTAAACCT GTTGGGAGTG
asel_s0045.g58.t2  CAACGATACT CCCTCACAGT CTGTCAAGCG TCCTAAACCT GTTGGGAGTG
ahya_s0003.g68.t2  CAACGATACT CCCTCACAGT CTGTCAAGCG TCCTAAACCT GTTGGGAGTG
acyt_s1402.g1.t1   ---------- ---------- ---------- ---------- ----------
amur_s0006.g105.t2 CAACGATACT CCCTCACAGT CTGTCAAGCG CCCTAAACCT GTTGGGAGTG
ech_s0159.g30.t2   CAACGATACT CCCTCACAGT CTGTCAAGCG TCCTAAACCT GTTGGGAGTG
aacu_s0038.g69.t1  CAACGATACT CCCGCACAGT CTGCCAAGCG CCCTAAACCT GTTGGGAGTC
anas_s0109.g72.t2  CAATGATACA CCCTCACAGT CTGTCAAGCG CCCTAAACCT GTTGGGAGTG
amic_s0245.g8.t1   CAACGATACT CCCTTACAGT CTGTCAAGCG CCCTAAACCT GTTGGGAGTG
adig_s0048.g28.t1  CAACGATACT CCCTCACAGT CTGTCAAGCG CCCTAAACCT GTTGGGAGTG
Asp1_sequenced_cds CAATGATACT CCCTCACAG- ---------- ---------- ----------

aten_s0183.g20.t1   TCACCGATAC TATGGTTACT GAATCATCAC CA------AG TGAGTGG---
ayon_s0004.g209.t1  TCACCGATAC TATGGTTACT GAATCATCAC CA------AG TGAGTGG---
aint_s0143.g6.t1    TTCCCGATCC TATGGCTACT GAATCATCAC CA------AG TGAGTGG---
agem_s0013.g175.t1  TTCCCGATCC TATGGCTACT GAATCATCAC CA------AG TGAGTGG---
aawi_s0007.g175.t1  TTCCCGATCC TATGGCTACT GAATCATCAC CA------AG TGAGTGG---
aflo_s0310.g20.t1   TTCCCGATCC TATGGCTACT GAATCATCAC CA------AG TGAGTGG---
XM_029340177.2      TTTCCGATCC TATGGTTAAT GAATCATCAC CA------AG TGAGTGG---
asel_s0045.g58.t2   TTCCCGATCC TATGGGTAAT GAATCATCAC CA------AG TGAGTGG---
ahya_s0003.g68.t2   TTCCCGATCC TATGGGTAAT GAATCATCAC CA------AG TGAGTGG---
acyt_s1402.g1.t1    ---------- ---------- ---------- ---------- ----------
amur_s0006.g105.t2  TTCCCGATCC TATGGTTACT GAATCATCAT CA------AG TGAGTGG---
ech_s0159.g30.t2    TTCCCGATCC TATGGGTAAT GAATCATCAC CA------AG TGAGTGG---
aacu_s0038.g69.t1   TTCCCGATCC TATGGTTACT GAATCATCAC CAATAAGGAT CCAATGGAAG
anas_s0109.g72.t2   TTCTCGTTCC TATGGTTACT GAATCATCAC CA------AG TGAGTGG---
amic_s0245.g8.t1    TTCCCGATCC TACGGGTAAT GAATCATCAC CA------AG TGAGTGG---
adig_s0048.g28.t1   TTCCCGATCC TATGGTTACT GAATCATCAC CA------AG TGAGTGG---
Asp1_sequenced_cds  ---------- ---------- ---------- ---------- ----------

aten_s0183.g20.t1  ---AGTGAAG TGTCGTTTCC GCTGAGTGGC GGATTGTCTC CACCAGTTTT
ayon_s0004.g209.t1 ---AGTGAAG TGTCGTTTCC GCTGAGTGGC GGATTGTCTC CACCAGTTTT
aint_s0143.g6.t1   ---AGTGAAG TGTCGTTTCC GCTGAGTGGC GGATTGTCTC CAACAGTTTT
agem_s0013.g175.t1 ---AGTGAAG TGTCGTTTCC GCTGAGTGGC GGATTGTCTC CAACCGTTTT
aawi_s0007.g175.t1 ---AGTGAAG TGTCGTTTCC GCTGAGTGGC GGATTGTCTC CAACCGTTTT
aflo_s0310.g20.t1  ---AGTGAAG TGTCGTTTCC GCTGAGTGGC GGATTGTCTC CAACCGTTTT
XM_029340177.2     ---AGTGAAG TGACGTTTCC GCTGAGTGGC GGATTGTCTC CAACAGTTTT
asel_s0045.g58.t2  ---AGTGAAG TGACGTTTCC GCTGAGTGGC GGATTGTCTC CAACAGTTTT
ahya_s0003.g68.t2  ---AGTGAAG TGTCGTTTCC GCTGAGTGGC GGATTGTCTC CAACAGTTTT
acyt_s1402.g1.t1   ---------- ---------- ---------- ---------- ----------
amur_s0006.g105.t2 ---AGTGAAG TGTCGTTTCC GCTGAGTGGC GGATTGTCTC CAACAGTTTT
ech_s0159.g30.t2   ---AGTGAAG TGTCGTTTCC GCTGAGTGGC GGATTGTCTC CAACAGTTTT
aacu_s0038.g69.t1  AACAAAGAAA GCAGTTTCAA AGTGACTGCA GATTTGTCGA CCCCT-----
anas_s0109.g72.t2  ---AGTGAAG TGTCGTTTCC GCTGAGTGGC GGACTGTCTC CAACAGTTTC
amic_s0245.g8.t1   ---AGTGAAT TGTCGTTTCC GCTGAGTGGC GGATTGTCTC CAACAGTTTT
adig_s0048.g28.t1  ---AGTGAAT TGTCGTTTCC GCTGAGTGGC GGATTGTCTC CAACAGTTTT
Asp1_sequenced_cds ---------- ---------- ---------- ---------- ----------
```

### Fig S4 continued

```
aten_s0183.g20.t1  TATTCGAAGT AAGGATCCAA AGGAAGAGCA AAGAAAACAG TATCAAAGTG
ayon_s0004.g209.t1 TATTCGAAGT AAGGATCCAA AGGAAGAGCA AAGAAAACAG TATCAAAGTG
aint_s0143.g6.t1   TATTCGAAGT AAGGATCCAA TGGAAGAACA AAGAAAGCAA TATCAAAGTG
agem_s0013.g175.t1 TATTCGAAGT AAGGATCCAA TGGAAGAACA AAGAAAGCAG TATCAAAGTG
aawi_s0007.g175.t1 TATTCGAAGT AAGGATCCAA TGGAAGAACA AAGAAAGCAG TATCAAAGTG
aflo_s0310.g20.t1  TATTCGAAGT AAGGATCCAA TGGAAGAACA AAGAAAGCAG TATCAAAGTG
XM_029340177.2     TATTCGAAGT AAGGATCCAA TGGAAGAACA AAGAAAGCAG TTTCAAAGTG
asel_s0045.g58.t2  TATTCGAAGT AAGGATCCAA TGGAAGAACA AAGAAAGCAG TTTCAAAGTG
ahya_s0003.g68.t2  TATTCGAAGT AAGGATCCAA TGGAAGAACA AAGAAAGCAG TTTGAAAGTG
acyt_s1402.g1.t1   ---------- ---------- ---------- ---------- ----------
amur_s0006.g105.t2 TATTCGAAGT AAGGATCCAA TGGAAGAACA AAGAAAGCAG TTTCAAAGTG
ech_s0159.g30.t2   TATTCGAAGT AAGGATCCAA TGGAAGAACA AAGAAAGCAG TTTCAAAGTG
aacu_s0038.g69.t1  ---------- ---------- ---------- ---------- CGCAAGTCAA
anas_s0109.g72.t2  TATTCGAAGT AAGGATCCAA TGGAAGAACA AAAAAAGCAG TTTAAAAGTG
amic_s0245.g8.t1   TATTCGAAGT AAGGATCCAA TGGAAGAACA AAAGAAGCAG TTTAAAAGTG
adig_s0048.g28.t1  TATTCGAAGT AAGGATCCAA TGGAAGAACA AAGAAAGCAG TTTAAAAGTG
Asp1_sequenced_cds ---------- ---------- ---------- ---------- ----------

aten_s0183.g20.t1  ACTGCAGATT GGTCGACCCT TCGCAGGTCA AACTGCATGA CCATTACATA
ayon_s0004.g209.t1 ACTGCAGATT GGTCGACCCT TCGCAGGTCA AACTGCATGA CCATTACATA
aint_s0143.g6.t1   ACTGCAGATT TGTCGACCCT TCGCAGGTTA AACTGCATGA CCATTACATA
agem_s0013.g175.t1 ACTGCAGATT TGTCGACCCT TCGCAGGTTA AACTGCATGA CCATTATATA
aawi_s0007.g175.t1 ACTGCAGATT TGTCGACCCT TCGCAGGTTA AACTGCATGA CCATTACATA
aflo_s0310.g20.t1  ACTGCAGATT TGTCGACCCT TCGCAGGTTA AACTGCATGA CCATTACATA
XM_029340177.2     ACTGCAGATT TGTCGACCCT TCGCAGGTTA AACTGCATGA CCATTACATA
asel_s0045.g58.t2  ACTGCAGGTG A--------- ---------- ---------- ----------
ahya_s0003.g68.t2  ACTGCAGGTG A--------- ---------- ---------- ----------
acyt_s1402.g1.t1   ---------- ---------- ---------- ---------- ----------
amur_s0006.g105.t2 ACTGCAGGTG A--------- ---------- ---------- ----------
ech_s0159.g30.t2   ACTGCAGGTG A--------- ---------- ---------- ----------
aacu_s0038.g69.t1  ACTGCATGAC CATTACATAA ---------- ---------- ----------
anas_s0109.g72.t2  ACTGCAGGTG A--------- ---------- ---------- ----------
amic_s0245.g8.t1   ACTCCAGGTG A--------- ---------- ---------- ----------
adig_s0048.g28.t1  ACTGCAGATT TGTCGACCCT TCGCAGGTTA AACTGCATGA CCATTACATA
Asp1_sequenced_cds ---------- ---------- ---------- ---------- ----------

aten_s0183.g20.t1  AAGCAGTACG CTGATGTTCT GTACCGATGG AGTTTGCTGG GCAAAAGAGC
ayon_s0004.g209.t1 AAGCAGTACG CTGATGTTCT GTACCGATGG AGTTTGCTGG GCAAAAGAGC
aint_s0143.g6.t1   AAACAGTACG CTGATGTTCT GTACCGATGG AGTTTGCTAG GCAAAAGAGC
agem_s0013.g175.t1 AAACAGTACG CTGATGTTCT GTACCGATGG AGTTTGCTGG GCAAAAGAGC
aawi_s0007.g175.t1 AAACAGTACG CTGATGTTCT GTACCGATGG AGTTTGCTGG GCAAAAGAGC
aflo_s0310.g20.t1  AGACAGTACG CAGATGTTCT GTACCGATGG AGTTTGCTGG GCAAAAGAGC
XM_029340177.2     AAACAGTACG CTGATGTTCT GTACCGATGG AGTTTGCTGG GCAAAAGAGC
asel_s0045.g58.t2  ---------- ---------- ---------- ---------- ----------
ahya_s0003.g68.t2  ---------- ---------- ---------- ---------- ----------
acyt_s1402.g1.t1   ---------- ---------- ---------- ---------- ----------
amur_s0006.g105.t2 ---------- ---------- ---------- ---------- ----------
ech_s0159.g30.t2   ---------- ---------- ---------- ---------- ----------
aacu_s0038.g69.t1  ---------- ---------- ---------- ---------- ----------
anas_s0109.g72.t2  ---------- ---------- ---------- ---------- ----------
amic_s0245.g8.t1   ---------- ---------- ---------- ---------- ----------
adig_s0048.g28.t1  AAACAGTACG CTGATGTTCT GTACCGATGG AGTTTGCTGG GCAAAAGAGC
Asp1_sequenced_cds ---------- ---------- ---------- ---------- ----------

aten_s0183.g20.t1  TGAAGTTACC AAATTCTTGA GTGAAACGCA AGTGCCTCAC AGTGGCGCAG
ayon_s0004.g209.t1 TGAAGTTACC AAATTCTTGA GTGAAACGCA ATTGCCTCAC AGTGGCGCAG
aint_s0143.g6.t1   TGAAGTTACC AAATTCTTGA GTCAACCGCA ATTGCCTCAC AGTGGCGCAG
agem_s0013.g175.t1 TGAAGTTACC AAATTCTTGA GTGAACCGCA ATTGCCTCAC AGTGGCGCAG
aawi_s0007.g175.t1 TGAAGTTACC AAATTCTTGA GTGAACCGCA ATTGCCTCAC AGTGGCGCAG
aflo_s0310.g20.t1  TGAAGTTACC AAATTCTTGA GTGAACCGCA ATTGCCTCAC AGTGGCGCAG
XM_029340177.2     TGAAGTTACC AAATTCTTAA GTGAACCGCA ATTGCCTCAC AGTGGCGCAG
asel_s0045.g58.t2  ---------- ---------- ---------- ---------- ----------
ahya_s0003.g68.t2  ---------- ---------- ---------- ---------- ----------
acyt_s1402.g1.t1   ---------- ---------- ---------- ---------- ----------
amur_s0006.g105.t2 ---------- ---------- ---------- ---------- ----------
ech_s0159.g30.t2   ---------- ---------- ---------- ---------- ----------
aacu_s0038.g69.t1  ---------- ---------- ---------- ---------- ----------
anas_s0109.g72.t2  ---------- ---------- ---------- ---------- ----------
amic_s0245.g8.t1   ---------- ---------- ---------- ---------- ----------
adig_s0048.g28.t1  TGAAGTTACC AAATTCTTGA GTGAACCGCA ATTGCCTCAC AGTGGCGCAG
Asp1_sequenced_cds ---------- ---------- ---------- ---------- ----------
```

### Fig S4 continued

```
aten_s0183.g20.t1  AATTCAGTAC GCGTTGCTAC AATTGTTCCC GAAACCTTCG TGGTGCTCAG
ayon_s0004.g209.t1 AATTCAGTAC GCGTTGCTAC AATTGTTCCC GAAACTTTCG TGGTGCTCAG
aint_s0143.g6.t1   AATTCAGTAC GCGTTGCTAC AATTGTTCCC GAAACCTTCG TGGTGCTCAG
agem_s0013.g175.t1 AATTCAGTAC GCGTTGCTAC AATTGTTCCC GAAACCTTCG TGGTGCTCAG
aawi_s0007.g175.t1 AATTCAGTAC GCGTTGCTAC AATTGTTCCC GAAACCTTCG TGGTGCTCAG
aflo_s0310.g20.t1  AATTCAGTAC GCGTTGCTAC AATTGTTCCC GAAACCTTCG TGGTGCTCAG
XM_029340177.2     AATTCAGTAC GCGTTGCTAC AATTGTTCCC GAAACCTTCG TGGTGCTCAG
asel_s0045.g58.t2  ---------- ---------- ---------- ---------- ----------
ahya_s0003.g68.t2  ---------- ---------- ---------- ---------- ----------
acyt_s1402.g1.t1   ---------- ---------- ---------- ---------- ----------
amur_s0006.g105.t2 ---------- ---------- ---------- ---------- ----------
ech_s0159.g30.t2   ---------- ---------- ---------- ---------- ----------
aacu_s0038.g69.t1  ---------- ---------- ---------- ---------- ----------
anas_s0109.g72.t2  ---------- ---------- ---------- ---------- ----------
amic_s0245.g8.t1   ---------- ---------- ---------- ---------- ----------
adig_s0048.g28.t1  AATTCAGTAC GCGTTGCTAC AATTGTTCCC GAAACCTTCG TGGTGCTCAG
Asp1_sequenced_cds ---------- ---------- ---------- ---------- ----------

aten_s0183.g20.t1  TGCGGTTCTT GTAAATCGTT TGGACTACAG TGTGTGATTT GTCATGTGGC
ayon_s0004.g209.t1 TGCGGTTCTT GTAAATCGTT TGGACTACAG TGTGTGATTT GTCATGTGGC
aint_s0143.g6.t1   TGTGGTTCTT GTAAATCGTT TGGACTACAG TGTGTGATAT GTCATGTGGC
agem_s0013.g175.t1 TGTGGTTCTT GTAAATCGTT TGGACTACAG TGTGTGATAT GTCATGTGGC
aawi_s0007.g175.t1 TGTGGTTCTT GTAAATCGTT TGGACTACAG TGTGTGATAT GTCATGTGGC
aflo_s0310.g20.t1  TGTGGTTCTT GTAAATCGTT TGGACTACAG TGTGTGATAT GTCATGTGGC
XM_029340177.2     TGTGGGTCTT GTAAATCGTT TGGACTACAG TGTGTGATAT GTCATGTGGC
asel_s0045.g58.t2  ---------- ---------- ---------- ---------- ----------
ahya_s0003.g68.t2  ---------- ---------- ---------- ---------- ----------
acyt_s1402.g1.t1   ---------- ---------- ---------- ---------- ----------
amur_s0006.g105.t2 ---------- ---------- ---------- ---------- ----------
ech_s0159.g30.t2   ---------- ---------- ---------- ---------- ----------
aacu_s0038.g69.t1  ---------- ---------- ---------- ---------- ----------
anas_s0109.g72.t2  ---------- ---------- ---------- ---------- ----------
amic_s0245.g8.t1   ---------- ---------- ---------- ---------- ----------
adig_s0048.g28.t1  TGTGGTTCTT GTAAATCGTT TGGACTACAG TGTGTGATAT GTCATGTGGC
Asp1_sequenced_cds ---------- ---------- ---------- ---------- ----------

aten_s0183.g20.t1  TGTAAGAGGT GCCAGTAATT TCTGTGTCGC CTGCGGTCAC GGCGGTCACG
ayon_s0004.g209.t1 TGTAAGAGGT GCCAGTAATT TCTGTGTCGC CTGCGGTCAC GGCGGTCACG
aint_s0143.g6.t1   TGTAAGAGGT GCCAGTAATT TCTGTGTCGC CTGCGGTCAC GGCGGTCACG
agem_s0013.g175.t1 TGTAAGAGGT GCCAGTAATT TCTGTGTCGC CTGCGGTCAC GGCGGTCACG
aawi_s0007.g175.t1 TGTAAGAGGT GCCAGTAATT TCTGTGTCGC CTGCGGTCAC GGCGGTCACG
aflo_s0310.g20.t1  TGTAAGAGGT GCCAGTAATT TCTGTGTCGC CTGCGGTCAC GGCGGTCACG
XM_029340177.2     TGTAAGAGGT GCCAGTAATT TCTGTGTCGC CTGCGGTCAC GGCGGTCACG
asel_s0045.g58.t2  ---------- ---------- ---------- ---------- ----------
ahya_s0003.g68.t2  ---------- ---------- ---------- ---------- ----------
acyt_s1402.g1.t1   ---------- ---------- ---------- ---------- ----------
amur_s0006.g105.t2 ---------- ---------- ---------- ---------- ----------
ech_s0159.g30.t2   ---------- ---------- ---------- ---------- ----------
aacu_s0038.g69.t1  ---------- ---------- ---------- ---------- ----------
anas_s0109.g72.t2  ---------- ---------- ---------- ---------- ----------
amic_s0245.g8.t1   ---------- ---------- ---------- ---------- ----------
adig_s0048.g28.t1  TGTAAGAGGT GCCAGTAATT TCTGTGTCGC CTGCGGTCAC GGCGGTCACG
Asp1_sequenced_cds ---------- ---------- ---------- ---------- ----------

aten_s0183.g20.t1  CGTATCATCT ACTGACGTGG TTTGAATCTA TGAATGTCTG CCCAACAGGC
ayon_s0004.g209.t1 CGTATCATCT ACTGACGTGG TTTGAATCTA TGAATGTCTG CCCAACAGGC
aint_s0143.g6.t1   CGTATCATCT ATTGACGTGG TTTGAATCTA TGGATGTCTG CCCAACAGGC
agem_s0013.g175.t1 CGTATCATCT ATTGACGTGG TTTGAATCTA TGGATGTCTG CCCAACAGGC
aawi_s0007.g175.t1 CGTATCATCT ATTGACGTGG TTTGAATCTA TGGATGTCTG CCCAACAGGC
aflo_s0310.g20.t1  CGTATCATCT ATTGACGTGG TTTGAATCTA TGGATGTCTG CCCAACAGGC
XM_029340177.2     CGTATCATCT ACTGACGTGG TTTGAATCTA TGGATGTCTG CCCAACAGGC
asel_s0045.g58.t2  ---------- ---------- ---------- ---------- ----------
ahya_s0003.g68.t2  ---------- ---------- ---------- ---------- ----------
acyt_s1402.g1.t1   ---------- ---------- ---------- ---------- ----------
amur_s0006.g105.t2 ---------- ---------- ---------- ---------- ----------
ech_s0159.g30.t2   ---------- ---------- ---------- ---------- ----------
aacu_s0038.g69.t1  ---------- ---------- ---------- ---------- ----------
anas_s0109.g72.t2  ---------- ---------- ---------- ---------- ----------
amic_s0245.g8.t1   ---------- ---------- ---------- ---------- ----------
adig_s0048.g28.t1  CGTATCATCT ATTGACGTGG TTTGAATCTA TGGATGTCTG CCCAACAGGC
Asp1_sequenced_cds ---------- ---------- ---------- ---------- ----------
```

### Fig S4 continued

```
aten_s0183.g20.t1  TGTGGATGTA GGTGTCTGGA AGTCGAT--- ACATTCATTG TGGATTGA
ayon_s0004.g209.t1 TGTGGATGTA GGTGTCTGGA AGTCGAT--- ACATTCATTG TGGATTGA
aint_s0143.g6.t1   TGTGGATGTA GGTGTTTGGA AGTCGGT--- ACATTCATTG TGGATTGA
agem_s0013.g175.t1 TGTGGATGTA GGTGTCTGGA AGTCGGT--- ACATTCATTG TAGATTGA
aawi_s0007.g175.t1 TGTGGATGTA GGTGTCTGGA AGTCGGT--- ACATTCATTG TGGATTGA
aflo_s0310.g20.t1  TGTGGATGTA GGTGTCTGGA AGTCGGT--- ACATTCATTG TGGATTGA
XM_029340177.2     TGTGGATGTA GGTGTCGGGA AGTCGGGGTA ACATTTATTG TGGATTGA
asel_s0045.g58.t2  ---------- ---------- ---------- ---------- --------
ahya_s0003.g68.t2  ---------- ---------- ---------- ---------- --------
acyt_s1402.g1.t1   ---------- ---------- ---------- ---------- --------
amur_s0006.g105.t2 ---------- ---------- ---------- ---------- --------
ech_s0159.g30.t2   ---------- ---------- ---------- ---------- --------
aacu_s0038.g69.t1  ---------- ---------- ---------- ---------- --------
anas_s0109.g72.t2  ---------- ---------- ---------- ---------- --------
amic_s0245.g8.t1   ---------- ---------- ---------- ---------- --------
adig_s0048.g28.t1  TGTGGATGTA GGTGTCGGGA AGTCGGT--- ACATTCACTG TGGATTGA
Asp1_sequenced_cds ---------- ---------- ---------- ---------- --------
```

**Figure S5.** Alignment of *WDR59* sequences from 16 *Acropora* corals and the *Acropora* sp.1 *WDR59* sequence determined by PCR and sequencing.

### Fig S6

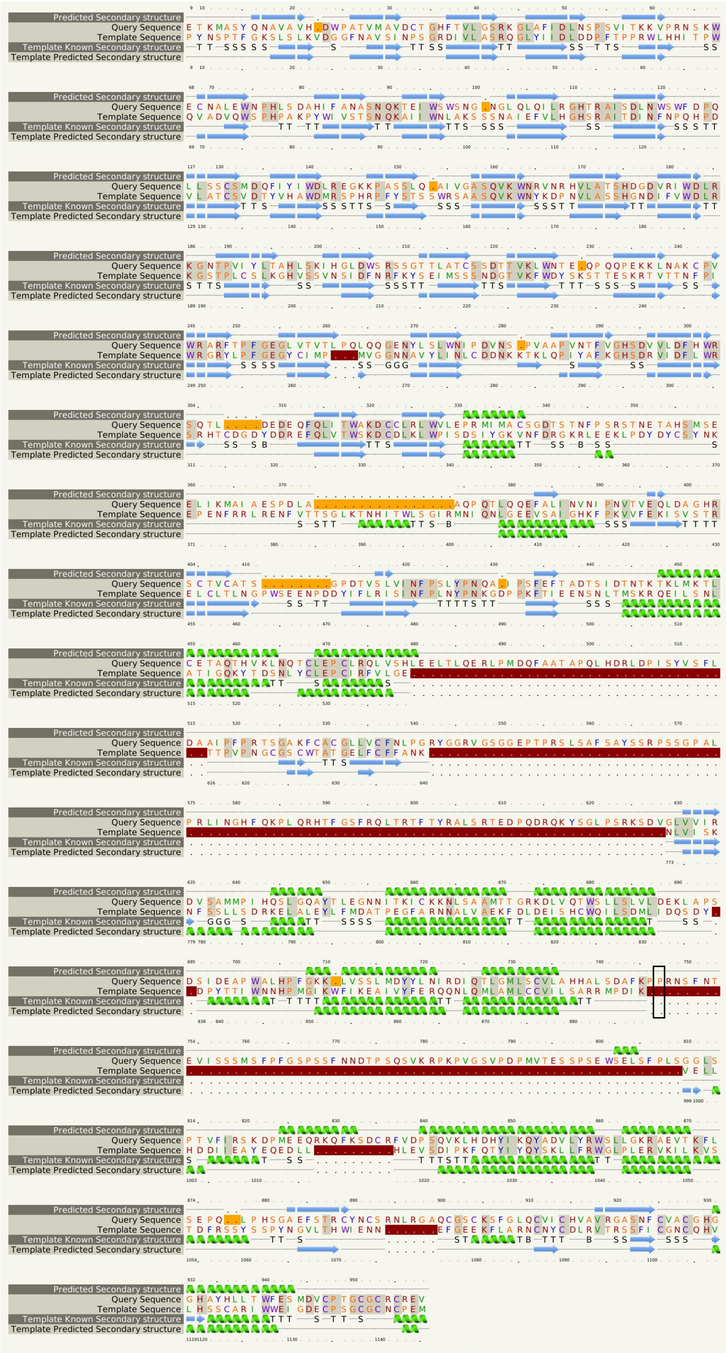

Predicted Secondary structure
Query Sequence
Template Sequence
Template Known Secondary structure
Template Predicted Secondary structure

**Figure S6.** Alignment of *A. digitifera WDR59* (Gene ID: adig_s0048_g28) as a query and template sequence (c8adlQ) using phyre2. Insertions relative to the template are shown in red. Deletions relative to the template are shown in orange. A black line surrounds the position of the *Acropora* sp1-specific mutation.

**3 Supplemental Tables**

Table S1. Sample information used in this study.

Table S2. The genomic location of HDRs in the *A. digitifera* genome assembly ver 2.0.

Table S3. The result of a Blastn search against the NCBI database using 39 genes as queries.

Table S4. Non-synonymous differentiated SNPs in candidate genes.

Table S4. Orthologous genes of WDR59 in 15 Acropora species. * identified by KEGG information.
